# Dispersal Behaviour and Movement Patterns of Pheasants from Woodland Release Pens

**DOI:** 10.64898/2026.02.11.705276

**Authors:** Jennifer L. Page, Daniel A. Warren, Julia Coats, Izzy Rochester, Kate L. Palphramand, Dave Parrott

## Abstract

The large-scale release of ring-necked pheasants, *Phasianus colchicus*, for recreational shooting in the UK raises concerns about ecological impacts, particularly on sensitive ecological sites. Current assumptions suggest dispersal is typically <500m from release pens, yet empirical evidence is limited. This study tracked 110 GPS-tagged pheasants from 11 woodland release pens across nine shooting estates, monitoring movements through pre-shooting, shooting and post-shooting phases. Most birds (73%) travelled a maximum distance beyond 500m during at least one of the three phases, with mean maximum distances of 863m, 1,493m and 1,307m per phase. During at least one phase, 26% of the 110 tagged birds spent most of their time (>50%) beyond 500m and 16% beyond 1,000m from their release pens. Early post-release movements were concentrated near pens, but ranging behaviour expanded during subsequent phases, with the percentages of birds spending >50% of their time beyond 500m and 1,000m, respectively: pre-shooting 6%, 2%; shooting 24%, 16%; post-shooting 13%, 9%. Accounting for mortality, the percentages of surviving birds spending >50% of their time beyond 500m and 1,000m increased: pre-shooting (n=110) 6%, 2%; shooting (n=71) 37%, 25%; post-shooting (n=27) 52%, 37%. Dispersal was greater with earlier release dates, higher pen and estate stocking densities and lower vegetative habitat quality in pens. Movements were directional rather than uniform, with most cohorts concentrating activity within a limited directional arc specific to the release site. Conservation site incursions occurred in 28 (25%) tagged birds, particularly where pens were closest to site boundaries; although 10 (36%) tagged birds encroached on conservation sites 872-2,319m from their release pen. These findings show that dispersal of released pheasants is further, more directed, and persistent than currently assumed.

## Introduction

The artificial rearing and release of gamebirds, particularly the non-native ring-necked pheasant, *Phasianus colchicus*, and red-legged partridge, *Alectoris rufa*, to supplement natural populations for recreational shooting, is commonplace across the UK (Madden, 2021). The number of gamebirds released annually has increased over the years. Recent estimate suggests that 31.5 million pheasants (range 29.8 – 33.7 million), 9.1 million red-legged partridges (range 5.6 – 12.5 million) and 2.6 million mallards (range 0.9 – 6.0 million) are released annually in the UK (Madden, 2021), with the combined biomass of released and naturalised gamebirds estimated to be double that of all UK native breeding birds combined (Mason et al., 2020).

The widespread mass release of gamebirds and their management is associated with positive and negative effects on habitats and wildlife. High densities may lead to several effects, including habitat alteration (through herbivory and eutrophication), predation of invertebrates and small vertebrates, inter-species competition, and the spread of disease (Madden & Sage, 2020; Mason et al., 2020; Sage et al., 2020). Concerns have been raised over the release of gamebirds and their potential negative ecological effects, and their potential impact on European Protected Sites, which are designated under two main directives: the Birds Directive (for Special Protection Areas, SPA) and the Habitats Directive (for Special Areas of Conservation, SAC). These sites incorporate Sites of Special Scientific Interest (SSSI), which are themselves covered by consenting processes to protect them from detrimental impacts.

Assessing the movements of pheasants in the landscape following their release will help identify the risk of potential ecological impacts to these sensitive sites. Current assumptions about gamebird dispersal are reflected in The Defra Gamebird Review (GOV.UK, 2020) conclusion that ‘…*dispersal of birds tends to be less than 500 m from the release sites*…’. Current licensing regulations are predicated on this assumption.

From previous research, pheasants are thought to be reasonably sedentary, with past studies indicating that wild birds tend to stay within 5km of their hatching place (Gatti et al., 1989; Krauss et al., 1987; Leif, 2005; Ridley, 1983; Wilson et al., 1992). Several European radio-tracking studies on both wild birds (Bagliacca et al., 2008) and reared and released birds (Alonso et al., 2005; Bagliacca et al., 2008; Duarte et al., 2011; Gruychev, 2014; Pérez et al., 2004) have reported mean dispersal distances ranging from 260m to 1,209m for pheasants and 495m to 833m for red-legged partridge. Home ranges of wild pheasants vary widely in the literature and are often influenced by habitat, seasonality and sex (Gatti et al., 1989; Leif, 2005; Smith et al., 1999). In studies based outside of the UK, however, landscape, habitat and game management differ from those of UK shooting estates. Additionally, many studies focus on dispersal distances of caught and released wild birds, which may behave differently from captive reared birds raised for UK shoots, or track birds at different times of the year other than late summer when birds are released in the UK. For this reason, studies are often not comparable with pheasant release and dispersal in the UK.

When considering dispersal patterns of pheasant and red-legged partridge in the UK, studies are limited, but provide evidence that female dispersal is greater than male dispersal (Beardsworth et al., 2021; Hill & Ridley, 1987; Turner, 2008), wild bred birds travel greater distances than game farm bred birds (Sage et al., 2001) and movement of birds can be influenced by game management practice, habitat quality and season (Sage et al., 2001; Turner, 2008).

However, most gamebird tracking studies have used VHF radio telemetry and triangulation to locate pheasants and track their movements, which comes with limitations: (i) researchers are required to be close to their target species to accurately triangulate their position; (ii) the topography of a landscape can make triangulation difficult and can lead to errors in identifying a target animal’s location; (iii) data collected from radiotelemetry studies can be sparse due to researchers having to be present in the field to record an animal’s location, in some cases, as few as two fixes per week (e.g. Turner, 2008). A few studies have used the reverse GPS ATLAS system (Beardsworth et al., 2021) in which tags, fitted to birds, emit signals to multiple ground-based receivers distributed throughout the study area, thus automatically triangulating locations. The ATLAS system, however, requires line of sight from three or more receivers, which is facilitated more by open habitats rather than the more occluded woodland edge habitats favoured by pheasants.

There are, however, very few empirical studies on the dispersal of reared and released pheasants that have employed the advances in GPS tracker technology. The current study investigated pheasant dispersal behaviour and movement patterns from woodland release pens on working shoots, across a range of management and landscape types that may influence variation in dispersal between different release pens and/or shoot sites. Specifically, GPS tracking technology was used to monitor: (i) dispersal distances, (ii) relative time spent at varying distances from the release pen, (iii) directional preference of movements in relation to the release pen, and (iv) rate and nature of incursions onto neighbouring Conservation Sites, across three sequential phases of the ‘gamebird-year’ (pre-shooting, shooting, and post-shooting season). The advancement of telemetry using GPS and the ability to continuously locate an animal with precision and accuracy over 24 hours provides a research advantage and increases the temporal resolution of collected data. In addition, the study evaluated the influence of several key management parameters on dispersal distance: (i) release date, (ii) pen stocking density, (iii) total estate stocking density, and (iv) quality of release pen vegetative cover.

## Materials and Methods

### Study sites

Nine estates/working shoots (sites A-I) were selected for the study located in a range of landscape types as defined under Natural England’s National Character Area Profiles <u>National Character Area profiles - GOV.UK</u> (www.gov.uk), encompassing lowland, upland and transition landscapes. At two estates (C and F), two different release pens were monitored on different parts of the estate.

All 11 release pens were located within 4.5 km (range = 149 – 4,305m) of at least one conservation site, designated as either a Site of Special Scientific Interest (SSSI), Special Area of Conservation (SAC), or Special Protection Area (SPA). Most release pens (8/11) were situated within 1km (range = 149 – 959m) of such a designated conservation site.

### GPS tagging and programming

Across all 11 release pens, 110 ring-necked pheasant poults (median 10 per pen, aged 7-11 weeks) were fitted with PinPoint VHF GPS transmitters (Lotek), attached via Perlon® necklaces (Table 1). Tags were fitted on birds as they were transferred between delivery crates and release pens. Birds too light to achieve a suitable tag to body weight ratio (tags should not exceed 2%-5% of body weight, Kenward, 2001) were excluded. Young pheasants continue to grow quickly meaning that tags would be expected to represent 0.8% to 2.7% of adult body weight. Tags recorded locations every 60 minutes (up to 24 fixes per day) and were programmed using PinPoint Host software under a SWIFT fix system (satellite search time of 12 to 40 seconds compared to a standard fix system of 35 to 70 seconds) to reduce energy use and extend battery life. Each tag transmitted GPS data, a Beacon signal for locating birds (using a Biotracker receiver and Yagi antenna) and VHF communication for data download (via a PinPoint Commander (PPC) and Yagi antenna). Beacon and VHF schedules were initially set for daily two-hour windows for welfare checks (first week only), then adjusted to weekly four-hour windows to maximise battery life. Tags included a mortality feature, indicated by a change in Beacon pulse frequency, after 10 hours of inactivity, indicating either a shed tag or the death of the pheasant. On average, data was collected once a fortnight at each site.

**Table 1.** Summary of GPS tagging details, including release date, total birds (males and females), and ranges for bird weight, tag weight, and tag-to-body weight ratio.

| Site | Date | No. tags | No. males | No. females | Mean bird wt. (g) | Mean tag wt. (g) | Mean % body wt. |
| --- | --- | --- | --- | --- | --- | --- | --- |
| A | 16/07/23 | 10 | 5 | 5 | 591 | 18.3 | 3.1 |
| B | 17/07/23 | 10 | 5 | 5 | 515 | 17.9 | 3.5 |
| C1 | 19/07/23 | 10 | 5 | 5 | 616 | 18.5 | 3.0 |
| C2<sup>1</sup> | 21/07/23 | 13 | 6 | 7 | 625 | 18.1 | 2.9 |
| D | 25/07/23 | 10 | 5 | 5 | 571 | 17.3 | 3.1 |
| E | 31/07/23 | 10 | 5 | 5 | 591 | 17.5 | 3.0 |
| F1 | 03/08/23 | 10 | 5 | 5 | 541 | 18.2 | 3.4 |
| F2 | 03/08/23 | 10 | 5 | 5 | 543 | 18.1 | 3.3 |
| G | 11/08/23 | 12 | 6 | 6 | 572 | 16.8 | 3.0 |
| H | 22/08/23 | 11 | 6 | 5 | 734 | 17.9 | 2.5 |
| I<sup>2</sup> | 05/09/23 | 6 | 4 | 2 | 660 | 17.6 | 2.8 |
| **Total/Range** |  | 112 | 57 | 55 | 515-734 | 16.8-18.5 | 2.5-3.5 |

<sup>1</sup> Pen C2 received 3 extra birds four weeks after initial 10 birds tagged. Of these three birds, one bird died within a few days, one tag immediately failed, and one bird was predated during the pre-shooting season.
<sup>2</sup> Pen I received tags already used on previous birds that were found dead and reapplied to birds in this pen.

### Management practices

Where possible, information regarding release dates, stocking density (per pen and estate-wide), feeding method and feeder location(s), the total number of shoots and date of first shoot, and the distribution of cover/game crops was obtained through consultation with estate managers and gamekeepers (Table 2). However, the amount and type of information provided was inconsistent across shooting estates: where information was consistent across all estates, it was incorporated into analyses, when possible.

**Table 2.** Summary of pheasant release parameters for each release estate/pen.

| Pen | Date | Age (weeks) | No. Birds pen | Density pen (birds/ha) | Area of estate (ha) <sup>1</sup> | Tot. no. pheasants on estate | Density estate (birds/ha) | First shoot | Pen habitat quality <sup>2</sup> |
| --- | --- | --- | --- | --- | --- | --- | --- | --- | --- |
| A | 16.07.23 | 8 | 300 | 3,614 | 146 | 600 | 4.1 | 14th Oct | 6.8 |
| B | 17.07.23 | 7/8 | 2,000 | 2,208 | 3,237 | 8,000 | 2.5 | 17th Oct | 5.0 |
| C1 | 19.07.23 | 7/8 | 1,000 | 347 | 1,619 | 4,000 | 2.5 | 4th Nov | 11.0 |
| C2 | 21.07.23 | 7/8 | 1,200 | 1,728 | 1,619 | 4,000 | 2.5 | 4th Nov | 11.0 |
| D | 25.07.23 | 7 | 2,500 | 1,870 | 2,630 | 14,000 | 5.3 | 21st Oct | 10.0 |
| E | 31.07.23 | 8 | 1,500 | 2,142 | 567 | 2,250 | 4.0 | 16th Sept | 5.3 |
| F1 | 03.08.23 | 8 | 500 | 883 | 3,237 | 6,000 | 1.9 | 10th Nov | 9.5 |
| F2 | 03.08.23 | 8 | 500 | 513 | 3,237 | 6,000 | 1.9 | 10th Nov | 9.5 |
| G | 11.08.23 | 7/8 | 880 | 1,212 | 809 | 3,800 | 4.8 | 17th Oct | 10.3 |
| H | 22.08.23 | 8/10 | 500 | 594 | 1,011 | 700 | 0.7 | 21st Oct | 6.0 |
| I | 05.09.23 | 11 | 300 | 557 | 1,618 | 600 | 0.4 | 9th Nov | 7.3 |

<sup>1</sup> Area of estate shot over
<sup>2</sup>Based on pheasant release pen scoring system checklist devised by BASC Pheasant release pen scoring system checklist - BASC. Though unable to assess all aspects of pen quality (e.g., feeders and drinkers, water, food, poults at release, predator vermin control) each pen was individually assessed by three different scorers on aspects of vegetative habitat quality (roosting area, area of cover, open area); the possible score ranged from a minimum of 0 (low quality) to a maximum of 11 (high quality). Average values between scorers are presented in the above table.

### Tracking period (‘gamebird year’)

Once tagged, pheasant poults were placed in woodland release pens (Table 2) and monitored throughout the ‘gamebird year’, encompassing the pre-shooting, shooting and post-shooting seasons. Each of these three phases impose different ‘push’ and ‘pull’ stimuli in respect to pheasant movements relative to the release pen. During the pre-shooting phase (July-September 2023), poults were initially fed within or near the pen before gradually being provisioned via feeders at increasing distances into areas scheduled for shoots. The pre-shooting phase, defined as the period from release to the day before the estate’s first shoot, varied between estates (46-107 days). By the start of the shooting phase (the period between the first estate shoot date and official end date of the pheasant shooting season), pheasants were fully mature and ranging at varying distances from the pen. Shooting phase duration also varied (83-138 days), followed by a post-shooting phase extending from the official end date of the shooting season to the date of final data collection at each site. Details of GPS data download, post-processing and validation procedures are provided in Supplementary Materials (S1, Section 1).

### Statistical Analysis

#### Dispersal distance

Analyses were performed using the R Statistical Programme (version 4.2.2; R Core Team) and the RStudio Integrated Development Environment (IDE; version 2022.12.0; Posit Software, PBC). A statistically significant threshold of 95% (i.e. *p* = 0.05) was applied across all analyses.

Analyses of dispersal behaviours were conducted using generalised linear mixed effect models (GLMMs), using the glmmTMB package (version 1.1.10; Brooks et al., 2017). For all GLMMs, the identification of individual GPS-tagged birds and release sites were set as nested random effects, thereby controlling for possible random variations in individual behaviours, as well as entire release cohorts.

The dispersal distance of individual pheasants, relative to the boundary of the release pen, was analysed using a zero-inflated GLMM. This was selected given the high number of GPS fixes that occurred within the limits of a release pen, representing 27.4% of all fixes. Fixes that occurred within a release pen were assigned a distance of zero metres (i.e. 0m). Given the existence of true zeroes (i.e. birds were recorded within the release pen), a zero-inflated gamma distribution was fitted to account for the inherent inflated right-skewed distribution that existed in the data. Study phase, release site ID, and sex were selected as main categorical predictor variables, with a two-way interaction term between phase and site (phase x site) included to identify if dispersal varied between certain phases and sites.

Further analysis was performed, to identify the effect that site-specific management might have on dispersal, in particular: a) the timing of pheasant release (day of year), b) the density at which pheasants are stocked in the pen (individuals ha^−1^), c) the density at which pheasants are stocked across the overall estate (individuals ha-1), and d) the quality of the vegetative cover in the release pen. For each comparison, a separate zero-inflated GLMM was fitted with a zero-inflated gamma distribution and a log-link function. For each model, dispersal distance was set as the response variable, and either release date, pen density, estate density, or pen quality, set as the predictor variable, respectively.

#### Ranging profile

The ranging profiles of individual birds, represented as the number of GPS fixes that occurred < 500m, 500 – 1,000m, or >1,000m from the pen were analysed using a GLMM fitted with a negative binomial error structure and a log-link function. This was selected to account for issues with overdispersion in model residuals (a common consequence of count data). Study phase, site, sex, and distance bin (i.e. <500m, 500 – 1,000m, >1,000m from the pen) were specified as main effects, along with a two-way interaction between study phase x distance bin ID, and between site x distance bin ID.

#### Directionality

As with ranging profiles, directionality was assessed using the number of GPS fixes that occurred within a set of octants, describing 45-degree (°) intervals around a release pen (i.e. 0 – 45°, 45 - 90°, 90 - 135°, 135 - 180°, 180 - 225°, 225 – 270°, 270 - 315°, 315 - 360°). Directional preference was defined as the average directionality exhibited by all GPS-tagged birds across these octants, indicating whether birds within a release cohort showed a consistent preferred direction of movement. For each octant, maximum distance refers to the average maximum distance travelled by all birds, while mean distance represents the average distance per bird, averaged across the cohort. Directionality was analysed using a GLMM fitted using a negative binomial error structure with log-link function. Study phase, site, sex and directional bin ID were included as main effects, with two-way interactions between study phase x directional bin ID and site x directional bin ID.

#### Conservation Site incursions

Analysis of conservation site incursions was applied to those sites where at least one incursion was recorded (i.e. 7/11 sites), whether it be an SSSI, SPA, or SAC. A logistic GLMM was fitted, with a binomial error structure and a logit-link function. Non-/incursions onto a conservation site were coded as a binary outcome, with 0 = non-incursion and 1 = incursion. Study phase, site and sex were included as categorical predictors, with dispersal distance (relative to release pens) included as a covariate predictor, scaled and centred based on the mean (centre) and standard deviation (scale). A two-way interaction was included between site and the scaled/centred dispersal distance.

## Results

### Tag performance and bird survival

Tags continued to return data on pheasant movements until one of four events: tag shedding, tag damage (signal loss), bird mortality, or battery depletion (Table 3). Of 110 tagged birds producing any data, 71 (65%) were confirmed alive at the start of shooting, 27 (25%) at the start of post-shooting, and 13 (12%) at study end, although figures represent minimum estimates due to tag failures, where the fate of the bird was unknown. Survival varied between release pens: pre-shooting 17-92%, shooting 9-50%, post-shooting 0-20%.

**Table 3.** Summary of mortality rates across the gamebird-year for each estate/pen. Values during each phase are cumulative from the date of tagging.

| Tagging |  |  | Pre-shooting |  |  |  |  | Shooting |  |  |  |  | Post-shooting |  |  |  |  |
| --- | --- | --- | --- | --- | --- | --- | --- | --- | --- | --- | --- | --- | --- | --- | --- | --- | --- |
| Site | Date tagged | No. Tagged<sup>1</sup> | Bird Died<sup>2</sup> | Tag Shed | Tag Failed<sup>3</sup> | No. Survived | % Survived | Bird Died<sup>2</sup> | Tag Shed | Tag Failed<sup>3</sup> | No. Survived | % Survived | Bird Died<sup>2</sup> | Tag Shed | Tag Failed<sup>3</sup> | No. Survived | % Survived |
| A | 16/07/23 | 10 | 1 | 0 | 1 | 8 | 80 | 4 | 0 | 3 | 3 | 30 | 4 | 0 | 5 | 1 | 10 |
| B | 17/07/23 | 10 | 1 | 0 | 0 | 9 | 90 | 3 | 0 | 2 | 5 | 50 | 3 | 0 | 6 | 1 | 10 |
| C1 | 19/07/23 | 10 | 0 | 0 | 6 | 4 | 40 | 1 | 0 | 7 | 2 | 20 | 1 | 0 | 8 | 1 | 10 |
| C2 | 21/07/23 | 11 | 4 | 1 | 2 | 6 | 55 | 9 | 1 | 2 | 1 | 9 | 9 | 1 | 2 | 1 | 9 |
| D | 25/07/23 | 10 | 1 | 0 | 2 | 7 | 70 | 3 | 0 | 4 | 3 | 30 | 3 | 0 | 5 | 2 | 20 |
| E | 31/07/23 | 10 | 1 | 0 | 1 | 8 | 80 | 3 | 0 | 6 | 1 | 10 | 3 | 0 | 6 | 1 | 10 |
| F1 | 03/08/23 | 10 | 2 | 0 | 3 | 5 | 50 | 5 | 0 | 4 | 1 | 10 | 5 | 0 | 4 | 1 | 10 |
| F2 | 03/08/23 | 10 | 1 | 0 | 5 | 4 | 40 | 3 | 0 | 6 | 1 | 10 | 3 | 0 | 7 | 0 | 0 |
| G | 11/08/23 | 12 | 0 | 0 | 1 | 11 | 92 | 2 | 0 | 4 | 6 | 50 | 2 | 0 | 8 | 2 | 17 |
| H | 22/08/23 | 11 | 1 | 0 | 2 | 8 | 73 | 6 | 0 | 2 | 3 | 27 | 6 | 0 | 3 | 2 | 18 |
| I | 05/09/23 | 6 | 2 | 0 | 3 | 1 | 17 | 2 | 0 | 3 | 1 | 17 | 2 | 0 | 3 | 1 | 17 |
| **Total** |  | **110** | **14** | **1** | **26** | **71** | **65** | **41** | **1** | **43** | **27** | **25** | **41** | **1** | **57** | **13** | **12** |

<sup>1</sup>Number of tagged birds *returning at least some data*. Two further birds that were tagged returned no data at all are *not* included in numbers.
<sup>2</sup>Includes birds that were predated, birds found dead, shot birds and birds with a 'mortality' signal (mortality signal indicates either the tag has been shed, or the bird has died).
<sup>3</sup>Includes limited data download (consistent with damage to the tag aerial) and total signal loss (without a mortality signal).

### Maximum dispersal distance

Considering all 110 tagged birds, the majority of birds travelled a maximum distance >500m from their release pen, representing 63%, 61% and 21% birds during the pre-shooting, shooting, and post-shooting phases, respectively. Accounting for mortality, this represents 63%, 94% and 85% of tagged birds alive at the start of pre-shooting (110 birds), shooting (71 birds), and post-shooting (27 birds) phases, respectively.

Movement beyond 500m involved different pools of birds in different phases. When considering all 110 birds across all three phases, 73% of individuals exhibited a maximum distance >500m during at least one phase, ranging from 33 – 100% birds per pen.

Considering release pen cohorts, movement beyond 500m occurred in all 11 cohorts, during at least one phase, but the proportion of birds involved varied between pen cohorts: *stocked birds* (i.e. the original 110 birds at point of release) – pre-shooting 33-100%, shooting 17-90% birds, and post-shooting 0-50% birds; *surviving birds* (i.e. those bird still alive at the start of each phase) – pre-shooting 33-100%, shooting 67-100%, post-shooting 0-100%.

Across pen cohorts, *mean maximum distances* exceeded 500m in all phases, with birds travelling 863m (457-2,183m) during pre-shooting, 1,493m (877-2,899m) during shooting, and 1,307m (457-3,598m) during post-shooting (Table 4). The maximum distance travelled by any bird per cohort ranged from 949-8,794m during pre-shooting, 1,246-7,028m during shooting and 457-6,415m during post-shooting. See Supplementary Materials (S2, Section 1) for values of individual birds.

**Table 4.** *Mean (± SE) maximum distance* travelled from release pens for each site combined (all birds), males and females, during (a) pre-shooting phase, (b) shooting phase, (c) post-shooting phase. Note: data includes birds/tags that died/failed during the respective phases. * Denotes early release sites (July).

| Site | All birds |  | Male |  | Female |  |
| --- | --- | --- | --- | --- | --- | --- |
|  | No. birds | Mean max. dist. from pen (m) | No. male | Mean max. dist. from pen (m) | No. female | Mean max. dist. from pen (m) |
| A* | 10 | 1054 ± 73.3 | 5 | 955 ± 103.6 | 5 | 1153 ± 92.3 |
| B* | 10 | 2183 ± 793.6 | 5 | 1230 ± 112.4 | 5 | 3135 ± 1538.7 |
| C1* | 10 | 731 ± 154.8 | 5 | 475 ± 154.7 | 5 | 986 ± 226.5 |
| C2* | 11 | 806 ± 172.5 | 6 | 806 ± 194.4 | 5 | 807 ± 327 |
| D* | 10 | 1262 ± 262.1 | 5 | 961 ± 286.6 | 5 | 1563 ± 426.3 |
| E* | 10 | 532 ± 62.9 | 5 | 457 ± 72.7 | 5 | 608 ± 98.4 |
| F1 | 10 | 557 ± 171.8 | 5 | 883 ± 249.3 | 5 | 231 ± 132.5 |
| F2 | 10 | 479 ± 129.6 | 5 | 523 ± 228 | 5 | 435 ± 150.6 |
| G | 12 | 762 ± 192.1 | 6 | 597 ± 140.3 | 6 | 928 ± 362.9 |
| H | 11 | 666 ± 81.8 | 6 | 774 ± 80 | 5 | 536 ± 139.8 |
| I | 6 | 457 ± 186.7 | 4 | 592 ± 262.2 | 2 | 186 ± 22.6 |
| Range | 6 - 12 | 457 – 2183 | 4 - 6 | 457 - 1230 | 2 - 6 | 186 – 3135 |
| Mean | 10 ± 0.4 | 863 ± 151.4 | 5 ± 0.2 | 750 ± 73.8 | 5 ± 0.3 | 961 ± 249.9 |

(b)
|  | All birds |  | Male |  | Female |  |
| --- | --- | --- | --- | --- | --- | --- |
| Site | No. birds | Mean max. dist. from pen (m) | No. male | Mean max. dist. from pen (m) | No. female | Mean max. dist. from pen (m) |
| A* | 8 | 1747 ± 275.1 | 4 | 2043 ± 496.5 | 4 | 1451 ± 219.4 |
| B* | 9 | 2899 ± 735.4 | 5 | 2263 ± 752.2 | 4 | 3694 ± 1386.4 |
| C1* | 4 | 1308 ± 107.7 | 1 | 1009 | 3 | 1408 ± 58.3 |
| C2* | 6 | 877 ± 193.9 | 4 | 600 ± 107.3 | 2 | 1431 ± 183.3 |
| D* | 7 | 1991 ± 466.1 | 2 | 1854 ± 1190.2 | 5 | 2045 ± 558.5 |
| E* | 8 | 1603 ± 421.7 | 4 | 1609 ± 584.4 | 4 | 1597 ± 698.7 |
| F1 | 5 | 1036 ± 222.4 | 4 | 1014 ± 285.6 | 1 | 1128 |
| F2 | 4 | 1021 ± 103.8 | 2 | 1162 ± 84 | 2 | 880 ± 134.1 |
| G | 11 | 1305 ± 223.2 | 6 | 1031 ± 280.1 | 5 | 1635 ± 326.5 |
| H | 8 | 939 ± 150 | 5 | 1039 ± 209.5 | 3 | 772 ± 203.6 |
| I | 1 | 1699 | 1 | 1699 | 0 | NA |
| Range | 1 - 11 | 877 - 2899 | 1-6 | 600 - 2263 | 0-5 | 772 - 3694 |
| Mean | 6 ± 0.8 | 1493 ± 179.2 | 3 ± 0.5 | 1393 ± 158.5 | 3 ± 0.5 | 1604 ± 260.3 |

(c)
|  | All birds |  | Male |  | Female |  |
| --- | --- | --- | --- | --- | --- | --- |
| Site | No. birds | Mean max. dist. from pen (m) | No. male | Mean max. dist. from pen (m) | No. female | Mean max. dist. from pen (m) |
| A* | 3 | 1836 ± 690.7 | 2 | 1679 ± 1165.1 | 1 | 2150 |
| B* | 5 | 3598 ± 1111.4 | 3 | 2621 ± 1521.5 | 2 | 5062 ± 1353.4 |
| C1* | 2 | 936 ± 268.3 | 0 | NA | 2 | 936 ± 268.3 |
| C2* | 1 | 623 | 0 | NA | 1 | 623 |
| D* | 3 | 2122 ± 377.5 | 1 | 2392 | 2 | 1987 ± 610.7 |
| E* | 1 | 705 | 1 | 705 | 0 | NA |
| F1 | 1 | 784 | 1 | 784 | 0 | NA |
| F2 | 1 | 457 | 1 | 457 | 0 | NA |
| G | 6 | 1036 ± 336.6 | 1 | 422 | 5 | 1158 ± 383.8 |
| H | 3 | 689 ± 236.9 | 2 | 471 ± 160.3 | 1 | 1125 |
| I | 1 | 1590 | 1 | 1590 | 0 | NA |
| Range | 1 - 6 | 457 - 3598 | 0 - 3 | 422 - 2621 | 0 - 5 | 623 - 5062 |
| Mean | 2 ± 0.5 | 1307 ± 281.1 | 1 ± 0.3 | 1236 ± 286.7 | 1 ± 0.4 | 1863 ± 572.6 |

A number of birds travelled a maximum distance >1,000m from their release pen: 32%, 44% and 13% of the 110 tagged birds in the pre-shooting, shooting, and post-shooting phases respectively, or accounting for mortality 32%, 68% and 52% of tagged birds alive at the start of pre-shooting (110 birds), shooting (71 birds), and post-shooting (27 birds) phases respectively.

When considering all 110 birds across all three phases, 53% of individuals exhibited a maximum distance >1,000m during at least one phase, ranging from 17 – 90% birds per pen.

Considering release pen cohorts, this longer movement beyond 1,000m occurred in all 11 cohorts, during at least one phase (but not necessarily during each phase). The proportion of birds exhibiting a maximum distance >1000m varied between cohorts: *stocked birds* - pre-shooting phase 0-70% of tagged birds per pen; shooting 17-80% birds, and post-shooting 0-40% birds: *surviving birds* – pre-shooting 0-70%, shooting 33-100%, post-shooting 0-100%. See Supplementary Materials (S2, Section 1) for values of individual birds.

### Mean dispersal distance

Although most of the 110 tagged birds exhibited a single maximum distance >500m, fewer tagged birds were recorded with a *mean* distance >500m from their release pen, representing 6%, 24% and 12% of the 110 tagged birds in the pre-shooting, shooting, and post-shooting phases respectively (Table 5). Accounting for mortality, this represents 6%, 37% and 48% of tagged birds alive at the start of pre-shooting (110 birds), shooting (71 birds), and post-shooting (27 birds) phases respectively.

**Table 5.**
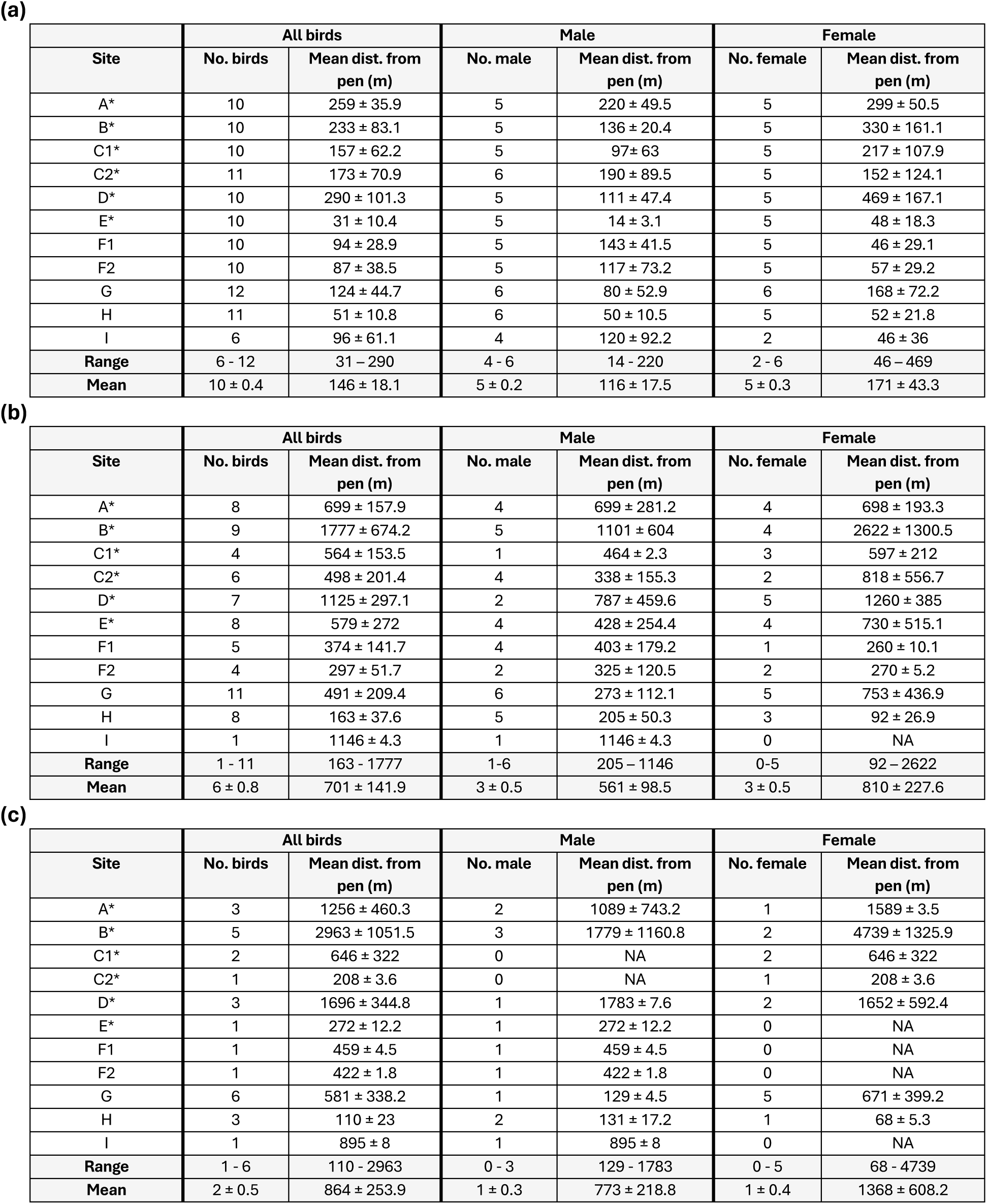
Mean (± SE) distance travelled from release pens for each site combined (all birds), males and females, during (a) pre-shooting phase, (b) shooting phase, (c) post-shooting phase. Note: data includes birds/tags that died/failed during the respective phases. * Denotes early release sites (July).

| Site | All birds |  | Male |  | Female |  |
| --- | --- | --- | --- | --- | --- | --- |
|  | No. birds | Mean dist. from pen (m) | No. male | Mean dist. from pen (m) | No. female | Mean dist. from pen (m) |
| A* | 10 | 259 ± 35.9 | 5 | 220 ± 49.5 | 5 | 299 ± 50.5 |
| B* | 10 | 233 ± 83.1 | 5 | 136 ± 20.4 | 5 | 330 ± 161.1 |
| C1* | 10 | 157 ± 62.2 | 5 | 97 ± 63 | 5 | 217 ± 107.9 |
| C2* | 11 | 173 ± 70.9 | 6 | 190 ± 89.5 | 5 | 152 ± 124.1 |
| D* | 10 | 290 ± 101.3 | 5 | 111 ± 47.4 | 5 | 469 ± 167.1 |
| E* | 10 | 31 ± 10.4 | 5 | 14 ± 3.1 | 5 | 48 ± 18.3 |
| F1 | 10 | 94 ± 28.9 | 5 | 143 ± 41.5 | 5 | 46 ± 29.1 |
| F2 | 10 | 87 ± 38.5 | 5 | 117 ± 73.2 | 5 | 57 ± 29.2 |
| G | 12 | 124 ± 44.7 | 6 | 80 ± 52.9 | 6 | 168 ± 72.2 |
| H | 11 | 51 ± 10.8 | 6 | 50 ± 10.5 | 5 | 52 ± 21.8 |
| I | 6 | 96 ± 61.1 | 4 | 120 ± 92.2 | 2 | 46 ± 36 |
| Range | 6 - 12 | 31 - 290 | 4 - 6 | 14 - 220 | 2 - 6 | 46 - 469 |
| Mean | 10 ± 0.4 | 146 ± 18.1 | 5 ± 0.2 | 116 ± 17.5 | 5 ± 0.3 | 171 ± 43.3 |
|  | No. birds | Mean dist. from pen (m) | No. male | Mean dist. from pen (m) | No. female | Mean dist. from pen (m) |
| A* | 8 | 699 ± 157.9 | 4 | 699 ± 281.2 | 4 | 698 ± 193.3 |
| B* | 9 | 1777 ± 674.2 | 5 | 1101 ± 604 | 4 | 2622 ± 1300.5 |
| C1* | 4 | 564 ± 153.5 | 1 | 464 ± 2.3 | 3 | 597 ± 212 |
| C2* | 6 | 498 ± 201.4 | 4 | 338 ± 155.3 | 2 | 818 ± 556.7 |
| D* | 7 | 1125 ± 297.1 | 2 | 787 ± 459.6 | 5 | 1260 ± 385 |
| E* | 8 | 579 ± 272 | 4 | 428 ± 254.4 | 4 | 730 ± 515.1 |
| F1 | 5 | 374 ± 141.7 | 4 | 403 ± 179.2 | 1 | 260 ± 10.1 |
| F2 | 4 | 297 ± 51.7 | 2 | 325 ± 120.5 | 2 | 270 ± 5.2 |
| G | 11 | 491 ± 209.4 | 6 | 273 ± 112.1 | 5 | 753 ± 436.9 |
| H | 8 | 163 ± 37.6 | 5 | 205 ± 50.3 | 3 | 92 ± 26.9 |
| I | 1 | 1146 ± 4.3 | 1 | 1146 ± 4.3 | 0 | NA |
| Range | 1 - 11 | 163 - 1777 | 1-6 | 205 - 1146 | 0-5 | 92 - 2622 |
| Mean | 6 ± 0.8 | 701 ± 141.9 | 3 ± 0.5 | 561 ± 98.5 | 3 ± 0.5 | 810 ± 227.6 |
|  | No. birds | Mean dist. from pen (m) | No. male | Mean dist. from pen (m) | No. female | Mean dist. from pen (m) |
| A* | 3 | 1256 ± 460.3 | 2 | 1089 ± 743.2 | 1 | 1589 ± 3.5 |
| B* | 5 | 2963 ± 1051.5 | 3 | 1779 ± 1160.8 | 2 | 4739 ± 1325.9 |
| C1* | 2 | 646 ± 322 | 0 | NA | 2 | 646 ± 322 |
| C2* | 1 | 208 ± 3.6 | 0 | NA | 1 | 208 ± 3.6 |
| D* | 3 | 1696 ± 344.8 | 1 | 1783 ± 7.6 | 2 | 1652 ± 592.4 |
| E* | 1 | 272 ± 12.2 | 1 | 272 ± 12.2 | 0 | NA |
| F1 | 1 | 459 ± 4.5 | 1 | 459 ± 4.5 | 0 | NA |
| F2 | 1 | 422 ± 1.8 | 1 | 422 ± 1.8 | 0 | NA |
| G | 6 | 581 ± 338.2 | 1 | 129 ± 4.5 | 5 | 671 ± 399.2 |
| H | 3 | 110 ± 23 | 2 | 131 ± 17.2 | 1 | 68 ± 5.3 |
| I | 1 | 895 ± 8 | 1 | 895 ± 8 | 0 | NA |
| Range | 1 - 6 | 110 - 2963 | 0 - 3 | 129 - 1783 | 0 - 5 | 68 - 4739 |
| Mean | 2 ± 0.5 | 864 ± 253.9 | 1 ± 0.3 | 773 ± 218.8 | 1 ± 0.4 | 1368 ± 608.2 |

Sustained movement beyond 500m involved different pools of birds in different phases. When considering all 110 birds across all three phases 25% (0 - 50% per pen) of individuals exhibited a mean distance >500m during at least one phase, ranging from 0 – 50% birds per pen.

This sustained movement beyond 500m occurred in up to 9 pens per phase during at least one phase (but not necessarily during each phase) but the proportion of birds involved varied between pen cohorts: *stocked birds* - pre-shooting phase 0-30% of tagged birds per pen; shooting 0-50% of birds per pens, and post-shooting 0-40% of birds per pen; *surviving birds* - pre-shooting 0-30%, shooting 0-100%, post-shooting 0-100%.

Some birds were also observed with a mean distance >1,000m from their release pen: 0%, 16% and 8% of the 110 tagged birds in the pre-shooting, shooting, and post-shooting phases respectively, or 0%, 24% and 33% of tagged birds alive at the start of pre-shooting (110 birds), shooting (71 birds), and post-shooting (27 birds) phases respectively.

When considering all 110 birds across all three phases, 16% (0 - 50% per pen) of individuals exhibited a mean distance >1,000m during at least one phase.

Across release pen cohorts, this longer sustained movement >1,000m occurred in up to 8 pens per phase but the proportion of birds involved varied across release pen cohorts: *stocked birds* - pre-shooting phase 0% of tagged birds; shooting 0-50% of birds per pen, and post-shooting 0-30% of birds per pen; *surviving birds* – pre-shooting 0%, shooting 0-100%, post-shooting 0-100%.

Mean dispersal distance of GPS-tagged pheasants varied considerably across study phases, and between release sites (Table 5). Individual estimates increased from between 2.0 ± 0.3m (site D) and 924.0 ± 39.7m (site B) during the pre-shooting phase, to between 29.0 ± 1.4m (site G) and 6,027 ± 4.7m (site B) during the shooting phase, and to between 68.0 ± 5.3m (site H) and 6,065 ± 6.3 (site B) during the post-shooting phase (see Supplementary Materials S2, Section 1) for mean values of individual birds). Accounting for individual-level variations in dispersal behaviours, nested across release sites, analysis revealed a statistically significant two-way interaction between study phase and release site (c^2^ = 3,5214.4, d.f. = 20, *p* < 0.0001). All sites showed a significant increase between the pre-shooting and shooting phases, most notably at sites A, B, D, E and G, (*p* < 0.0001). Between the shooting and post-shooting phase, dispersal continued to increase significantly at site A, B, E and G, as well as sites C1, F1 and F2 (*p* < 0.05 for all), although these latter sites may be a consequence of mortality with only a single bird surviving to the post-shooting phase. At site D, dispersal remained stable (coefficient = 0.037, *p* = 0.143), while sites H and I showed significant decreases (*p* < 0.0001). Dispersal also decreased for site C2, although this was statistically non-significant (coefficient = -0.060, *p* = 0.138)

No statistically significant difference in dispersal was found between sexes (c^2^ = 0.4, d.f. = 1, *p* = 0.539), with estimates comparable between males and females overall (Table 5).

Analysis of dispersal, relative to site-specific management conditions (i) release date, (ii) pen stocking density, (iii) total estate stocking density, and (iv) quality of release pen vegetative cover revealed statistically significant differences. Earlier release dates were linked to greater dispersal (*coefficient* = -0.024, *p* = 0.006), with early releases predicted to travel up to 71% further. Mean dispersal was positively associated with pheasant density, both within pens (*coefficient* = 0.001, *p* < 0.0001) and across estates (*coefficient* = 0.084, *p* = 0.004) and negatively associated with pen vegetative habitat quality (*coefficient* = -0.001, *p* = 0.045).

### Ranging profile

Ranging profiles of GPS-tagged pheasants varied significantly across study phases (distance bin x study phase interaction: c^2^ = 70.7, d.f. = 20, *p* < 0.0001). During the pre-shooting phase, a significantly greater proportion of GPS fixes were, on average, recorded within 500m of all release pens (90.16 ± 1.66% of fixes), when compared to those recorded within 500 – 1,000m (6.68 ± 1.16% of fixes; *coefficient* = 1.625, *p* < 0.001) and beyond 1,000m (3.16 ± 0.98% of fixes; *coefficient* = 1.283, *p* < 0.001).This is consistent with post-release acclimation of birds to the local environment and management actions (e.g. within-pen feeding and ‘dogging-in’) designed to initially retain birds close to the pen.

As shooting commenced, the proportion of fixes within 500m of release pens decreased to 65.91 ± 4.62%, and to 61.54 ± 9.24% during the post-shooting phase, although they were not statistically significant (pre-shooting – shooting: coefficient = -0.262, p = 0.051; shooting – post-shooting: coefficient = -0.32, p = 0.271). A statistically significant decrease in the proportion of fixes recorded within 500m of release pens was identified overall (pre-shooting – post-shooting: coefficient = 0.583, p = 0.009). The decrease in fixes recorded within 500m of release pens, coincided with a statistically significant increase in the proportion of fixes recorded beyond 1,000m between pre-shooting (3.16 ± 0.98%), and shooting (21.87 ± 4.32%; coefficient = 0.617, p = 0.011), increasing further still during the post-shooting phase (36.96 ± 8.78%), although not significantly (coefficient = 0.321, p = 0.447) when compared to the shooting phase. Again, a statistically significant increase in the proportion of GPS fixes recorded beyond 1,000m was identified overall (pre-shooting – post-shooting; coefficient = 0.938, p = 0.001).

Overall, of the 110 tagged birds released, 6%, 24% and 13% spent >50% of their time >500m from their release pen, during the pre-shooting, shooting, and post-shooting phases respectively. Across all three phases, 26% of the original 110 stocked birds spent >50% of their time >500m from the release pen, during at least one of the three phases (but not necessarily all phases) (see Supplementary Materials S2, Section 2).

Between release pen cohorts, time spent at distance occurred in most cohorts (9 of the 11) during at least one phase, but the proportion of original tagged birds (6 – 12 birds per pen) involved per cohort varied: pre-shooting phase 0-30% of tagged birds per pen; shooting 0-50% birds per pen, and post-shooting 0-50% birds per pen.

A number of the 110 tagged birds spent >50% of their time >1,000m from their release pen, representing 2%, 16% and 9% of birds during pre-shooting, shooting, and post-shooting phases respectively. Across all three phases, 16% of the original 110 stocked birds spent >50% of their time >1,000m from the release pen.

Between release pen cohorts, time spent at distance occurred in up to 9 cohorts during at least one phase, but the proportion of original tagged birds (6 – 12 birds per pen) involved per cohort varied: pre-shooting phase 0-20% of tagged birds; shooting 0-50% birds per pen, and post-shooting 0-30% birds per pen.

Considering mortality, for those birds spending >50% of their time >500m from their pen, this represents 6%, 37% and 52% of tagged birds alive at the start of pre-shooting (110 birds), shooting (71 birds), and post-shooting (27 birds) phases respectively (Table 6). Likewise for those birds spending >50% of their time >1,000m from their release pen, this represents 2%, 25% and 37% of birds alive.

**Table 6.** Summary of the number of birds spending >50% of their total time beyond 500m and 1,000m from the release pen. Note: Percentage values are calculated using the number of birds known to be alive at the start of each phase (i.e., 110, 71 and 27 for pre-shooting, shooting and post-shooting respectively).

|  | Pre-shooting |  |  |  |  | Shooting |  |  |  |  | Post-shooting |  |  |  |  |
| --- | --- | --- | --- | --- | --- | --- | --- | --- | --- | --- | --- | --- | --- | --- | --- |
|  |  | >50% >500 m |  | >50% >1000 m |  |  | >50% >500 m |  | >50% >1000 m |  |  | >50% >500 m |  | >50% >1000 m |  |
| Site | No. birds | No. | % | No. | % | No. birds | No. | % | No. | % | No. birds | No. | % | No. | % |
| A | 10 | 1 | 10.0 | 0 | 0.0 | 8 | 5 | 62.5 | 2 | 25.0 | 3 | 2 | 66.7 | 2 | 66.7 |
| B | 10 | 0 | 0.0 | 0 | 0.0 | 9 | 5 | 55.6 | 5 | 55.6 | 5 | 5 | 100.0 | 3 | 60.0 |
| C1 | 10 | 1 | 10.0 | 0 | 0.0 | 4 | 1 | 25.0 | 1 | 25.0 | 2 | 1 | 50.0 | 1 | 50.0 |
| C2 | 11<sup>1</sup> | 2 | 18.2 | 0 | 0.0 | 6 | 3 | 50.0 | 1 | 16.7 | 1 | 0 | 0.0 | 0 | 0.0 |
| D | 10 | 3 | 30.0 | 2 | 20.0 | 7 | 5 | 71.4 | 4 | 57.1 | 3 | 3 | 100.0 | 3 | 100.0 |
| E | 10 | 0 | 0.0 | 0 | 0.0 | 8 | 3 | 37.5 | 2 | 25.0 | 1 | 0 | 0.0 | 0 | 0.0 |
| F1 | 10 | 0 | 0.0 | 0 | 0.0 | 5 | 1 | 20.0 | 1 | 20.0 | 1 | 0 | 0.0 | 0 | 0.0 |
| F2 | 10 | 0 | 0.0 | 0 | 0.0 | 4 | 0 | 0.0 | 0 | 0.0 | 1 | 0 | 0.0 | 0 | 0.0 |
| G | 12 | 0 | 0.0 | 0 | 0.0 | 11 | 2 | 18.2 | 1 | 9.1 | 6 | 2 | 33.3 | 1 | 16.7 |
| H | 11 | 0 | 0.0 | 0 | 0.0 | 8 | 0 | 0.0 | 0 | 0.0 | 3 | 0 | 0.0 | 0 | 0.0 |
| I | 6 | 0 | 0.0 | 0 | 0.0 | 1 | 1 | 100.0 | 1 | 100.0 | 1 | 1 | 100.0 | 0 | 0.0 |
| Total | 110 | 7 | 6.4 | 2 | 1.8 | 71 | 26 | 36.6 | 18 | 25.4 | 27 | 14 | 51.9 | 10 | 37.0 |
| Mean ± SE | 10 ± 0.4 | 0.6 ± 0.3 |  | 0.2 ± 0.2 |  | 6.5 ± 0.8 | 2.4 ± 0.6 |  | 1.6 ± 0.5 |  | 2.5 ± 0.5 | 1.3 ± 0.5 |  | 0.9 ± 0.4 |  |

Across individual sites, dispersal profiles were shown to vary significantly overall (distance bin x release site interaction: c^2^ = 26.4, d.f. = 4, *p* < 0.001), but not in conjunction with the inclusion of study phases within the interaction term (study phase x distance bin x release site interaction: c^2^ = 37.6, d.f. = 33, *p* = 0.268). For certain sites, the proportion of GPS fixes were largely limited to within 500m of a release pen, such as site A, F2, H and G (p < 0.05 for all when compared to other distance bins); however, this was likely to be largely driven by restricted movement during the pre-shooting phase (see Table 6). For other sites, the proportion of GPS fixes, recorded across distance bins, were statistically comparable (i.e. p > 0.05), suggesting more consistent ranging across the available landscape, especially for site D and I where fixes beyond 1,000m were comparable to those recorded within 500m (p = 0.845 and 0.318 respectively). However, in the latter case this may be attributed to significant mortality following the pre-shooting phase (Table 6). Ranging profiles did not differ statistically between sexes (c^2^ = 0.6, d.f. = 1, *p* = 0.432).

### Direction of dispersal

Directional analysis revealed that, overall, pheasants exhibited non-random dispersal patterns, with statistically significant variation across release sites (c^2^ = 645.174, d.f. =70, *p* < 0.0001) (Table 7). However, no significant interaction was identified between dispersal direction and study phase (c^2^ = 23.571, d.f. = 4, *p* = 0.052), suggesting that the overall direction in which birds dispersed did not systematically change between the pre-shooting, shooting and post-shooting phases. However, males showed consistently more directed movement than females (c^2^ = 5.087, d.f. = 1, *p* = 0.024).

**Table 7.**
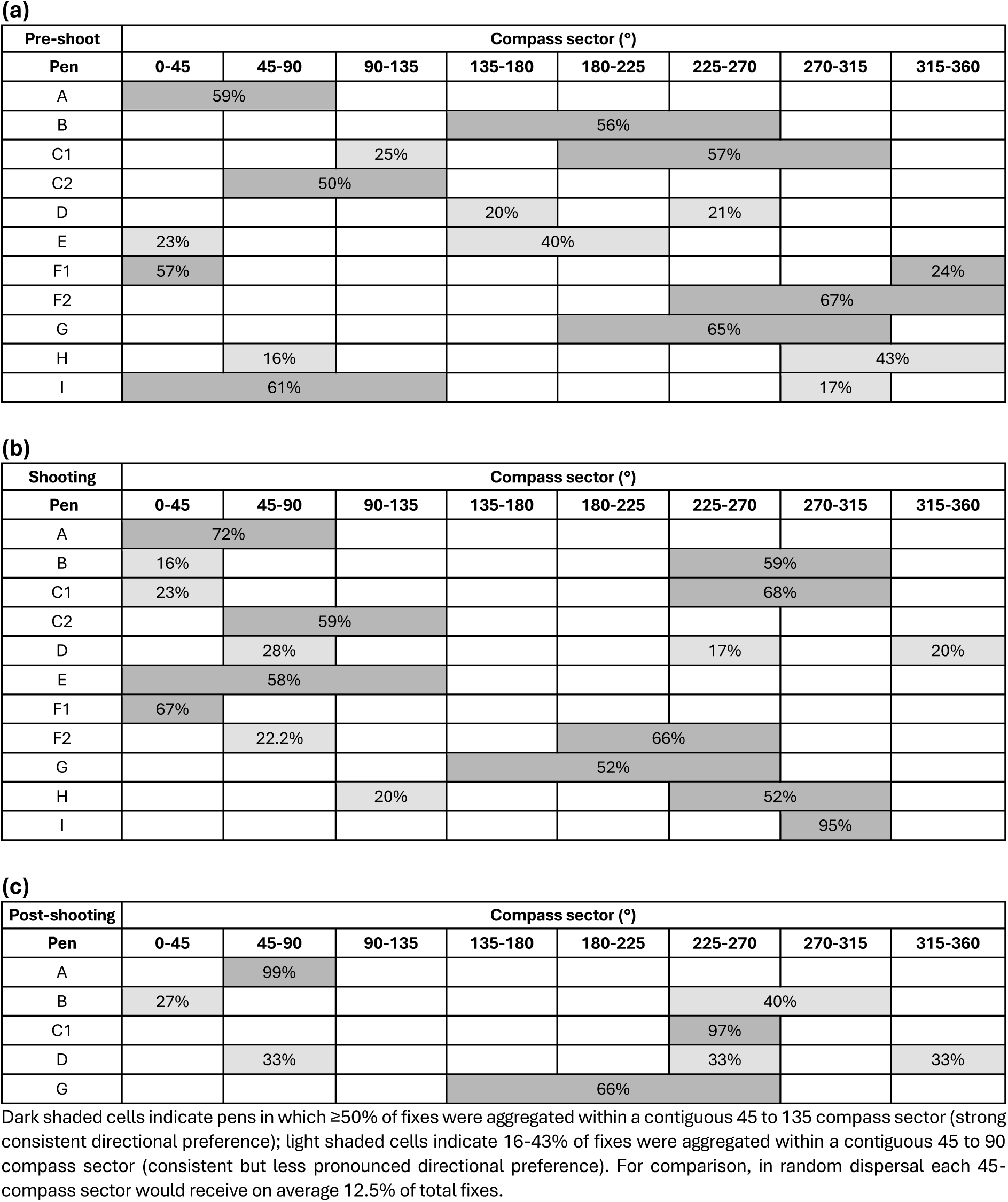
Summary of mean directional preference of birds from each estate/pen during the pre-shooting, shooting, and post-shooting phases. Post-shooting results are presented only for sites with multiple birds still alive during this phase, and some of these sites only had a small number of birds remaining. Values represent the mean percentage of total GPS fixes within a compass sector/s averaged across mean values for individual birds in a pen cohort.

Across all phases, most sites exhibited consistent directional preference, with mean GPS fixes across birds in a pen cohort aggregated within one (45 ° arc) to three (135 °) adjacent compass octants. During pre-shooting, 8 of 11 sites (73%) showed ≥50% of fixes concentrated within a contiguous 45-135° sector. This increased to 10 of 11 sites (91%) during shooting. Of the five sites with more than one bird alive during post-shooting, three (60%) showed birds had a consistent directional preference.

For data on individual birds see Supplementary Materials S2, Section 3.

### Conservation site incursions

Across all study phases, 28 (25%) of the 110 GPS-tagged pheasants from 7 (64%) of the 11 release pens (range 1-11 or 10-92% of birds per pen) spent time on designated conservation sites. Statistical modelling revealed a significant interaction between release site and dispersal distance in predicting the likelihood of incursion (c^2^ = 1,064.742, d.f. = 7, *p* < 0.0001), with proximity to conservation sites being a key driver. Birds released closer to conservation boundaries were more likely to encroach (*coefficient* = -0.544, *p* < 0.001), exemplified at site G with the highest number of intrusions, and the pen closest (149m) to a site designated as an SSSI, SPA and SAC. Compared to site G, incursions were significantly lower at sites A, B, and F1 (*p* < 0.05 for all), with proximity ranging from 455 – 872m. For the remaining four sites (proximity to a conservation site ranged from 414 – 1,744m), where in each only a single bird was responsible for intrusions, it was not possible for effective statistical comparison.

Study phase significantly influenced incursion frequency with incursions greater during the shooting phase, compared to both pre-shooting (*coefficient* = 1.012, *p* < 0.0001) and post-shooting (*coefficient* = 0.169, *p* = 0.0006). This pattern is consistent with the increased ranging behaviour during the shooting season, as birds moved more extensively throughout the landscape, and, subsequently, the markedly lower number of birds surviving to the post-shooting phase. No statistically significant difference was found between sexes (c^2^ = 0.009, d.f. = 1, *p* = 0.923), with males and females equally likely to encroach upon conservation sites.

During the pre-shooting phase, 15 of the 110 birds (14%) from 6 of 11 release pens (55%) entered conservation areas, typically at the nearest site (149 – 879m). Site G had the highest incidence, with encroachment recorded in 75% of all tagged birds. Shooting phase incursions increased to 23 birds (32% of surviving birds) from 7 of 11 release sites, with 52% being new incursions and 48% representing returns. During post-shooting, 10 birds (37% of surviving birds) from 7 of 11 release sites spent time on conservation sites, with only a single new incursion recorded.

Across all phases, most incursions involved birds accessing the nearest conservation site, though some individuals reached more distant sites (up to 2,319m), with site G having the greatest number of intruding individuals. Time spent on designated sites varied widely between individuals, from <1% (Site A) to nearly 49% of tracking time (Site I). Results are summarised in Table 8.

**Table 8.** Summary of intrusion metrics for each study phase (pre-shooting, shooting, post-shooting). Note: Some intruding birds were the same birds across phases. In total, 29 individual birds were recorded spending time on Conservation Sites across all study phases.

|  | Pre-shooting | Shooting | Post-shooting |
| --- | --- | --- | --- |
| Birds at start of phase | 110 | 71 | 27 |
| No. birds intruding from original stock | 15 | 23 | 10 |
| % birds intruding from original stock | 14 | 21 | 9 |
| % intruding from birds alive at the start of phase | 14 | 32 | 37 |
| % pens with $\geq 1$ bird intruding onto CS | 55 | 64 | 64 |
| Range of birds intruding across pens (%) | 10-75 | 10-75 | 10-33 |
| Distance range of CS from pen (m) | 149-1802 | 149-2319 | 149-2319 |
| Range of time spent on CS (%) | 0.10-28.8 | 0.03-48.9 | 0.60-24.4 |

## Discussion

Spatial data was collected on the movements of 110 reared and released pheasants, fitted with GPS tracking tags, across eleven release pens on nine working shoots, located in a range of landscape types. The principal aim was to investigate the dispersal behaviour and movement patterns of pheasants from woodland release pens, to address evidence gaps relating to the nature and magnitude of dispersal, with a focus on the extent of activity beyond 500m from the point of release.

### Maximum dispersal distance

Of the 110 tagged birds, 73% travelled a maximum distance >500m, and 53% travelled a maximum distance >1,000m from their release pen. For release pen cohorts, mean maximum distances exceeded 500m in all phases (pre-shooting: 863m, range: 457-2,183m; shooting: 1,493m, range: 877-2899m; post-shooting: 1,307m, range: 457-3598m), with one bird travelling almost 9km. These values broadly align with Turner (2008), who recorded radio-tagged pheasants travelling a mean maximum distance of 913 ± 82m during pre-shooting, though the current study’s upper maximum (8,794m) was nearly double that of Turner’s (4,685m). Beardsworth et al. (2021) found most pheasants released in August remained near pens until mid-September, only venturing further several weeks after release. The development of longer dispersal movements with increased post-release time is consistent with the current study’s findings that releases earlier in the year dispersed further. Shooting pressure and competition for resources may also promote longer movements (Swift et al., 2021), the latter consistent with the current findings that dispersal was greater at pens with higher stocking densities and poorer quality of vegetative habitat in pens. The Game and Wildlife Conservation Trust (GWCT) recommend a release pen stocking density of 1,000 birds/hectare of pen for plantations woodlands, but a reduced stocking density of up to 700 birds/hectare, for those sites proximate to conservation sites, which are inherently more vulnerable to ecological impacts (GWCT, 2021). In the current study, pen stocking density ranged from 347 – 3,614 birds per ha, with 6 of the 11 pens holding >1,000 birds per ha.

Although females tended to disperse further than males, which has also been reported in past studies (Beardsworth et al., 2021; Hill & Ridley, 1987), in the current study these differences were not statistically significant.

### Mean dispersal distance

The distances observed in the current study indicate pheasant dispersal was not consistent with the Defra Gamebird Review (GOV.UK, 2020) conclusion that ‘…*dispersal of birds tends to be less than 500m from the release sites*…’. Although birds stayed close to pens shortly after release, i.e. during the pre-shooting phase (pen cohort mean: 145m range: 31-290m), mean distances increased beyond 500m in subsequent phases (shooting: 701m, range 163-1,777m; post-shooting: 864m, range 110-2,963m). Of the 110 tagged individuals, 25% travelled a mean distance >500m, and 16% travelled a mean distance >1,000m from their release pen.

The pre-shooting phase was analysed as a single period, likely underestimating early dispersal due to initial acclimation and management to retain birds near pens. Sage et al. (2001) reported mean dispersal distances of 738 ± 131m for wild-type and 512 ± 66m for farm-type female pheasants over an extended period (August-April). These distances are smaller (particularly farm-type) than the mean dispersal for female pheasants in the current study, during a similar period: shooting 810 ± 228m and post-shooting phases 1,360 ± 608m.

### Ranging profile

In addition to mean and maximum distance, the current study also assessed time spent ‘at distance’ from release pens, as ecological risk posed by pheasants will depend on both distance and duration of activity. Of the 110 birds released, the proportion spending more than 50% of their time beyond 500m from the release pen was 6% during pre-shooting, 24% during shooting and 13% post-shooting; for distance beyond 1,000m, the proportions were 2%, 16% and 9%. Across all three phases, 25% of the 110 tagged birds spent >50% of their time beyond 500m, and 16% beyond 1,000m from their release pen, during any one phase.

In respect to the birds surviving at the start of each phase, the proportion spending >50% of their time beyond 500m increased markedly from 6% in pre-shooting to 37% during shooting and 52% post-shooting; for distances beyond 1,000m the proportions were 2%, 25% and 37%. This activity was represented across most release pens. During the shooting phase, birds from nine of the eleven release pens spent >50% of their time beyond both 500m and 1,000m. During this phase, in pens with multiple surviving birds (10 of the 11 pens; one pen had a single bird) 0-71% of birds spent >50% of their time beyond 500m, and 0-57% of birds spent >50% of their time beyond 1,000m.

Although patterns varied between sites, moderate percentages of birds spending prolonged periods at distance can scale up substantially at pen and estate level. For the most extreme case in the study, at site D, 50% of tagged birds from that release pen spent >50% of their time beyond 500m, and 40% of tagged birds spent >50% of their time beyond 1,000m, during at least one phase. When extrapolated to the full pen cohort (2,500 birds), this equates to 1,250 birds, and 1,000 birds spending the majority of their time beyond 500m and 1,000m, respectively, from the release pen and potentially 7,000 birds and 5,600 birds beyond 500m and 1,000m, respectively across the estate (14,000 released from all pens). This highlights the potential extent of pheasant activity ‘at distance’ from release pens in some circumstances.

### Direction of dispersal

Simplistic models of pheasant dispersal (e.g., Sage et al., 2021) assume birds disperse evenly in all directions and therefore estimate low densities of pheasant at 500m from release pens (approximately 10 birds/ha if 1,000 are released). However, empirical data from the current study and similar research (e.g. Turner, 2008) indicate that dispersal is directional, rather than uniform. In the current study, tagged cohorts generally showed directional preference (defined as the average movement direction across all birds) rather than uniform spread. Eight of eleven sites exhibited a general preference towards a contiguous 45-135° compass sector during the pre-shooting phase, increasing to 10 of 11 sites during the shooting phase. During this latter phase, at 8 of these 11 sites preference was more focused with fixes aggregated within a narrower 45-90 ° arc.

This consistency across phases suggests dispersal direction was shaped primarily by site-specific environmental cues (such as habitat cover, topography, and supplementary feed), rather than temporal changes or shooting pressure. In contrast to distance, which increased significantly over time, the compass bearing of movement remained relatively stable, reflecting a form of spatial fidelity or route persistence in the birds’ ranging behaviour. Previous work (Sage et al., 2021; Turner, 2008) associate directed movement to habitat quality and management practices, with birds favouring cover crops or areas of higher quality and resource availability. Such non-uniform dispersal creates uneven densities across landscapes, potentially increasing ecological risk where protected sites fall within preferred dispersal direction. Overall, these findings suggest that while directionality of dispersal was not significantly influenced by study phase, it was strongly site-dependent and, in many cases, consistent across time.

### Conservation site incursions

All release sites in the study were located near designated conservation areas, including Sites of Special Scientific Interest (SSSI), Special Protected Areas (SPA) and Special Areas of Conservation (SAC).

Overall, of the 110 tagged birds, 28 (25%) from 7 pens (64%) encroached onto a Conservation Site during the overall study, representing 1 - 11 birds (10 – 92%) of the respective pen cohort. Considering mortality, the proportion of surviving tagged birds entering Conservation Sites increased across phases: 14% pre-shooting (15/110), 32% shooting (23/71), and 37% post-shooting (10/27), as did the time spent within protected areas.

Although the highest encroachment rate onto a conservation site was associated with close proximity to the release pen, three (43%) of the pens associated with encroaching birds were located >500m from the nearest Conservation Site. Of the total 28 individuals (from 7 pens) encroaching over the three phases of the study, 18 (64%) were associated with release pens <500m (149 – 455m) from a conservation site. However, 10 (56%) of these 18 birds were associated with release pen G, which was the closest (149m) to a Conservation Site. Conservation sites beyond 500m from a pen also experienced incursions from pheasants – 10 (36%) of all birds encroaching were associated with pens 872 – 2,319m from the release pen, with 3 birds (11%) encroaching onto sites located over 1.5km from a release pen.

### Gamebird survival

It is estimated that around 60% of pheasants released for shooting in the UK do not end up being shot (Madden et al., 2018), and previous studies suggest only about 10% survive to potentially supplement wild populations by the end of the season (Hoodless et al., 2001; Madden et al., 2018; Turner, 2008). In the current study, at least 25% (27/110) of pheasants survived to the post-shooting phase and 12% (13/110) survived until the end of data collection (March). Survival varied between release pens (0 – 20%) with those pens exhibiting greater survival potentially posing greater ecological risk; exacerbated if those pens are also associated with greater dispersal and directional preference toward neighbouring conservation sites.

### Considerations and future directions

Several factors should be considered when interpreting these findings. Estate-level information was often incomplete or inconsistently reported across shooting estates, with some managers reluctant to share details on management practices; where data were consistent across estates, they were incorporated into analyses when possible.

The pre-shooting phase was analysed as a single period, though it likely comprised an initial settlement stage followed by dispersal, which may have led to underestimation of activity at greater distances later in this phase. Exploratory analysis suggested that dispersal behaviour shifts between approximately 2.5- and 6.5-weeks post-release, varying by site. Future analysis using segmented regression (Kileen et al., 2014) and the inclusion of time as a fine-scale predictor could improve estimates of the timing of initial dispersal movements and habitat use. Methods such as step-selection functions (Thurfjell et al., 2014) may also provide more accurate insights when data is of a fine enough scale, with limited missing data.

Finally, although the study encompassed a range of landscape types and working shoots, replication across additional landscapes and estate types would allow further assessment of whether observed patterns are consistent under different ecological and management contexts and increase confidence in the generality of these findings.

## Ethical statement

All animal procedures were conducted in accordance with the Animals (Scientific Procedures) Act 1986 and approved by the APHA’s Animal Welfare and Ethical Review Board.

## Supporting information

Supplementary Material S1 - Methodology

Supplementary Material S2 - Results

## Acknowledgements

We thank the estate owners and gamekeepers for granting access to their sites and enabling the tagging and tracking of their pheasants. These permissions along with their cooperation in providing management information was invaluable to the success of this study.

## References

Alonso, M. E., Pérez, J. A., Gaudioso, V. R., Díez, C., & Prieto, R. (2005). Study of survival, dispersal and home range of autumn-released red-legged partridges (*Alectoris rufa*). British Poultry Science, 46(4), 401–406.

Bagliacca, M., Falcini, F., Porrini, S., Zalli, F., & Fronte, B. (2008). Pheasant (*Phasianus colchicus*) hens of different origin: dispersion and habitat use after release. Italian Journal of Animal Science, 7(3), 321–333.

Beardsworth, C. E., Whiteside, M. A., Capstick, L. A., Laker, P. R., Langley, E. J., Nathan, R., Orchan, Y., Toledo, S., Van Horik, J. O., & Madden, J. R. (2021). Spatial cognitive ability is associated with transitory movement speed but not straightness during the early stages of exploration. Royal Society Open Science, 8(3), 201758.

Brooks, M. E., Kristensen, K., van Benthem, K. J., Magnusson, A., Berg, C. W., Nielsen, A., Skaug, H. J., Maechler, M., & Bolker, B. (2017). glmmTMB: Balances speed and flexibility among packages for zero-inflated generalized linear mixed modeling. The R Journal, 9(2), 378–400.

Duarte, J., Farfán, M. Á., & Vargas, J. M. (2011). New data on mortality, home range, and dispersal of red-legged partridges (*Alectoris rufa*) released in a mountain range. European Journal of Wildlife Research, 57, 675–678.

Gatti, R. C., Dumke, R. T., & Pils, C. M. (1989). Habitat use and movements of female ring-necked pheasants during fall and winter. Journal of Wildlife Management, 462–475.

GOV.UK. (2020). Review of gamebird releases on and around European protected sites. https://www.gov.uk/government/publications/review-of-gamebird-releases-on-and-around-european-protected-sites

Gruychev, G. (2014). Radiotelemetric tracking of farm pheasant (*Phasianus colchicus*) released in the Pazardzhik region, Bulgaria. Acta Zoologica Bulgarica, 66, 425–429.

GWCT. (2021). Guidelines for sustainable gamebird releasing. Game & Wildlife Conservation Trust.

Hill, D. A., & Ridley, M. W. (1987). Sexual segregation in winter, spring dispersal and habitat use in the pheasant (*Phasianus colchicus*). Journal of Zoology, 212(4), 657–668.

Hoodless, A. N., Draycott, R. A. H., Ludiman, M. N., & Robertson, P. A. (2001). Effects of supplementary feeding on territoriality, breeding success and survival of pheasants. Journal of Applied Ecology, 36(1), 147–156. 10.1046/j.1365-2664.1999.00388.x

Kenward, R. (2001). A manual for wildlife radio tagging. London, UK: Academic Press.

Krauss, G. D., Graves, H. B., & Zervanos, S. M. (1987). Survival of wild and game-farm cock pheasants released in Pennsylvania. Journal of Wildlife Management, 51(3), 555–559.

Leif, A. P. (2005). Spatial ecology and habitat selection of breeding male pheasants. Wildlife Society Bulletin, 33(1), 130–141.

Madden, J. R. (2021). How many gamebirds are released in the UK each year? European Journal of Wildlife Research, 67(4). 10.1007/s10344-021-01508-z

Madden, J. R., Hall, A., & Whiteside, M. A. (2018). Why do many pheasants released in the UK die, and how can we best reduce their natural mortality? Eur J Wildl Res, 64(4), 40. 10.1007/s10344-018-1199-5

Madden, J. R., & Sage, R. B. (2020). Ecological consequences of gamebird releasing and management on lowland shoots in England: A review by Rapid Evidence Assessment for Natural England and the British Association of Shooting and Conservation. Natural England Evidence Review NEER016.

Mason, L. R., Bicknell, J. E., Smart, J., & Peach, W. J. (2020). The impacts of non-native gamebird release in the UK: an updated evidence review. RSPB Research Report No. 66.

Pérez, J. A., Alonso, M. E., Gaudioso, V. R., Olmedo, J. A., Díez, C., & Bartolomé, D. (2004). Use of radiotracking techniques to study a summer repopulation with red-legged partridge (*Alectoris rufa*) chick. Poultry Science, 83(6), 882–888.

R Core Team. (2022). https://www.R-project.org/

Ridley, M. W. (1983). The mating system of the pheasant (Phasianus colchicus) Unpublished D.Phil. thesis, University of Oxford].

Sage, R. B., Brewin, J., Stevens, D. C., & Draycott, R. A. H. (2021). Gamebird releasing and management in the UK: A review of ecological considerations, best practice management and delivering net biodiversity gain. Game & Wildlife Conservation Trust.

Sage, R. B., Hoodless, A. N., Woodburn, M. I. A., Draycott, R. A. H., Madden, J. R., & Sotherton, N. W. (2020). Summary review and synthesis: effects on habitats and wildlife of the release and management of pheasants and red-legged partridges on UK lowland shoots. Wildlife Biology(4), 1–12. 10.2981/wlb.00766

Sage, R. B., Robertson, P. A., & Wise, D. R. (2001). Survival and breeding success of two pheasant (*Phasianus colchicus*) strains released into the wild. Proceedings Perdix VII Tome 2 Game Wildlife Science, 18, 331–340.

Smith, S. A., Stewart, N. J., & Gates, J. E. (1999). Home ranges, habitat selection and mortality of ring-necked pheasants (*Phasianus colchicus*) in north-central Maryland. The American Midland Naturalist, 141(1), 185–197.

Swift, R. J., Anteau, M. J., Ellis, K. S., Ring, M. M., Sherfy, M. H., & Toy, D. L. (2021). Dispersal distance is driven by habitat availability and reproductive success in Northern Great Plains piping plovers. Movement ecology, 9, 1–14.

Thurfjell, H., Ciuti, S., & Boyce, M. S. (2014). Applications of step-selection functions in ecology and conservation. Mov Ecol, 2(1), 4. 10.1186/2051-3933-2-4

Turner, C. V. (2008). The fate and management of pheasants (Phasianus colchicus) released in the UK Unpublished PhD thesis, Imperial College, University of London].

Wilson, R. J., Drobney, R. D., & Hallett, D. L. (1992). Survival, dispersal, and site fidelity of wild female ring-necked pheasants following translocation. Journal of Wildlife Management, 56, 79–86.

