## Supplementary Material S1 - Methodology for "Dispersal Behaviour and Movement Patterns of Pheasants from Woodland Release Pens"

**Supplementary Material 1 (S1) - Methodology**

**Section 1: Tag data download and processing of GPS data**

At each study site, GPS data were downloaded from tags every one to two weeks and converted into usable formats through post-processing steps. Raw SWIFT fix files were decoded to produce KML files (for visualisation in Google Earth) and text files containing key details for each fix (e.g., date, time, latitude, longitude, satellite count). Once compiled, the GPS files were formatted and 'cleaned' using the R statistical programme (version 4.2.2: *R Core Team*, 2022) to ensure that only valid GPS fixes were retained and that they were compatible with the Geographic Information System platform (ArcGIS Pro (version 3.0.1). All associated data layers were projected in the British National Grid (EPSG: 27700)., The validity of GPS data was determined based on manufacturers specifications, in particular the requirement for a Horizontal Diffusion/Dilution of Precision (HDOP) value of five or less to be considered valid, and a minimum number of four or more satellites involved in each GPS fix. Any GPS fixes that did not conform to those requirements were deemed invalid and excluded from subsequent processing and analysis. Similarly, any fixes which did not return any information, likely due to a failure to connect to overhead satellites during the transmission window (12 – 40 seconds per fix), were also excluded from the dataset. Following initial cleaning and validation, 94% of GPS fixes (n = 96,428 / 102,177), recorded for all birds, were retained for analysis.

Further data screening was conducted to remove any potential outliers in respect to distance travelled, essentially removing any outlying fixes that occurred within a time frame considered too short (e.g. 60 minutes) to be a natural pheasant movement. This resulted in the removal of a single data point from each of two birds (from two independent pen cohorts) out of the 110 tagged individuals. Leaving a final retention of 96,426/102,177 GPS fixes.

GPS data were initially processed to determine the frequency with which GPS fixes occurred inside or outside the release pen and how this frequency changed over time. For those GPS fixes which occurred outside the release pen, the straight-line distance (i.e. Euclidean distance) between each fix and the boundary of the release pen was calculated. Measures of dispersal included: (1) maximum distance (single greatest straight-line distance from the release pen boundary for an individual bird); (2) mean distance (average distance measured from release pen boundary for an individual bird); (3) cohort mean distance (average distance measured from release pen boundary across all birds from the same pen). Distance was also represented as a binned value, ranging from 0 to 3km at 100m intervals, thereby allowing the amount of time each tagged bird spent at different distances, relative to the release pen, to be estimated.
