## Supplementary Material S2 - Results for "Dispersal Behaviour and Movement Patterns of Pheasants from Woodland Release Pens"

### Supplementary Material 2 (S2) - Results

**Section 1: *Maximum* distance moved (i.e., single largest straight-line distance measured from the boundary of a release pen) and *mean* distance moved (i.e., average distance measured from the boundary of a release pen) is presented for each bird. The total number of weeks each tag was active, before the bird or tag died (signal loss), is also presented. A value of 'NA' indicates no date, where birds/tags had died/failed before any data was collected for that phase.**

#### Site A

| Tagged | | Pre-shooting | | | | Shooting | | | | Post-shooting | | | |
|---|---|---|---|---|---|---|---|---|---|---|---|---|---|
| Bird ID | Sex | Max dist. from pen (m) [1] | Mean ± SE dist. from pen (m) [2] | Total weeks | Bird/tag outcome | Max dist. from pen (m) [1] | Mean ± SE dist. from pen (m) [2] | Total weeks | Bird/tag outcome | Max dist. from pen (m) [1] | Mean ± SE dist. from pen (m) [2] | Total weeks | Bird/tag outcome |
| 57142 | M | 790 | 135 ± 5.9 | 10 | Predated - fox | NA | NA | 10 | Predated - fox | NA | NA | 10 | Predated – fox |
| 57143 | F | 1076 | 334 ± 7.4 | 13 | Survived | 2043 | 1210 ± 10.7 | 29 | Survived | 2150 | 1589 ± 3.5 | 35 | Tag signal loss |
| 57144 | M | 1256 | 401 ± 10 | 13 | Survived | 3164 | 1533 ± 13 | 29 | Survived | 2844 | 1832 ± 11.7 | 35 | Tag signal loss |
| 57145 | M | 675 | 179 ± 4.7 | 13 | Survived | 1080 | 320 ± 2.5 | 29 | Survived | 513 | 346 ± 1.4 | 37 | Survived |
| 57146 | F | 904 | 116 ± 4.4 | 13 | Survived | 1048 | 280 ± 7.5 | 17 | Tag signal loss | NA | NA | 17 | Tag signal loss |
| 57147 | F | 1450 | 350 ± 6.9 | 13 | Tag signal loss | NA | NA | 13 | Tag signal loss | NA | NA | 13 | Tag signal loss |
| 57148 | F | 1251 | 415 ± 7.8 | 13 | Survived | 1509 | 715 ± 8.8 | 17 | Shot | NA | NA | 17 | Shot |
| 57149 | M | 1091 | 246 ± 5.5 | 13 | Survived | 2581 | 416 ± 9.5 | 19 | Shot | NA | NA | 19 | Shot |
| 57150 | F | 1084 | 279 ± 7.1 | 13 | Survived | 1204 | 589 ± 5.4 | 17 | Shot | NA | NA | 17 | Shot |
| 57151 | M | 962 | 138 ± 5.1 | 13 | Survived | 1348 | 526 ± 11.2 | 20 | Tag signal loss | NA | NA | 20 | Tag signal loss |
| *Mean ± SE* | *All* | *1054 ± 73.3* | *259 ± 35.9* | | | *1747 ± 275.1* | *699 ± 157.9* | | | *1836 ± 690.7* | *1256 ± 460.3* | | |
| | *M* | *955 ± 103.6* | *220 ± 49.5* | | | *2043 ± 496.5* | *699 ± 281.2* | | | *1679 ± 1165.1* | *1089 ± 743.2* | | |
| | *F* | *1153 ± 92.3* | *299 ± 50.5* | | | *1451 ± 219.4* | *698 ± 193.3* | | | *2150[3]* | *1589 ± 3.5* | | |

[1] The single largest straight-line distance recorded for each bird from the release pen.
[2] Calculated from the weekly average distance from the pen.
[3] Only one female survived to the post-shooting phase, thus SE not calculated.

### Site B

| Tagged | | Pre-shooting | | | | Shooting | | | | Post-shooting | | | |
|---|---|---|---|---|---|---|---|---|---|---|---|---|---|
| Bird ID | Sex | Max dist. from pen (m) [1] | Mean ± SE dist. from pen (m) [2] | Total weeks | Bird/tag outcome | Max dist. from pen (m) [1] | Mean ± SE dist. from pen (m) [2] | Total weeks | Bird/tag outcome | Max dist. from pen (m) [1] | Mean ± SE dist. from pen (m) [2] | Total week s | Bird/tag outcome |
| 57122 | F | 796 | 89 ± 4 | 13 | Survived | 802 | 101 ± 4.7 | 24 | Shot | NA | NA | 24 | Shot |
| 57123 | M | 1614 | 176 ± 7.4 | 13 | Survived | 4992 | 3357 ± 31.7 | 29 | Survived | 5597 | 4088 ± 41.5 | 36 | Survived |
| 57124 | M | 1183 | 129 ± 4.8 | 13 | Survived | 1462 | 336 ± 3.3 | 29 | Survived | 583 | 411 ± 7.8 | 31 | Tag signal loss |
| 57125 | M | 1177 | 186 ± 6 | 13 | Survived | 645 | 192 ± 3.9 | 19 | Tag signal loss | NA | NA | 19 | Tag signal loss |
| 57126 | M | 915 | 77 ± 3.6 | 13 | Survived | 2638 | 1370 ± 12.2 | 29 | Survived | 1683 | 839 ± 32.5 | 29 | Tag signal loss |
| 57127 | F | 3518 | 330 ± 18.2 | 13 | Survived | 4794 | 3168 ± 4.1 | 29 | Survived | 3709 | 3413 ± 5.4 | 34 | Tag signal loss |
| 57128 | M | 1260 | 112 ± 3.5 | 13 | Survived | 1577 | 250 ± 4.1 | 29 | Predated - fox | NA | NA | 29 | Predated - fox |
| 57129 | F | 87 | 3 ± 0.6 | 3 | Predated - fox | NA | NA | 3 | Predated - fox | NA | NA | 3 | Predated - fox |
| 57130 | F | 2482 | 305 ± 11.8 | 13 | Survived | 2153 | 1192 ± 20.3 | 17 | Tag signal loss | NA | NA | 17 | Tag signal loss |
| 57131 | F | 8794 | 924 ± 39.7 | 13 | Survived | 7028 | 6027 ± 4.7 | 29 | Survived | 6415 | 6065 ± 6.3 | 31 | Tag signal loss |
| *Mean ± SE* | *All* | *2183 ± 793.6* | *233 ± 83.1* | | | *2899 ± 735.4* | *1777 ± 674.2* | | | *3598 ± 1111.4* | *2963 ± 1051.5* | | |
| | *M* | *1230 ± 112.4* | *136 ± 20.4* | | | *2263 ± 752.2* | *1101 ± 604* | | | *2621 ± 1521.5* | *1779 ± 1160.8* | | |
| | *F* | *3135 ± 1538.7* | *330 ± 161.1* | | | *3694 ± 1386.4* | *2622 ± 1300.5* | | | *5062 ± 1353.4* | *4739 ± 1325.9* | | |

[1] The single largest straight-line distance recorded for each bird from the release pen.
[2] Calculated from the weekly average distance from the pen.

### Site C1

| Tagged | | Pre-shooting | | | | Shooting | | | | Post-shooting | | | |
|---|---|---|---|---|---|---|---|---|---|---|---|---|---|
| Bird ID | Sex | Max dist. from pen (m) [1] | Mean ± SE dist. from pen (m) [2] | Total weeks | Bird/tag outcome | Max dist. from pen (m) [1] | Mean ± SE dist. from pen (m) [2] | Total weeks | Bird/tag outcome | Max dist. from pen (m) [1] | Mean ± SE dist. from pen (m) [2] | Total week s | Bird/tag outcome |
| 57152 | F | 761 | 88 ± 2.5 | 16 | Survived | 1370 | 377 ± 4.5 | 29 | Survived | 668 | 324 ± 4 | 32 | Tag signal loss |
| 57153 | F | 1529 | 628 ± 10.4 | 16 | Survived | 1331 | 1021 ± 1.8 | 29 | Survived | 1204 | 968 ± 3.1 | 36 | Survived |
| 57154 | M | 99 | 20 ± 3.2 | 1 | Tag signal loss | NA | NA | 1 | Tag signal loss | NA | NA | 1 | Tag signal loss |
| 57155 | F | 452 | 13 ± 3 | 2 | Tag signal loss | NA | NA | 2 | Tag signal loss | NA | NA | 2 | Tag signal loss |
| 57156 | F | 663 | 140 ± 8.9 | 6 | Tag signal loss | NA | NA | 6 | Tag signal loss | NA | NA | 6 | Tag signal loss |
| 57157 | M | 359 | 7 ± 4 | 1 | Tag signal loss | NA | NA | 1 | Tag signal loss | NA | NA | 1 | Tag signal loss |
| 57158 | M | 404 | 29 ± 3.3 | 3 | Tag signal loss | NA | NA | 3 | Tag signal loss | NA | NA | 3 | Tag signal loss |
| 57159 | M | 473 | 87 ± 8.2 | 3 | Tag signal loss | NA | NA | 3 | Tag signal loss | NA | NA | 3 | Tag signal loss |
| 57160 | M | 1040 | 343 ± 4.4 | 16 | Survived | 1009 | 464 ± 2.3 | 24 | Tag signal loss | NA | NA | 24 | Tag signal loss |
| 57161 | F | 1525 | 218 ± 4.3 | 16 | Survived | 1522 | 393 ± 7 | 26 | Shot | NA | NA | 26 | Shot |
| *Mean ± SE* | *All* | *731 ± 154.8* | *157 ± 62.2* | | | *1308 ± 107.7* | *564 ± 153.5* | | | *936 ± 268.3* | *646 ± 322* | | |
| | *M* | *475 ± 154.7* | *97± 63* | | | *1009[3]* | *464 ± 2.3* | | | *NA[4]* | *NA[4]* | | |
| | *F* | *986 ± 226.5* | *217 ± 107.9* | | | *1408 ± 58.3* | *597 ± 212* | | | *936 ± 268.3* | *646 ± 322* | | |

[1] The single largest straight-line distance recorded for each bird from the release pen.
[2] Calculated from the weekly average distance from the pen.
[3] Only one male survived to the shooting phase, thus SE not calculated.
[4] No males survived to the post-shooting phase.

### Site C2

| Tagged | | Pre-shooting | | | | Shooting | | | | Post-shooting | | | |
|---|---|---|---|---|---|---|---|---|---|---|---|---|---|
| Bird ID | Sex | Max dist. from pen (m) [1] | Mean ± SE dist. from pen (m) [2] | Total weeks | Bird/tag outcome | Max dist. from pen (m) [1] | Mean ± SE dist. from pen (m) [2] | Total weeks | Bird/tag outcome | Max dist. from pen (m) [1] | Mean ± SE dist. from pen (m) [2] | Total weeks | Bird/tag outcome |
| 57162 | M | 1297 | 569 ± 7.9 | 15 | Survived | 821 | 664 ± 22.5 | 16 | Shot | NA | NA | 16 | Shot |
| 57163 | M | 1268 | 145 ± 4.5 | 15 | Survived | 733 | 542 ± 17.3 | 16 | Tag mortality signal | NA | NA | 16 | Tag mortality signal |
| 57164 | F | 1573 | 21 ± 17.9 | 2 | Tag shed | NA | NA | 2 | Tag shed | NA | NA | 2 | Tag shed |
| 57165 | M | 338 | 27 ± 3.9 | 4 | Bird mortality | NA | NA | 4 | Bird mortality | NA | NA | 4 | Bird mortality |
| 57166 | M | 348 | 33 ± 1.2 | 15 | Survived | 489 | 54 ± 2.2 | 25 | Shot | NA | NA | 25 | Shot |
| 57167 | F | 350 | 42 ± 1.4 | 15 | Survived | 1248 | 261 ± 4 | 28 | Survived | 623 | 208 | 36 | Survived |
| 57169 | M | 439 | 36 ± 1.3 | 15 | Survived | 356 | 91 ± 3 | 18 | Shot | NA | NA | 18 | Shot |
| 57170 | F | 1637 | 648 ± 12.5 | 15 | Survived | 1614 | 1375 ± 2.3 | 24 | Predated | NA | NA | 24 | Predated |
| 57171 | F | 273 | 13 ± 2.1 | 4 | Bird mortality | NA | NA | 4 | Bird mortality | NA | NA | 4 | Bird mortality |
| 57137* | F | 201 | 36 ± 7 | 1 | Bird mortality | NA | NA | 1 | Bird mortality | NA | NA | 1 | Bird mortality |
| 57165* | M | 1146 | 331 ± 5.3 | 11** | Predated - fox | NA | NA | 11** | Predated - fox | NA | NA | 11** | Predated - fox |
| *Mean ± SE* | *All* | *806 ± 172.5* | *173 ± 70.9* | | | *877 ± 193.9* | *498 ± 201.4* | | | *623*[3] | *208 ± 3.6* | | |
| | *M* | *806 ± 194.4* | *190 ± 89.5* | | | *600 ± 107.3* | *338 ± 155.3* | | | *NA*[4] | *NA*[4] | | |
| | *F* | *807 ± 327* | *152 ± 124.1* | | | *1431 ± 183.3* | *818 ± 556.7* | | | *623*[3] | *208 ± 3.6* | | |

[1] The single largest straight-line distance recorded for each bird from the release pen.
[2] Calculated from the weekly average distance from the pen.
[3] Only one female bird survived to the post shooting season thus no SE calculated.
[4] No male birds survived to the post shooting season.
*Tags already used on previous birds that were found dead very early and reapplied to new birds.
**Tag was reapplied one month after initial tagging date; hence why total weeks is less for this bird.

### Site D

| Tagged | | Pre-shooting | | | | Shooting | | | | Post-shooting | | | |
|---|---|---|---|---|---|---|---|---|---|---|---|---|---|
| Bird ID | Sex | Max dist. from pen (m) [1] | Mean ± SE dist. from pen (m) [2] | Total weeks | Bird/tag outcome | Max dist. from pen (m) [1] | Mean ± SE dist. from pen (m) [2] | Total weeks | Bird/tag outcome | Max dist. from pen (m) [1] | Mean ± SE dist. from pen (m) [2] | Total weeks | Bird/tag outcome |
| 57132 | F | 680 | 91 ± 2.9 | 13 | Survived | 927 | 228 ± 4.9 | 18 | Tag signal loss | NA | NA | 18 | Tag signal loss |
| 57133 | M | 1414 | 274 ± 6.9 | 13 | Survived | 664 | 327 ± 5.3 | 16 | Tag mortality signal | NA | NA | 16 | Tag mortality signal |
| 57134 | M | 729 | 51 ± 2.7 | 9 | Tag signal loss | NA | NA | 9 | Tag signal loss | NA | NA | 9 | Tag signal loss |
| 57135 | F | 1393 | 581 ± 13.6 | 13 | Survived | 1414 | 1057 ± 1.7 | 28 | Survived | 1377 | 1060 ± 2.5 | 35 | Survived |
| 57136 | M | 1711 | 151 ± 6.4 | 13 | Survived | 3044 | 1247 ± 15.8 | 28 | Survived | 2392 | 1783 ± 7.6 | 31 | Tag signal loss |
| 57137 | M | 57 | 2 ± 0.3 | 3 | Bird mortality | NA | NA | 3 | Bird mortality | NA | NA | 3 | Bird mortality |
| 57138 | F | 2526 | 879 ± 15.5 | 13 | Survived | 3556 | 1928 ± 39.1 | 19 | Tag mortality signal | NA | NA | 19 | Tag mortality signal |
| 57139 | F | 634 | 62 ± 1.9 | 13 | Survived | 1097 | 751 ± 4 | 25 | Tag signal loss | NA | NA | 25 | Tag signal loss |
| 57140 | F | 2580 | 732 ± 15.6 | 13 | Survived | 3232 | 2334 ± 3.2 | 28 | Survived | 2598 | 2245 ± 3.9 | 35 | Survived |
| 57141 | M | 895 | 79 ± 3.4 | 12 | Tag signal loss | NA | NA | 12 | Tag signal loss | NA | NA | 12 | Tag signal loss |
| *Mean ± SE* | *All* | *1262 ± 262.1* | *290 ± 101.3* | | | *1991 ± 466.1* | *1125 ± 297.1* | | | *2122 ± 377.5* | *1696 ± 344.8* | | |
| | *M* | *961 ± 286.6* | *111 ± 47.4* | | | *1854 ± 1190.2* | *787 ± 459.6* | | | *2392[3]* | *1783 ± 7.6* | | |
| | *F* | *1563 ± 426.3* | *469 ± 167.1* | | | *2045 ± 558.5* | *1260 ± 385* | | | *1987 ± 610.7* | *1652 ± 592.4* | | |

[1] The single largest straight-line distance recorded for each bird from the release pen.
[2] Calculated from the weekly average distance from the pen.
[3] Only one male bird survived to the post shooting season thus no SE calculated.

### Site E

| Tagged | | Pre-shooting | | | | Shooting[3] | | | | Post-shooting | | | |
|---|---|---|---|---|---|---|---|---|---|---|---|---|---|
| Bird ID | Sex | Max dist. from pen (m) [1] | Mean ± SE dist. from pen (m) [2] | Total weeks | Bird/tag outcome | Max dist. from pen (m) [1] | Mean ± SE dist. from pen (m) [2] | Total weeks | Bird/tag outcome | Max dist. from pen (m) [1] | Mean ± SE dist. from pen (m) [2] | Total weeks | Bird/tag outcome |
| 57202 | F | 386 | 37 ± 2.5 | 7 | Survived | 212 | 51 ± 4.2 | 9 | Tag signal loss | NA | NA | 9 | Tag signal loss |
| 57203 | M | 358 | 18 ± 1.9 | 7 | Survived | 802 | 120 ± 3.1 | 27 | Survived | 705 | 272 ± 12.2 | 29 | Survived |
| 57204 | M | 447 | 23 ± 2 | 7 | Survived | 2002 | 334 ± 6.6 | 24 | Tag mortality signal | NA | NA | 24 | Tag mortality signal |
| 57205 | F | 444 | 5 ± 1 | 7 | Survived | 3535 | 2240 ± 27.3 | 20 | Tag signal loss | NA | NA | 20 | Tag signal loss |
| 57206 | F | 633 | 22 ± 2.6 | 7 | Survived | 1195 | 92 ± 5.3 | 14 | Tag signal loss | NA | NA | 14 | Tag signal loss |
| 57207 | M | 475 | 9 ± 1.6 | 7 | Tag signal loss | NA | NA | 7 | Tag signal loss | NA | NA | 7 | Tag signal loss |
| 57208 | M | 288 | 7 ± 1 | 7 | Survived | 550 | 85 ± 4 | 12 | Predated - fox | NA | NA | 12 | Predated - fox |
| 57209 | F | 626 | 64 ± 10 | 2 | Predated - fox | NA | NA | 2 | Predated - fox | NA | NA | 2 | Predated - fox |
| 57210 | F | 949 | 110 ± 6.6 | 7 | Survived | 1445 | 538 ± 5.1 | 22 | Tag signal loss | NA | NA | 22 | Tag signal loss |
| 57211 | M | 716 | 16 ± 2.2 | 7 | Survived | 3083 | 1173 ± 13.1 | 23 | Tag signal loss | NA | NA | 23 | Tag signal loss |
| *Mean ± SE* | *All* | *532 ± 62.9* | *31 ± 10.4* | | | *1603 ± 421.7* | *579 ± 272* | | | *705[4]* | *272 ± 12.2* | | |
| | *M* | *457 ± 72.7* | *14 ± 3.1* | | | *1609 ± 584.4* | *428 ± 254.4* | | | *705[4]* | *272 ± 12.2* | | |
| | *F* | *608 ± 98.4* | *48 ± 18.3* | | | *1597 ± 698.7* | *730 ± 515.1* | | | *NA[5]* | *NA[5]* | | |

[1] The single largest straight-line distance recorded for each bird from the release pen.
[2] Calculated from the weekly average distance from the pen.
[3] Shooting season is based on the start of red legged partridge shooting as this takes place in the same area that pheasants are present.
[4] Only one male bird survived to the post shooting season thus no SE calculated.
[5] No female birds survived to the post shooting season.

### Site F1

| Tagged | | Pre-shooting | | | | Shooting | | | | Post-shooting | | | |
|---|---|---|---|---|---|---|---|---|---|---|---|---|---|
| Bird ID | Sex | Max dist. from pen (m)[1] | Mean ± SE dist. from pen (m)[2] | Total weeks | Bird/tag outcome | Max dist. from pen (m)[1] | Mean ± SE dist. from pen (m)[2] | Total weeks | Bird/tag outcome | Max dist. from pen (m)[1] | Mean ± SE dist. from pen (m)[2] | Total weeks | Bird/tag outcome |
| 57172 | M | 895 | 146 ± 2.7 | 14 | Survived | 790 | 281 ± 3 | 26 | Shot | NA | NA | 26 | Shot |
| 57173 | M | 660 | 124 ± 2.2 | 14 | Survived | 862 | 182 ± 3.5 | 22 | Tag signal loss | NA | NA | 22 | Tag signal loss |
| 57174 | M | 139 | 12 ± 1 | 3 | Tag signal loss | NA | NA | 3 | Tag signal loss | NA | NA | 3 | Tag signal loss |
| 57175 | F | 751 | 161 ± 2.7 | 14 | Survived | 1128 | 260 ± 10.1 | 18 | Shot | NA | NA | 18 | Shot |
| 57176 | M | 1663 | 271 ± 7.1 | 14 | Survived | 1847 | 937 ± 9.1 | 26 | Survived | 784 | 459 ± 4.5 | 30 | Survived |
| 57177 | F | 199 | 28 ± 2.2 | 4 | Faulty tag | NA | NA | 4 | Faulty tag | NA | NA | 4 | Faulty tag |
| 57178 | F | 60 | 11 ± 1 | 3 | Bird mortality | NA | NA | 3 | Bird mortality | NA | NA | 3 | Bird mortality |
| 57179 | M | 1057 | 163 ± 3.6 | 14 | Survived | 555 | 213 ± 34.5 | 15 | Shot | NA | NA | 15 | Shot |
| 57180 | F | 78 | 12 ± 1 | 3 | Bird mortality | NA | NA | 3 | Bird mortality | NA | NA | 3 | Bird mortality |
| 57181 | F | 67 | 16 ± 9.2 | 1 | Tag signal loss | NA | NA | 1 | Tag signal loss | NA | NA | 1 | Tag signal loss |
| *Mean ± SE* | *All* | *557 ± 171.8* | *94 ± 28.9* | | | *1036 ± 222.4* | *374 ± 141.7* | | | *784*[4] | *459 ± 4.5* | | |
| | *M* | *883 ± 249.3* | *143 ± 41.5* | | | *1014 ± 285.6* | *403 ± 179.2* | | | *784*[4] | *459 ± 4.5* | | |
| | *F* | *231 ± 132.5* | *46 ± 29.1* | | | *1128*[3] | *260 ± 10.1* | | | *NA*[5] | *NA*[5] | | |

[1] The single largest straight-line distance recorded for each bird from the release pen.
[2] Calculated from the weekly average distance from the pen.
[3] Only one female bird survived to the shooting phase, thus SE not calculated.
[4] Only one male bird survived to the post-shooting phase, thus SE not calculated.
[5] No females survived to the post-shooting phase.

### Site F2

| Tagged | | Pre-shooting | | | | Shooting | | | | Post-shooting | | | |
|---|---|---|---|---|---|---|---|---|---|---|---|---|---|
| Bird ID | Sex | Max dist. from pen (m)[1] | Mean ± SE dist. from pen (m)[2] | Total weeks | Bird/tag outcome | Max dist. from pen (m)[1] | Mean ± SE dist. from pen (m)[2] | Total weeks | Bird/tag outcome | Max dist. from pen (m)[1] | Mean ± SE dist. from pen (m)[2] | Total weeks | Bird/tag outcome |
| 57182 | M | 59 | 4 ± 0.6 | 2 | Tag signal loss | NA | NA | 2 | Tag signal loss | NA | NA | 2 | Tag signal loss |
| 57183 | F | 117 | 3 ± 0.5 | 3 | Tag signal loss | NA | NA | 3 | Tag signal loss | NA | NA | 3 | Tag signal loss |
| 57184 | F | 74 | 3 ± 1.2 | 2 | Bird mortality | NA | NA | 2 | Bird mortality | NA | NA | 2 | Bird mortality |
| 57185 | F | 874 | 136 ± 3.2 | 14 | Survived | 1015 | 275 ± 3 | 22 | Tag signal loss | NA | NA | 22 | Tag signal loss |
| 57186 | M | 540 | 69 ± 2.8 | 9 | Tag signal loss | NA | NA | 9 | Tag signal loss | NA | NA | 9 | Tag signal loss |
| 57187 | M | 618 | 107 ± 2.6 | 14 | Survived | 1078 | 204 ± 3.8 | 25 | Shot | NA | NA | 25 | Shot |
| 57188 | M | 1312 | 399 ± 6.8 | 14 | Survived | 1246 | 445 ± 2.6 | 26 | Survived | 457 | 422 ± 1.8 | 27 | Tag signal loss |
| 57189 | M | 85 | 5 ± 0.8 | 2 | Tag signal loss | NA | NA | 2 | Tag signal loss | NA | NA | 2 | Tag signal loss |
| 57190 | F | 580 | 119 ± 3.2 | 14 | Survived | 746 | 264 ± 2.5 | 26 | Shot | NA | NA | 26 | Shot |
| 57191 | F | 528 | 24 ± 2 | 6 | Tag signal loss | NA | NA | 6 | Tag signal loss | NA | NA | 6 | Tag signal loss |

| | | | | | | | | | | | |
|---|---|---|---|---|---|---|---|---|---|---|---|
| *Mean ± SE* | *All* | *479 ± 129.6* | *87 ± 38.5* | | | *1021 ± 103.8* | *297 ± 51.7* | | | *457*[3] | *422 ± 1.8* |
| | *M* | *523 ± 228* | *117 ± 73.2* | | | *1162 ± 84* | *325 ± 120.5* | | | *457*[3] | *422 ± 1.8* |
| | *F* | *435 ± 150.6* | *57 ± 29.2* | | | *880 ± 134.1* | *270 ± 5.2* | | | *NA*[4] | *NA*[4] |

[1] The single largest straight-line distance recorded for each bird from the release pen.
[2] Calculated from the weekly average distance from the pen.
[3] Only one male bird survived to the post-shooting phase, thus SE not calculated.
[4] No females survived to the post-shooting phase.

### Site G

| Tagged | | Pre-shooting | | | | Shooting | | | | Post-shooting | | | |
|---|---|---|---|---|---|---|---|---|---|---|---|---|---|
| Bird ID | Sex | Max dist. from pen (m) [1] | Mean ± SE dist. from pen (m) [2] | Total weeks | Bird/tag outcome | Max dist. from pen (m) [1] | Mean ± SE dist. from pen (m) [2] | Total weeks | Bird/tag outcome | Max dist. from pen (m) [1] | Mean ± SE dist. from pen (m) [2] | Total weeks | Bird/tag outcome |
| 56561 | F | 44 | 5 ± 0.5 | 2 | Tag signal loss | NA | NA | 2 | Tag signal loss | NA | NA | 2 | Tag signal loss |
| 56562 | M | 894 | 40 ± 3.1 | 10 | Survived | 614 | 121 ± 3 | 15 | Tag signal loss | NA | NA | 15 | Tag signal loss |
| 57212 | M | 325 | 25 ± 1.1 | 10 | Survived | 955 | 182 ± 3.7 | 22 | Bird mortality | NA | NA | 22 | Bird mortality |
| 57213 | M | 1148 | 344 ± 10.6 | 10 | Survived | 1281 | 727 ± 3.4 | 19 | Tag signal loss | NA | NA | 19 | Tag signal loss |
| 57214 | F | 843 | 183 ± 8.4 | 10 | Survived | 1277 | 366 ± 4.9 | 25 | Survived | 392 | 72 ± 4.1 | 32 | Survived |
| 57215 | F | 442 | 76 ± 3.1 | 10 | Survived | 1431 | 405 ± 3.7 | 25 | Survived | 894 | 315 ± 8 | 30 | Tag signal loss |
| 57216 | M | 316 | 18 ± 1.1 | 10 | Survived | 222 | 29 ± 1.4 | 15 | Bird mortality | NA | NA | 15 | Bird mortality |
| 57217 | M | 491 | 34 ± 2.3 | 10 | Survived | 886 | 90 ± 2.7 | 25 | Survived | 422 | 129 ± 4.5 | 30 | Tag signal loss |
| 57218 | F | 1086 | 221 ± 7.4 | 10 | Survived | 1380 | 364 ± 4.3 | 25 | Survived | 1111 | 676 ± 4.7 | 32 | Survived |
| 57219 | F | 561 | 35 ± 2.2 | 10 | Survived | 1158 | 139 ± 3.8 | 25 | Survived | 773 | 85 ± 6 | 30 | Tag signal loss |
| 57220 | M | 409 | 18 ± 1.3 | 10 | Survived | 2229 | 489 ± 9 | 23 | Tag signal loss | NA | NA | 23 | Tag signal loss |
| 57221 | F | 2591 | 486 ± 17.1 | 10 | Survived | 2927 | 2490 ± 2.6 | 25 | Survived | 2621 | 2206 ± 15.7 | 28 | Tag signal loss |
| *Mean ± SE* | *All* | *762 ± 192.1* | *124 ± 44.7* | | | *1305 ± 223.2* | *491 ± 209.4* | | | *1036 ± 336.6* | *581 ± 338.2* | | |
| | *M* | *597 ± 140.3* | *80 ± 52.9* | | | *1031 ± 280.1* | *273 ± 112.1* | | | *422* [3] | *129 ± 4.5* | | |
| | *F* | *928 ± 362.9* | *168 ± 72.2* | | | *1635 ± 326.5* | *753 ± 436.9* | | | *1158 ± 383.8* | *671 ± 399.2* | | |

[1] The single largest straight-line distance recorded for each bird from the release pen.
[2] Calculated from the weekly average distance from the pen.
[3] Only one male survived to the post-shooting phase, thus SE not calculated.

### Site H

| Tagged | | Pre-shooting | | | | Shooting | | | | Post-shooting | | | |
|---|---|---|---|---|---|---|---|---|---|---|---|---|---|
| Bird ID | Sex | Max dist. from pen (m) [1] | Mean ± SE dist. from pen (m) [2] | Total weeks | Bird/tag outcome | Max dist. from pen (m) [1] | Mean ± SE dist. from pen (m) [2] | Total weeks | Bird/tag outcome | Max dist. from pen (m) [1] | Mean ± SE dist. from pen (m) [2] | Total weeks | Bird/tag outcome |
| 56565 | F | 355 | 130 ± 9.6 | 2 | Predated - fox | NA | NA | 2 | Predated - fox | NA | NA | 2 | Predated - fox |
| 57129 | F | 219 | 2 ± 0.4 | 9 | Survived | 638 | 69 ± 6.1 | 11 | Bird mortality | NA | NA | 11 | Bird mortality |
| 57192 | M | 723 | 82 ± 3.4 | 9 | Survived | 1184 | 286 ± 3.8 | 18 | Shot | NA | NA | 18 | Shot |
| 57196 | F | 435 | 23 ± 1.7 | 9 | Survived | 1172 | 145 ± 5.2 | 24 | Bird mortality | NA | NA | 24 | Bird mortality |
| 57198 | M | 824 | 52 ± 5.2 | 4 | Tag signal loss | NA | NA | 4 | Tag signal loss | NA | NA | 4 | Tag signal loss |
| 57199 | F | 652 | 49 ± 4.3 | 4 | Tag signal loss | NA | NA | 4 | Tag signal loss | NA | NA | 4 | Tag signal loss |
| 57200 | F | 1020 | 58 ± 3.5 | 9 | Survived | 506 | 61 ± 1.9 | 24 | Survived | 1125 | 68 ± 5.3 | 29 | Tag signal loss |
| 57201 | M | 1108 | 26 ± 3.6 | 9 | Survived | 576 | 154 ± 1.2 | 24 | Survived | 311 | 148 ± 0.9 | 31 | Survived |
| 57363 | M | 810 | 44 ± 3.2 | 9 | Survived | 1468 | 146 ± 5 | 24 | Survived | 631 | 114 ± 4 | 31 | Survived |
| 57364 | M | 541 | 18 ± 2 | 9 | Survived | 1460 | 356 ± 2.9 | 23 | Bird mortality | NA | NA | 23 | Bird mortality |
| 57365 | M | 634 | 75 ± 3.6 | 9 | Survived | 509 | 82 ± 7.8 | 11 | Tag mortality signal | NA | NA | 11 | Tag mortality signal |
| *Mean ± SE* | *All* | *666 ± 81.8* | *51 ± 10.8* | | | *939 ± 150* | *163 ± 37.6* | | | *689 ± 236.9* | *110 ± 23* | | |
| | *M* | *774 ± 80* | *50 ± 10.5* | | | *1039 ± 209.5* | *205 ± 50.3* | | | *471 ± 160.3* | *131 ± 17.2* | | |

| | | | | | | | | | | | |
|---|---|---|---|---|---|---|---|---|---|---|---|
| | *F* | *536 ± 139.8* | *52 ± 21.8* | | | *772 ± 203.6* | *92 ± 26.9* | | | *1125*[3] | *68 ± 5.3* |

[1] The single largest straight-line distance recorded for each bird from the release pen.
[2] Calculated from the weekly average distance from the pen.
[3] Only one female survived to the post-shooting phase, thus SE not calculated.

### Site I

| Tagged | | Pre-shooting | | | | Shooting | | | | Post-shooting | | | |
|---|---|---|---|---|---|---|---|---|---|---|---|---|---|
| Bird ID | Sex | Max dist. from pen (m)[1] | Mean ± SE dist. from pen (m)[2] | Total weeks | Bird/tag outcome | Max dist. from pen (m)[1] | Mean ± SE dist. from pen (m)[2] | Total weeks | Bird/tag outcome | Max dist. from pen (m)[1] | Mean ± SE dist. from pen (m)[2] | Total weeks | Bird/tag outcome |
| 56565* | M | 28 | 7 ± 1.4 | 1 | Tag signal loss | NA | NA | 1 | Tag signal loss | NA | NA | 1 | Tag signal loss |
| 57137* | M | 1286 | 395 ± 9.8 | 10 | Survived | 1699 | 1146 ± 4.3 | 22 | Survived | 1590 | 895 ± 8 | 29 | Survived |
| 57164* | F | 163 | 82 ± 6.6 | 1 | Tag mortality signal | NA | NA | 1 | Tag mortality signal | NA | NA | 1 | Tag mortality signal |
| 57171* | M | 617 | 58 ± 9.9 | 3 | Tag signal loss | NA | NA | 3 | Tag signal loss | NA | NA | 3 | Tag signal loss |
| 57178* | M | 436 | 21 ± 3 | 4 | Predated - fox | NA | NA | 4 | Predated - fox | NA | NA | 4 | Predated - fox |
| 57209* | F | 209 | 10 ± 1.7 | 3 | Tag signal loss | NA | NA | 3 | Tag signal loss | NA | NA | 3 | Tag signal loss |
| *Mean ± SE* | *All* | *457 ± 186.7* | *96 ± 61.1* | | | *1699*[3] | *1146 ± 4.3* | | | *1590*[3] | *895 ± 8* | | |
| | *M* | *592 ± 262.2* | *120 ± 92.2* | | | *1699*[3] | *1146 ± 4.3* | | | *1590*[3] | *895 ± 8* | | |
| | *F* | *186 ± 22.6* | *46 ± 36* | | | *NA*[4] | *NA*[4] | | | *NA*[4] | *NA*[4] | | |

[1] The single largest straight-line distance recorded for each bird from the release pen.
[2] Calculated from the weekly average distance from the pen.
[3] Only one male bird survived to the shooting phase and post-shooting phase, thus SE not calculated.
[4] No females survived to the shooting phase.
*Tags already used on previous birds that were found dead and reapplied to new birds.

**Section 2: Summary ranging profile for tagged birds, indicating the percentage of total fixes (i.e., total time) within specified distances from the release pen, during the '*pre-shooting phase*', '*shooting phase*' and '*post-shooting*' phase. Birds that spent >50% of time beyond 500m and 1,000m and time outside estate boundaries are noted.**

### Site A

| Pre-shooting | Time spent (number of fixes) (%) | | | | | | | | | | Mean ± SE |
|---|---|---|---|---|---|---|---|---|---|---|---|
| | 57142 | 57143 | 57144 | 57145 | 57146 | 57147 | 57148 | 57149 | 57150 | 57151 | |
| Total fixes | 1145 | 1722 | 1487 | 1708 | 1647 | 1685 | 1699 | 1666 | 1684 | 1655 | |
| <500m | 93.9 | 57.4 | 59.6 | 98.5 | 92.5 | 62.9 | 43.9 | 93.6 | 77.5 | 89.4 | 76.9 ± 6.1 |
| >500m<700m | 5.9 | 33.6 | 13.4 | 1.5 | 6.9 | 28.7 | 37.8 | 4.9 | 12.8 | 7.9 | 15.3 ± 4.1 |
| >700m<1000m | 0.2 | 8.8 | 15.4 | 0.0 | 0.6 | 7.8 | 17.3 | 1.3 | 9.6 | 2.7 | 6.4 ± 2.0 |
| >1000m<1500m | 0.0 | 0.2 | 11.6 | 0.0 | 0.0 | 0.6 | 0.9 | 0.2 | 0.1 | 0.0 | 1.4 ± 1.1 |
| >1500m<2000m | 0.0 | 0.0 | 0.0 | 0.0 | 0.0 | 0.0 | 0.0 | 0.0 | 0.0 | 0.0 | 0.0 |
| >2000m | 0.0 | 0.0 | 0.0 | 0.0 | 0.0 | 0.0 | 0.0 | 0.0 | 0.0 | 0.0 | 0.0 |
| ≥50% fixes >500m | - | - | - | - | - | - | Yes | - | - | - | - |

| Shooting | Time spent (number of fixes) (%) | | | | | | | | | | Mean ± SE |
|---|---|---|---|---|---|---|---|---|---|---|---|
| | 57142 | 57143 | 57144 | 57145 | 57146 | 57147 | 57148 | 57149 | 57150 | 57151 | |
| Total fixes | NA | 2407 | 2358 | 2517 | 521 | NA | 640 | 982 | 635 | 1021 | |
| <500m | NA | 19.1 | 10.6 | 98.4 | 82.5 | NA | 16.7 | 90.6 | 25.4 | 49.3 | 49.1 ± 12.9 |
| >500m<700m | NA | 8.5 | 1.6 | 0.9 | 16.1 | NA | 41.6 | 5.6 | 52.8 | 20.8 | 18.5 ± 6.8 |
| >700m<1000m | NA | 6.8 | 5.9 | 0.1 | 1.2 | NA | 35.5 | 0.7 | 20.8 | 17.0 | 11.0 ± 4.4 |
| >1000m<1500m | NA | 2.2 | 9.0 | 0.6 | 0.2 | NA | 6.1 | 0.6 | 1.1 | 12.9 | 4.1 ± 1.7 |
| >1500m<2000m | NA | 63.1 | 53.4 | 0.0 | 0.0 | NA | 0.2 | 0.4 | 0.0 | 0.0 | 14.6 ± 9.6 |
| >2000m | NA | 0.2 | 19.5 | 0.0 | 0.0 | NA | 0.0 | 2.0 | 0.0 | 0.0 | 2.7 ± 2.4 |
| ≥50% fixes >500m | - | Yes | Yes | - | - | - | Yes | - | Yes | Yes | - |
| ≥50% fixes >1000m | - | Yes | Yes | - | - | - | - | - | - | - | - |

| Post-shooting | Time spent (number of fixes) (%) | | | | | | | | | | Mean ± SE |
|---|---|---|---|---|---|---|---|---|---|---|---|
| | 57142 | 57143 | 57144 | 57145 | 57146 | 57147 | 57148 | 57149 | 57150 | 57151 | |
| Total fixes | NA | 847 | 871 | 1241 | NA | NA | NA | NA | NA | NA | |
| <500m | NA | 0.0 | 0.0 | 99.9 | NA | NA | NA | NA | NA | NA | 33.3 ± 33.3 |
| >500m<700m | NA | 0.0 | 0.0 | 0.1 | NA | NA | NA | NA | NA | NA | 0.03 ± 0.03 |
| >700m<1000m | NA | 0.0 | 0.0 | 0.0 | NA | NA | NA | NA | NA | NA | 0.0 |
| >1000m<1500m | NA | 2.6 | 2.2 | 0.0 | NA | NA | NA | NA | NA | NA | 1.6 ± 0.8 |
| >1500m<2000m | NA | 96.7 | 76.7 | 0.0 | NA | NA | NA | NA | NA | NA | 57.8 ± 29.5 |
| >2000m | NA | 0.7 | 21.1 | 0.0 | NA | NA | NA | NA | NA | NA | 7.3 ± 6.9 |
| ≥50% fixes >500m | - | Yes | Yes | - | - | - | - | - | - | - | - |
| ≥50% fixes >1000m | - | Yes | Yes | - | - | - | - | - | - | - | - |

*% fixes outside estate not calculated for this site as estate boundary was not available.

### Site B

| Pre-shooting | Time spent (number of fixes) (%) | | | | | | | | | | Mean ± SE |
|---|---|---|---|---|---|---|---|---|---|---|---|
| | 57122 | 57123 | 57124 | 57125 | 57126 | 57127 | 57128 | 57129 | 57130 | 57131 | |
| Total fixes | 1886 | 1726 | 1633 | 1648 | 1714 | 1710 | 1792 | 233 | 1864 | 1736 | |
| <500m | 92.4 | 85.7 | 92.2 | 91.5 | 95.6 | 83.8 | 96.8 | 100 | 72.9 | 59.9 | 87.1 ± 3.9 |
| >500m<700m | 7.2 | 7.1 | 7.0 | 2.4 | 4.3 | 4.7 | 2.3 | 0.0 | 9.1 | 4.2 | 4.8 ± 0.9 |
| >700m<1000m | 0.4 | 4.1 | 0.7 | 5.0 | 0.1 | 1.8 | 0.7 | 0.0 | 6.3 | 6.0 | 2.5 ± 0.8 |
| >1000m<1500m | 0.0 | 2.9 | 0.1 | 1.0 | 0.0 | 2.0 | 0.2 | 0.0 | 7.6 | 6.3 | 2.0 ± 0.9 |
| >1500m<2000m | 0.0 | 0.2 | 0.0 | 0.0 | 0.0 | 1.8 | 0.0 | 0.0 | 2.7 | 15.5 | 2.0 ± 1.5 |
| >2000m | 0.0 | 0.0 | 0.0 | 0.0 | 0.0 | 5.8 | 0.0 | 0.0 | 1.3 | 8.1 | 1.5 ± 0.9 |
| % fixes outside estate | 0.0 | 1.8 | 0.0 | 0.0 | 0.0 | 7.8 | 0.0 | 0.0 | 0.0 | 22.5 | 3.2 ± 2.3 |

| Shooting | Time spent (number of fixes) (%) | | | | | | | | | | Mean ± SE |
|---|---|---|---|---|---|---|---|---|---|---|---|
| | 57122 | 57123 | 57124 | 57125 | 57126 | 57127 | 57128 | 57129 | 57130 | 57131 | |
| Total fixes | 1658 | 2453 | 2177 | 865 | 2282 | 2330 | 2278 | NA | 573 | 2428 | |
| <500m | 88.8 | 16.4 | 84.8 | 98.8 | 9.6 | 0.0 | 93.0 | NA | 8.9 | 0.0 | 44.5 ± 15.0 |
| >500m<700m | 10.4 | 0.8 | 13.2 | 1.2 | 3.5 | 0.0 | 3.7 | NA | 7.3 | 0.0 | 4.5 ± 1.6 |
| >700m<1000m | 0.8 | 0.3 | 1.7 | 0.0 | 13.2 | 0.0 | 1.1 | NA | 5.1 | 0.0 | 2.5 ± 1.4 |
| >1000m<1500m | 0.0 | 0.5 | 0.2 | 0.0 | 34.6 | 0.0 | 2.1 | NA | 61.3 | 0.0 | 11.0 ± 7.3 |
| >1500m<2000m | 0.0 | 1.3 | 0.0 | 0.0 | 22.8 | 0.0 | 0.1 | NA | 12.7 | 0.0 | 4.1 ± 2.7 |
| >2000m | 0.0 | 80.6 | 0.0 | 0.0 | 16.3 | 100 | 0.0 | NA | 4.7 | 100 | 33.5 ± 15.2 |
| ≥50% fixes >500m | - | Yes | - | - | Yes | Yes | - | - | Yes | Yes | - |
| ≥50% fixes >1000m | - | Yes | - | - | Yes | Yes | - | - | Yes | Yes | - |
| % fixes outside estate | 0.0 | 0.7 | 0.1 | 0.0 | 4.4 | 100 | 0.0 | NA | 0.0 | 0.0 | 11.7 ± 11.1 |

| Post-shooting | Time spent (number of fixes) (%) | | | | | | | | | | Mean ± SE |
|---|---|---|---|---|---|---|---|---|---|---|---|
| | 57122 | 57123 | 57124 | 57125 | 57126 | 57127 | 57128 | 57129 | 57130 | 57131 | |
| Total fixes | NA | 1223 | 383 | NA | 92 | 853 | NA | NA | NA | 400 | |
| <500m | NA | 7.9 | 47.8 | NA | 18.5 | 0.0 | NA | NA | NA | 0.0 | 14.8 ± 8.9 |
| >500m<700m | NA | 0.1 | 52.2 | NA | 18.5 | 0.0 | NA | NA | NA | 0.0 | 14.2 ± 10.2 |
| >700m<1000m | NA | 2.5 | 0.0 | NA | 39.1 | 0.0 | NA | NA | NA | 0.0 | 8.3 ± 7.7 |
| >1000m<1500m | NA | 1.3 | 0.0 | NA | 22.8 | 0.0 | NA | NA | NA | 0.0 | 4.8 ± 4.5 |
| >1500m<2000m | NA | 0.4 | 0.0 | NA | 1.1 | 0.0 | NA | NA | NA | 0.0 | 0.3 ± 0.2 |
| >2000m | NA | 87.9 | 0.0 | NA | 0.0 | 100 | NA | NA | NA | 100 | 57.6 ± 23.6 |
| ≥50% fixes >500m | - | Yes | Yes | - | Yes | Yes | - | - | - | Yes | - |
| ≥50% fixes >1000m | - | Yes | - | - | - | Yes | - | - | - | Yes | - |
| % fixes outside estate | NA | 0.2 | 0.0 | NA | 0.0 | 100 | NA | NA | NA | 0.0 | 20.0 ± 20.0 |

### Site C1

| Pre-shooting | Time spent (number of fixes) (%) | | | | | | | | | | Mean ± SE |
|---|---|---|---|---|---|---|---|---|---|---|---|
| | 57152 | 57153 | 57154 | 57155 | 57156 | 57157 | 57158 | 57159 | 57160 | 57161 | |
| Total fixes | 2269 | 2232 | 111 | 227 | 642 | 102 | 362 | 337 | 2226 | 2172 | |
| <500m | 98.8 | 42.0 | 100 | 100 | 80.8 | 100 | 100 | 100 | 78.3 | 90.3 | 89 ± 5.9 |
| >500m<700m | 1.0 | 0.8 | 0.0 | 0.0 | 19.2 | 0.0 | 0.0 | 0.0 | 19.4 | 7.0 | 4.7 ± 2.5 |
| >700m<1000m | 0.2 | 26.7 | 0.0 | 0.0 | 0.0 | 0.0 | 0.0 | 0.0 | 2.2 | 1.7 | 3.1 ± 2.6 |
| >1000m<1500m | 0.0 | 30.0 | 0.0 | 0.0 | 0.0 | 0.0 | 0.0 | 0.0 | 0.0 | 1.0 | 3.1 ± 3.0 |
| >1500m<2000m | 0.0 | 0.5 | 0.0 | 0.0 | 0.0 | 0.0 | 0.0 | 0.0 | 0.0 | 0.0 | 0.1 ± 0.1 |
| >2000m | 0.0 | 0.0 | 0.0 | 0.0 | 0.0 | 0.0 | 0.0 | 0.0 | 0.0 | 0.0 | 0.0 |
| ≥50% fixes >500m | - | Yes | - | - | - | - | - | - | - | - | - |
| % fixes outside estate | 0.0 | 0.0 | 0.0 | 0.0 | 0.0 | 0.0 | 0.0 | 0.0 | 0.0 | 0.0 | 0.0 |

| Shooting | Time spent (number of fixes) (%) | | | | | | | | | | Mean ± SE |
|---|---|---|---|---|---|---|---|---|---|---|---|
| | 57152 | 57153 | 57154 | 57155 | 57156 | 57157 | 57158 | 57159 | 57160 | 57161 | |
| Total fixes | 1923 | 1938 | NA | NA | NA | NA | NA | NA | 1372 | 1633 | |
| <500m | 67.9 | 0.2 | NA | NA | NA | NA | NA | NA | 87.5 | 78.1 | 58.4 ± 19.8 |
| >500m<700m | 30.0 | 0.2 | NA | NA | NA | NA | NA | NA | 10.9 | 10.9 | 13.0 ± 6.2 |
| >700m<1000m | 1.9 | 26.4 | NA | NA | NA | NA | NA | NA | 1.5 | 5.2 | 8.8 ± 5.9 |
| >1000m<1500m | 0.3 | 73.2 | NA | NA | NA | NA | NA | NA | 0.1 | 5.7 | 19.8 ± 17.8 |
| >1500m<2000m | 0.0 | 0.0 | NA | NA | NA | NA | NA | NA | 0.0 | 0.1 | 0.03 ± 0.03 |
| >2000m | 0.0 | 0.0 | NA | NA | NA | NA | NA | NA | 0.0 | 0.0 | 0.0 |
| ≥50% fixes >500m | - | Yes | - | - | - | - | - | - | - | - | - |
| ≥50% fixes >1000m | - | Yes | - | - | - | - | - | - | - | - | - |
| % fixes outside estate | 0.0 | 0.0 | NA | NA | NA | NA | NA | NA | 0.0 | 0.0 | 0.0 |

| Post-shooting | Time spent (number of fixes) (%) | | | | | | | | | | Mean ± SE |
|---|---|---|---|---|---|---|---|---|---|---|---|
| | 57152 | 57153 | 57154 | 57155 | 57156 | 57157 | 57158 | 57159 | 57160 | 57161 | |
| Total fixes | 574 | 1192 | NA | NA | NA | NA | NA | NA | NA | NA | |
| <500m | 90.8 | 0.0 | NA | NA | NA | NA | NA | NA | NA | NA | 45.4 ± 45.4 |
| >500m<700m | 9.2 | 0.1 | NA | NA | NA | NA | NA | NA | NA | NA | 4.7 ± 4.6 |
| >700m<1000m | 0.0 | 43.5 | NA | NA | NA | NA | NA | NA | NA | NA | 21.8 ± 21.8 |
| >1000m<1500m | 0.0 | 56.5 | NA | NA | NA | NA | NA | NA | NA | NA | 28.3 ± 28.3 |
| >1500m<2000m | 0.0 | 0.0 | NA | NA | NA | NA | NA | NA | NA | NA | 0.0 |
| >2000m | 0.0 | 0.0 | NA | NA | NA | NA | NA | NA | NA | NA | 0.0 |
| ≥50% fixes >500m | - | Yes | - | - | - | - | - | - | - | - | - |
| ≥50% fixes >1000m | - | Yes | - | - | - | - | - | - | - | - | - |
| % fixes outside estate | 0.0 | 0.0 | NA | NA | NA | NA | NA | NA | NA | NA | 0.0 |

### Site C2

| Pre-shooting | 57162 | 57163 | 57164 | 57165 | 57166 | 57167 | 57169 | 57170 | 57171 | 57137* | 57165b* | Mean ± SE |
|---|---|---|---|---|---|---|---|---|---|---|---|---|
| Total fixes | 2074 | 1995 | 88 | 379 | 2092 | 2154 | 2071 | 1979 | 399 | 88 | 1398 | |
| <500m | 28.7 | 93.1 | 98.9 | 100 | 100 | 100 | 100 | 38.7 | 100 | 100 | 84.3 | 85.8 ± 7.9 |
| >500m<700m | 42.2 | 4.9 | 0.0 | 0.0 | 0.0 | 0.0 | 0.0 | 11.3 | 0.0 | 0.0 | 13.0 | 6.5 ± 3.9 |
| >700m<1000m | 18.2 | 1.1 | 0.0 | 0.0 | 0.0 | 0.0 | 0.0 | 14.7 | 0.0 | 0.0 | 1.7 | 3.2 ± 2.0 |
| >1000m<1500m | 10.8 | 1.0 | 0.0 | 0.0 | 0.0 | 0.0 | 0.0 | 35.0 | 0.0 | 0.0 | 0.9 | 4.3 ± 3.2 |
| >1500m<2000m | 0.0 | 0.0 | 1.1 | 0.0 | 0.0 | 0.0 | 0.0 | 0.3 | 0.0 | 0.0 | 0.0 | 0.1 ± 0.1 |
| >2000m | 0.0 | 0.0 | 0 | 0.0 | 0.0 | 0.0 | 0.0 | 0.0 | 0.0 | 0.0 | 0.0 | 0.0 |
| ≥50% fixes >500m | Yes | - | - | - | - | - | - | Yes | - | - | - | - |
| % fixes outside estate | 0.1 | 0.0 | 0.0 | 0.0 | 0.0 | 0.0 | 0.0 | 0.0 | 0.0 | 0.0 | 0.0 | 0.01 ± 0.01 |

| Shooting | 57162 | 57163 | 57164 | 57165 | 57166 | 57167 | 57169 | 57170 | 57171 | 57137* | 57165b* | Mean ± SE |
|---|---|---|---|---|---|---|---|---|---|---|---|---|
| Total fixes | 13 | 17 | NA | NA | 1360 | 1962 | 279 | 1252 | NA | NA | NA | |
| <500m | 0 | 17.6 | NA | NA | 100 | 85.6 | 100 | 0.0 | NA | NA | NA | 50.5 ± 20.3 |
| >500m<700m | 76.9 | 76.5 | NA | NA | 0.0 | 13.4 | 0.0 | 0.0 | NA | NA | NA | 27.8 ± 15.6 |
| >700m<1000m | 23.1 | 5.9 | NA | NA | 0.0 | 0.2 | 0.0 | 0.0 | NA | NA | NA | 4.9 ± 3.8 |
| >1000m<1500m | 0.0 | 0.0 | NA | NA | 0.0 | 0.8 | 0.0 | 97.1 | NA | NA | NA | 16.3 ± 16.2 |
| >1500m<2000m | 0.0 | 0.0 | NA | NA | 0.0 | 0.0 | 0.0 | 2.9 | NA | NA | NA | 0.5 ± 0.5 |
| >2000m | 0.0 | 0.0 | NA | NA | 0.0 | 0.0 | 0.0 | 0.0 | NA | NA | NA | 0.0 |
| ≥50% fixes >500m | Yes | Yes | - | - | - | - | - | Yes | - | - | - | - |
| ≥50% fixes >1000m | - | - | - | - | - | - | - | Yes | - | - | - | - |
| % fixes outside estate | 0.0 | 0.0 | NA | NA | 0.0 | 0.0 | 0.0 | 0.0 | NA | NA | NA | 0.0 |

| Post-shooting | 57162 | 57163 | 57164 | 57165 | 57166 | 57167 | 57169 | 57170 | 57171 | 57137* | 57165b* | Mean ± SE |
|---|---|---|---|---|---|---|---|---|---|---|---|---|
| Total fixes | NA | NA | NA | NA | NA | 1185 | NA | NA | NA | NA | NA | |
| <500m | NA | NA | NA | NA | NA | 97.0 | NA | NA | NA | NA | NA | 97.0 |
| >500m<700m | NA | NA | NA | NA | NA | 3.0 | NA | NA | NA | NA | NA | 3.0 |
| >700m<1000m | NA | NA | NA | NA | NA | 0.0 | NA | NA | NA | NA | NA | 0.0 |
| >1000m<1500m | NA | NA | NA | NA | NA | 0.0 | NA | NA | NA | NA | NA | 0.0 |
| >1500m<2000m | NA | NA | NA | NA | NA | 0.0 | NA | NA | NA | NA | NA | 0.0 |
| >2000m | NA | NA | NA | NA | NA | 0.0 | NA | NA | NA | NA | NA | 0.0 |
| % fixes outside estate | NA | NA | NA | NA | NA | 0.0 | NA | NA | NA | NA | NA | 0.0 |

### Site D

| Pre-shooting | Time spent (number of fixes) (%) | | | | | | | | | | Mean ± SE |
|---|---|---|---|---|---|---|---|---|---|---|---|
| | 57132 | 57133 | 57134 | 57135 | 57136 | 57137 | 57138 | 57139 | 57140 | 57141 | |
| Total fixes | 1713 | 1730 | 1276 | 1767 | 1749 | 391 | 1815 | 1788 | 1753 | 1718 | |
| <500m | 98.5 | 66.0 | 99.3 | 48.8 | 90.4 | 100 | 35.4 | 99.6 | 42.6 | 98.0 | 77.9 ± 8.5 |
| >500m<700m | 1.5 | 30.9 | 0.6 | 0.2 | 3.4 | 0.0 | 0.2 | 0.4 | 0.1 | 1.6 | 3.9 ± 3.0 |
| >700m<1000m | 0.0 | 2.7 | 0.1 | 13.4 | 3.7 | 0.0 | 0.7 | 0.0 | 1.5 | 0.3 | 2.2 ± 1.3 |
| >1000m<1500m | 0.0 | 0.4 | 0.0 | 37.6 | 2.5 | 0.0 | 57.1 | 0.0 | 50.2 | 0.0 | 14.8 ± 7.5 |
| >1500m<2000m | 0.0 | 0.0 | 0.0 | 0.0 | 0.1 | 0.0 | 5.2 | 0.0 | 1.7 | 0.0 | 0.7 ± 0.5 |
| >2000m | 0.0 | 0.0 | 0.0 | 0.0 | 0.0 | 0.0 | 1.3 | 0.0 | 3.9 | 0.0 | 0.5 ± 0.4 |
| ≥50% fixes >500m | - | - | - | Yes | - | - | Yes | - | Yes | - | - |
| ≥50% fixes >1000m | - | - | - | - | - | - | Yes | - | Yes | - | - |
| % fixes outside estate | 0.0 | 0.0 | 0.0 | 0.0 | 0.0 | 0.0 | 0.0 | 0.0 | 0.0 | 0.0 | 0.0 |

| Shooting | Time spent (number of fixes) (%) | | | | | | | | | | Mean ± SE |
|---|---|---|---|---|---|---|---|---|---|---|---|
| | 57132 | 57133 | 57134 | 57135 | 57136 | 57137 | 57138 | 57139 | 57140 | 57141 | |
| Total fixes | 795 | 443 | NA | 2314 | 2140 | NA | 884 | 1916 | 2290 | NA | |
| <500m | 93.5 | 89.2 | NA | 0.0 | 22.4 | NA | 17.2 | 4.4 | 0.0 | NA | 32.4 ± 15.6 |
| >500m<700m | 6.0 | 10.8 | NA | 0.1 | 2.8 | NA | 0.8 | 34.2 | 0.0 | NA | 7.8 ± 4.6 |
| >700m<1000m | 0.5 | 0.0 | NA | 7.2 | 14.0 | NA | 0.9 | 60.3 | 0.0 | NA | 11.8 ± 8.3 |
| >1000m<1500m | 0.0 | 0.0 | NA | 92.7 | 15.9 | NA | 25.9 | 1.0 | 0.0 | NA | 19.4 ± 12.8 |
| >1500m<2000m | 0.0 | 0.0 | NA | 0.0 | 33.6 | NA | 1.9 | 0.0 | 1.5 | NA | 5.3 ± 4.7 |
| >2000m | 0.0 | 0.0 | NA | 0.0 | 11.3 | NA | 53.3 | 0.0 | 98.5 | NA | 23.3 ± 14.5 |
| ≥50% fixes >500m | - | - | - | Yes | Yes | - | Yes | Yes | Yes | - | - |
| ≥50% fixes >1000m | - | - | - | Yes | Yes | - | Yes | - | Yes | - | - |
| % fixes outside estate | 0.0 | 0.0 | NA | 0.0 | 1.8 | NA | 45.7 | 0.0 | 0.0 | NA | 6.8 ± 6.5 |

| Post-shooting | Time spent (number of fixes) (%) | | | | | | | | | | Mean ± SE |
|---|---|---|---|---|---|---|---|---|---|---|---|
| | 57132 | 57133 | 57134 | 57135 | 57136 | 57137 | 57138 | 57139 | 57140 | 57141 | |
| Total fixes | NA | NA | NA | 1238 | 565 | NA | NA | NA | 1210 | NA | |
| <500m | NA | NA | NA | 0.0 | 0.0 | NA | NA | NA | 0.0 | NA | 0.0 |
| >500m<700m | NA | NA | NA | 0.0 | 0.0 | NA | NA | NA | 0.0 | NA | 0.0 |
| >700m<1000m | NA | NA | NA | 12.3 | 0.0 | NA | NA | NA | 0.0 | NA | 4.1 ± 4.1 |
| >1000m<1500m | NA | NA | NA | 87.7 | 4.1 | NA | NA | NA | 0.0 | NA | 30.6 ± 28.6 |
| >1500m<2000m | NA | NA | NA | 0.0 | 93.3 | NA | NA | NA | 0.7 | NA | 31.3 ± 31.0 |
| >2000m | NA | NA | NA | 0.0 | 2.7 | NA | NA | NA | 99.3 | NA | 34.0 ± 32.7 |
| ≥50% fixes >500m | - | - | - | Yes | Yes | - | - | - | Yes | - | - |
| ≥50% fixes >1000m | - | - | - | Yes | Yes | - | - | - | Yes | - | - |
| % fixes outside estate | NA | NA | NA | 0.0 | 0.0 | NA | NA | NA | 0.0 | NA | 0.0 |

### Site E

| Pre-shooting | Time spent (number of fixes) (%) | | | | | | | | | | Mean ± SE |
|---|---|---|---|---|---|---|---|---|---|---|---|
| | 57202 | 57203 | 57204 | 57205 | 57206 | 57207 | 57208 | 57209 | 57210 | 57211 | |
| Total fixes | 781 | 755 | 734 | 632 | 702 | 645 | 675 | 147 | 685 | 756 | |
| <500m | 100 | 100 | 100 | 100 | 99.7 | 100 | 100 | 99.3 | 93.0 | 99.7 | 99.2 ± 0.7 |
| >500m<700m | 0.0 | 0.0 | 0.0 | 0.0 | 0.3 | 0.0 | 0.0 | 0.7 | 6.7 | 0.1 | 0.8 ± 0.7 |
| >700m<1000m | 0.0 | 0.0 | 0.0 | 0.0 | 0.0 | 0.0 | 0.0 | 0.0 | 0.3 | 0.1 | 0.04 ± 0.03 |
| >1000m<1500m | 0.0 | 0.0 | 0.0 | 0.0 | 0.0 | 0.0 | 0.0 | 0.0 | 0.0 | 0.0 | 0.0 |
| >1500m<2000m | 0.0 | 0.0 | 0.0 | 0.0 | 0.0 | 0.0 | 0.0 | 0.0 | 0.0 | 0.0 | 0.0 |
| >2000m | 0.0 | 0.0 | 0.0 | 0.0 | 0.0 | 0.0 | 0.0 | 0.0 | 0.0 | 0.0 | 0.0 |
| % fixes outside estate | 0.1 | 0.0 | 0.0 | 0.0 | 0.0 | 0.0 | 0.0 | 0.0 | 0.1 | 0.3 | 0.1 ± 0.03 |

| Shooting | Time spent (number of fixes) (%) | | | | | | | | | | Mean ± SE |
|---|---|---|---|---|---|---|---|---|---|---|---|
| | 57202 | 57203 | 57204 | 57205 | 57206 | 57207 | 57208 | 57209 | 57210 | 57211 | |
| Total fixes | 209 | 2740 | 2571 | 1867 | 842 | NA | 595 | NA | 2175 | 2360 | |
| <500m | 100 | 93.5 | 82.2 | 21.7 | 96.8 | NA | 99.7 | NA | 38.1 | 18.6 | 68.8 ± 12.8 |
| >500m<700m | 0.0 | 6.3 | 8.6 | 0.2 | 2.7 | NA | 0.3 | NA | 42.3 | 0.5 | 7.6 ± 5.1 |
| >700m<1000m | 0.0 | 0.3 | 2.5 | 0.2 | 0.1 | NA | 0.0 | NA | 15.2 | 7.8 | 3.3 ± 2.0 |
| >1000m<1500m | 0.0 | 0.0 | 5.8 | 0.2 | 0.4 | NA | 0.0 | NA | 4.5 | 48.8 | 7.5 ± 6.0 |
| >1500m<2000m | 0.0 | 0.0 | 0.9 | 1.0 | 0.0 | NA | 0.0 | NA | 0.0 | 17.5 | 2.4 ± 2.2 |
| >2000m | 0.0 | 0.0 | 0.0 | 76.6 | 0.0 | NA | 0.0 | NA | 0.0 | 6.8 | 10.4 ± 9.5 |
| ≥50% fixes >500m | - | - | - | Yes | - | - | - | - | Yes | Yes | - |
| ≥50% fixes >1000m | - | - | - | Yes | - | - | - | - | - | Yes | - |
| % fixes outside estate | 0.0 | 0.1 | 7.4 | 77.8 | 0.6 | NA | 0.0 | NA | 0.1 | 80.9 | 20.9 ± 12.8 |

| Post-shooting | Time spent (number of fixes) (%) | | | | | | | | | | Mean ± SE |
|---|---|---|---|---|---|---|---|---|---|---|---|
| | 57202 | 57203 | 57204 | 57205 | 57206 | 57207 | 57208 | 57209 | 57210 | 57211 | |
| Total fixes | NA | 257 | NA | NA | NA | NA | NA | NA | NA | NA | |
| <500m | NA | 78.6 | NA | NA | NA | NA | NA | NA | NA | NA | 78.6 |
| >500m<700m | NA | 21.0 | NA | NA | NA | NA | NA | NA | NA | NA | 21.0 |
| >700m<1000m | NA | 0.4 | NA | NA | NA | NA | NA | NA | NA | NA | 0.4 |
| >1000m<1500m | NA | 0.0 | NA | NA | NA | NA | NA | NA | NA | NA | 0.0 |
| >1500m<2000m | NA | 0.0 | NA | NA | NA | NA | NA | NA | NA | NA | 0.0 |
| >2000m | NA | 0.0 | NA | NA | NA | NA | NA | NA | NA | NA | 0.0 |
| % fixes outside estate | NA | 0.0 | NA | NA | NA | NA | NA | NA | NA | NA | 0.0 |

### Site F1

| Pre-shooting | Time spent (number of fixes) (%) | | | | | | | | | | Mean ± SE |
|---|---|---|---|---|---|---|---|---|---|---|---|
| | 57172 | 57173 | 57174 | 57175 | 57176 | 57177 | 57178 | 57179 | 57180 | 57181 | |
| Total fixes | 1784 | 1776 | 242 | 1814 | 1837 | 290 | 137 | 1749 | 150 | 7 | |
| <500m | 99.3 | 99.8 | 100 | 98.8 | 94.0 | 100 | 100 | 96.1 | 100 | 100 | 98.8 ± 0.7 |
| >500m<700m | 0.5 | 0.2 | 0.0 | 1.0 | 0.7 | 0.0 | 0.0 | 2.8 | 0.0 | 0.0 | 0.5 ± 0.3 |
| >700m<1000m | 0.2 | 0.0 | 0.0 | 0.1 | 0.4 | 0.0 | 0.0 | 0.2 | 0.0 | 0.0 | 0.1 ± 0.04 |
| >1000m<1500m | 0.0 | 0.0 | 0.0 | 0.0 | 4.0 | 0.0 | 0.0 | 0.9 | 0.0 | 0.0 | 0.5 ± 0.4 |
| >1500m<2000m | 0.0 | 0.0 | 0.0 | 0.0 | 1.0 | 0.0 | 0.0 | 0.0 | 0.0 | 0.0 | 0.1 ± 0.1 |
| >2000m | 0.0 | 0.0 | 0.0 | 0.0 | 0.0 | 0.0 | 0.0 | 0.0 | 0.0 | 0.0 | 0.0 |
| % fixes outside estate | 0.0 | 0.0 | 0.0 | 0.0 | 0.0 | 0.0 | 0.0 | 0.0 | 0.0 | 0.0 | 0.0 |

| Shooting | Time spent (number of fixes) (%) | | | | | | | | | | Mean ± SE |
|---|---|---|---|---|---|---|---|---|---|---|---|
| | 57172 | 57173 | 57174 | 57175 | 57176 | 57177 | 57178 | 57179 | 57180 | 57181 | |
| Total fixes | 1595 | 1018 | NA | 449 | 1724 | NA | NA | 23 | NA | NA | |
| <500m | 98.7 | 98.4 | NA | 94.7 | 22.0 | NA | NA | 91.3 | NA | NA | 81.0 ± 14.8 |
| >500m<700m | 1.1 | 1.5 | NA | 0.9 | 8.0 | NA | NA | 8.7 | NA | NA | 4.0 ± 1.8 |
| >700m<1000m | 0.2 | 0.1 | NA | 1.1 | 1.3 | NA | NA | 0.0 | NA | NA | 0.5 ± 0.3 |
| >1000m<1500m | 0.0 | 0.0 | NA | 3.3 | 66.0 | NA | NA | 0.0 | NA | NA | 13.9 ± 13.1 |
| >1500m<2000m | 0.0 | 0.0 | NA | 0.0 | 2.7 | NA | NA | 0.0 | NA | NA | 0.5 ± 0.5 |
| >2000m | 0.0 | 0.0 | NA | 0.0 | 0.0 | NA | NA | 0.0 | NA | NA | 0.0 |
| ≥50% fixes >500m | - | - | - | - | Yes | - | - | - | - | - | - |
| ≥50% fixes >1000m | - | - | - | - | Yes | - | - | - | - | - | - |
| % fixes outside estate | 0.0 | 0.0 | NA | 0.0 | 0.0 | NA | NA | 0.0 | NA | NA | 0.0 |

| Post-shooting | Time spent (number of fixes) (%) | | | | | | | | | | Mean ± SE |
|---|---|---|---|---|---|---|---|---|---|---|---|
| | 57172 | 57173 | 57174 | 57175 | 57176 | 57177 | 57178 | 57179 | 57180 | 57181 | |
| Total fixes | NA | NA | NA | NA | 615 | NA | NA | NA | NA | NA | |
| <500m | NA | NA | NA | NA | 60.3 | NA | NA | NA | NA | NA | 60.3 |
| >500m<700m | NA | NA | NA | NA | 39.4 | NA | NA | NA | NA | NA | 39.4 |
| >700m<1000m | NA | NA | NA | NA | 0.3 | NA | NA | NA | NA | NA | 0.3 |
| >1000m<1500m | NA | NA | NA | NA | 0.0 | NA | NA | NA | NA | NA | 0.0 |
| >1500m<2000m | NA | NA | NA | NA | 0.0 | NA | NA | NA | NA | NA | 0.0 |
| >2000m | NA | NA | NA | NA | 0.0 | NA | NA | NA | NA | NA | 0.0 |
| % fixes outside estate | NA | NA | NA | NA | 0.0 | NA | NA | NA | NA | NA | 0.0 |

### Site F2

| Pre-shooting | Time spent (number of fixes) (%) | | | | | | | | | | Mean ± SE |
|---|---|---|---|---|---|---|---|---|---|---|---|
| | 57182 | 57183 | 57184 | 57185 | 57186 | 57187 | 57188 | 57189 | 57190 | 57191 | |
| Total fixes | 175 | 306 | 92 | 1886 | 1096 | 1869 | 2008 | 180 | 1822 | 732 | |
| <500m | 100 | 100 | 100 | 98.8 | 99.8 | 99.9 | 75.5 | 100 | 99.6 | 99.9 | 97.4 ± 2.4 |
| >500m<700m | 0.0 | 0.0 | 0.0 | 1.0 | 0.2 | 0.1 | 9.9 | 0.0 | 0.4 | 0.1 | 1.2 ± 1.0 |
| >700m<1000m | 0.0 | 0.0 | 0.0 | 0.2 | 0.0 | 0.0 | 9.7 | 0.0 | 0 | 0.0 | 1.0 ± 1.0 |
| >1000m<1500m | 0.0 | 0.0 | 0.0 | 0.0 | 0.0 | 0.0 | 4.9 | 0.0 | 0.1 | 0.0 | 0.5 ± 0.5 |
| >1500m<2000m | 0.0 | 0.0 | 0.0 | 0.0 | 0.0 | 0.0 | 0.0 | 0.0 | 0.0 | 0.0 | 0.0 |
| >2000m | 0.0 | 0.0 | 0.0 | 0.0 | 0.0 | 0.0 | 0.0 | 0.0 | 0.0 | 0.0 | 0.0 |
| % fixes outside estate | 0.0 | 0.0 | 0.0 | 0.0 | 0.0 | 0.0 | 0.0 | 0.0 | 0.0 | 0.0 | 0.0 |

| Shooting | Time spent (number of fixes) (%) | | | | | | | | | | Mean ± SE |
|---|---|---|---|---|---|---|---|---|---|---|---|
| | 57182 | 57183 | 57184 | 57185 | 57186 | 57187 | 57188 | 57189 | 57190 | 57191 | |
| Total fixes | NA | NA | NA | 1163 | NA | 1533 | 1943 | NA | 1683 | NA | |
| <500m | NA | NA | NA | 97.1 | NA | 96.5 | 94.9 | NA | 98.8 | NA | 96.8 ± 0.8 |
| >500m<700m | NA | NA | NA | 2.7 | NA | 2.2 | 1.4 | NA | 1.0 | NA | 1.8 ± 0.4 |
| >700m<1000m | NA | NA | NA | 0.1 | NA | 0.3 | 2.0 | NA | 0.2 | NA | 0.7 ± 0.5 |
| >1000m<1500m | NA | NA | NA | 0.2 | NA | 1.0 | 1.7 | NA | 0.0 | NA | 0.7 ± 0.4 |
| >1500m<2000m | NA | NA | NA | 0.0 | NA | 0.0 | 0.0 | NA | 0.0 | NA | 0.0 |
| >2000m | NA | NA | NA | 0.0 | NA | 0.0 | 0.0 | NA | 0.0 | NA | 0.0 |
| % fixes outside estate | NA | NA | NA | 0.0 | NA | 0.0 | 0.0 | NA | 0.0 | NA | 0.0 |

| Post-shooting | Time spent (number of fixes) (%) | | | | | | | | | | Mean ± SE |
|---|---|---|---|---|---|---|---|---|---|---|---|
| | 57182 | 57183 | 57184 | 57185 | 57186 | 57187 | 57188 | 57189 | 57190 | 57191 | |
| Total fixes | NA | NA | NA | NA | NA | NA | 154 | NA | NA | NA | |
| <500m | NA | NA | NA | NA | NA | NA | 100 | NA | NA | NA | 100 |
| >500m<700m | NA | NA | NA | NA | NA | NA | 0.0 | NA | NA | NA | 0.0 |
| >700m<1000m | NA | NA | NA | NA | NA | NA | 0.0 | NA | NA | NA | 0.0 |
| >1000m<1500m | NA | NA | NA | NA | NA | NA | 0.0 | NA | NA | NA | 0.0 |
| >1500m<2000m | NA | NA | NA | NA | NA | NA | 0.0 | NA | NA | NA | 0.0 |
| >2000m | NA | NA | NA | NA | NA | NA | 0.0 | NA | NA | NA | 0.0 |
| % fixes outside estate | NA | NA | NA | NA | NA | NA | 0.0 | NA | NA | NA | 0.0 |

### Site G

| Pre-shooting | Time spent (number of fixes) (%) | | | | | | | | | | | | Mean ± SE |
|---|---|---|---|---|---|---|---|---|---|---|---|---|---|
| | 56561 | 56562 | 57212 | 57213 | 57214 | 57215 | 57216 | 57217 | 57218 | 57219 | 57220 | 57221 | |
| Total fixes | 232 | 994 | 972 | 1267 | 1051 | 1317 | 1145 | 1171 | 1270 | 1230 | 1186 | 1265 | |
| <500m | 100 | 98.9 | 100 | 58.0 | 73.8 | 100 | 100 | 100 | 82.0 | 99.8 | 100 | 55.2 | 89.0 ± 5.0 |
| >500m<700m | 0.0 | 0.5 | 0.0 | 11.1 | 19.7 | 0.0 | 0.0 | 0.0 | 14.6 | 0.2 | 0.0 | 10.0 | 4.7 ± 2.1 |
| >700m<1000m | 0.0 | 0.6 | 0.0 | 27.6 | 6.5 | 0.0 | 0.0 | 0.0 | 3.4 | 0.0 | 0.0 | 17.6 | 4.6 ± 2.6 |
| >1000m<1500m | 0.0 | 0.0 | 0.0 | 3.2 | 0.0 | 0.0 | 0.0 | 0.0 | 0.1 | 0.0 | 0.0 | 10.1 | 1.1 ± 0.9 |
| >1500m<2000m | 0.0 | 0.0 | 0.0 | 0.0 | 0.0 | 0.0 | 0.0 | 0.0 | 0.0 | 0.0 | 0.0 | 3.9 | 0.3 ± 0.3 |
| >2000m | 0.0 | 0.0 | 0.0 | 0.0 | 0.0 | 0.0 | 0.0 | 0.0 | 0.0 | 0.0 | 0.0 | 3.2 | 0.3 ± 0.3 |
| % fixes outside estate | 0.0 | 0.0 | 0.0 | 0.0 | 0.0 | 0.0 | 0.0 | 0.0 | 0.0 | 0.0 | 0.0 | 3.1 | 0.3 ± 0.3 |

| Shooting | Time spent (number of fixes) (%) | | | | | | | | | | | | Mean ± SE |
|---|---|---|---|---|---|---|---|---|---|---|---|---|---|
| | 56561 | 56562 | 57212 | 57213 | 57214 | 57215 | 57216 | 57217 | 57218 | 57219 | 57220 | 57221 | |
| Total fixes | NA | 575 | 1652 | 1467 | 2200 | 2498 | 605 | 2268 | 2292 | 2312 | 1982 | 2437 | |
| <500m | NA | 99.5 | 97.2 | 5.0 | 67.5 | 62.9 | 100 | 98.9 | 81.5 | 94.8 | 52.6 | 0.0 | 69.1 ± 11.1 |
| >500m<700m | NA | 0.5 | 2.4 | 22.9 | 29.4 | 31.8 | 0.0 | 0.7 | 13.0 | 3.0 | 19.5 | 0.0 | 11.2 ± 3.8 |
| >700m<1000m | NA | 0.0 | 0.5 | 70.4 | 2.8 | 4.8 | 0.0 | 0.5 | 5.3 | 2.1 | 21.6 | 0.0 | 9.8 ± 6.3 |
| >1000m<1500m | NA | 0.0 | 0.0 | 1.7 | 0.2 | 0.4 | 0.0 | 0 | 0.3 | 0.1 | 3.7 | 0.0 | 0.6 ± 0.3 |
| >1500m<2000m | NA | 0.0 | 0.0 | 0.0 | 0.0 | 0.0 | 0.0 | 0.0 | 0.0 | 0.0 | 1.5 | 1.3 | 0.3 ± 0.2 |
| >2000m | NA | 0.0 | 0.0 | 0.0 | 0.0 | 0.0 | 0.0 | 0.0 | 0.0 | 0.0 | 1.1 | 98.7 | 9.1 ± 9.0 |
| ≥50% fixes >500m | - | - | - | Yes | - | - | - | - | - | - | - | Yes | - |
| ≥50% fixes >1000m | - | - | - | - | - | - | - | - | - | - | - | Yes | - |
| % fixes outside estate | NA | 0.0 | 0.1 | 25.8 | 0.2 | 0.0 | 0.0 | 0.0 | 0.0 | 0.0 | 4.2 | 98.8 | 11.7 ± 9.0 |

| Post-shooting | Time spent (number of fixes) (%) | | | | | | | | | | | | Mean ± SE |
|---|---|---|---|---|---|---|---|---|---|---|---|---|---|
| | 56561 | 56562 | 57212 | 57213 | 57214 | 57215 | 57216 | 57217 | 57218 | 57219 | 57220 | 57221 | |
| Total fixes | NA | NA | NA | NA | 944 | 731 | NA | 718 | 1009 | 681 | NA | 394 | |
| <500m | NA | NA | NA | NA | 100 | 81.7 | NA | 100 | 11.2 | 96 | NA | 0.0 | 64.8 ± 19.0 |
| >500m<700m | NA | NA | NA | NA | 0.0 | 7.9 | NA | 0.0 | 53.0 | 2.8 | NA | 0.0 | 10.6 ± 8.6 |
| >700m<1000m | NA | NA | NA | NA | 0.0 | 10.4 | NA | 0.0 | 35.2 | 1.2 | NA | 0.0 | 7.8 ± 5.7 |
| >1000m<1500m | NA | NA | NA | NA | 0.0 | 0.0 | NA | 0.0 | 0.5 | 0.0 | NA | 0.0 | 0.1 ± 0.1 |
| >1500m<2000m | NA | NA | NA | NA | 0.0 | 0.0 | NA | 0.0 | 0.0 | 0.0 | NA | 40.6 | 6.8 ± 6.8 |
| >2000m | NA | NA | NA | NA | 0.0 | 0.0 | NA | 0.0 | 0.1 | 0.0 | NA | 59.4 | 9.9 ± 9.9 |
| ≥50% fixes >500m | - | - | - | - | - | - | - | - | Yes | - | - | Yes | - |
| ≥50% fixes >1000m | - | - | - | - | - | - | - | - | - | - | - | Yes | - |
| % fixes outside estate | NA | NA | NA | NA | 0.0 | 0.0 | NA | 0.0 | 0.1 | 0.0 | NA | 58.9 | 9.8 ± 9.8 |

### Site H

| Pre-shooting | Time spent (number of fixes) (%) | | | | | | | | | | | Mean ± SE |
|---|---|---|---|---|---|---|---|---|---|---|---|---|
| | 56565 | 57129 | 57192 | 57196 | 57198 | 57199 | 57200 | 57201 | 57363 | 57364 | 57365 | |
| Total fixes | 231 | 999 | 1265 | 1221 | 542 | 606 | 1298 | 1254 | 1211 | 1117 | 1171 | |
| <500m | 100 | 100 | 99.4 | 100 | 98.3 | 99.7 | 97.8 | 97.8 | 98.8 | 99.9 | 99.5 | 99.2 ± 0.3 |
| >500m<700m | 0.0 | 0.0 | 0.6 | 0.0 | 1.5 | 0.3 | 1.8 | 0.7 | 0.8 | 0.1 | 0.5 | 0.6 ± 0.2 |
| >700m<1000m | 0.0 | 0.0 | 0.1 | 0.0 | 0.2 | 0 | 0.2 | 1.1 | 0.4 | 0.0 | 0.0 | 0.2 ± 0.1 |
| >1000m<1500m | 0.0 | 0.0 | 0.0 | 0.0 | 0.0 | 0.0 | 0.1 | 0.4 | 0.0 | 0.0 | 0.0 | 0.05 ± 0.04 |
| >1500m<2000m | 0.0 | 0.0 | 0.0 | 0.0 | 0.0 | 0.0 | 0.0 | 0.0 | 0.0 | 0.0 | 0.0 | 0.0 |
| >2000m | 0.0 | 0.0 | 0.0 | 0.0 | 0.0 | 0.0 | 0.0 | 0.0 | 0.0 | 0.0 | 0.0 | 0.0 |
| % fixes outside estate | 0.0 | 0.0 | 0.0 | 0.0 | 0.0 | 0.0 | 0.0 | 0.0 | 0.0 | 0.0 | 0.0 | 0.0 |

| Shooting | Time spent (number of fixes) (%) | | | | | | | | | | | Mean ± SE |
|---|---|---|---|---|---|---|---|---|---|---|---|---|
| | 56565 | 57129 | 57192 | 57196 | 57198 | 57199 | 57200 | 57201 | 57363 | 57364 | 57365 | |
| Total fixes | NA | 273 | 1416 | 2334 | NA | NA | 2297 | 2340 | 2310 | 2110 | 245 | |
| <500m | NA | 99.6 | 87.3 | 87.8 | NA | NA | 100 | 100 | 89.2 | 92.6 | 99.6 | 94.5 ± 2.1 |
| >500m<700m | NA | 0.4 | 11.7 | 2.1 | NA | NA | 0.0 | 0.0 | 3.6 | 4.4 | 0.4 | 2.8 ± 1.4 |
| >700m<1000m | NA | 0.0 | 0.8 | 9.9 | NA | NA | 0.0 | 0.0 | 6.3 | 2.7 | 0.0 | 2.5 ± 1.3 |
| >1000m<1500m | NA | 0.0 | 0.2 | 0.3 | NA | NA | 0.0 | 0.0 | 0.8 | 0.4 | 0.0 | 0.2 ± 0.1 |
| >1500m<2000m | NA | 0.0 | 0.0 | 0.0 | NA | NA | 0.0 | 0.0 | 0.0 | 0.0 | 0.0 | 0.0 |
| >2000m | NA | 0.0 | 0.0 | 0.0 | NA | NA | 0.0 | 0.0 | 0.0 | 0.0 | 0.0 | 0.0 |
| % fixes outside estate | NA | 0.0 | 0.0 | 0.0 | NA | NA | 0.0 | 0.0 | 0.2 | 0.0 | 0.0 | 0.03 ± 0.03 |

| Post-shooting | Time spent (number of fixes) (%) | | | | | | | | | | | Mean ± SE |
|---|---|---|---|---|---|---|---|---|---|---|---|---|
| | 56565 | 57129 | 57192 | 57196 | 57198 | 57199 | 57200 | 57201 | 57363 | 57364 | 57365 | |
| Total fixes | NA | NA | NA | NA | NA | NA | 890 | 1243 | 1249 | NA | NA | |
| <500m | NA | NA | NA | NA | NA | NA | 96.1 | 100 | 99.8 | NA | NA | 98.6 ± 1.3 |
| >500m<700m | NA | NA | NA | NA | NA | NA | 0.8 | 0.0 | 0.2 | NA | NA | 0.3 ± 0.2 |
| >700m<1000m | NA | NA | NA | NA | NA | NA | 2.8 | 0.0 | 0.0 | NA | NA | 0.9 ± 0.9 |
| >1000m<1500m | NA | NA | NA | NA | NA | NA | 0.3 | 0.0 | 0.0 | NA | NA | 0.1 ± 0.1 |
| >1500m<2000m | NA | NA | NA | NA | NA | NA | 0.0 | 0.0 | 0.0 | NA | NA | 0.0 |
| >2000m | NA | NA | NA | NA | NA | NA | 0.0 | 0.0 | 0.0 | NA | NA | 0.0 |
| % fixes outside estate | NA | NA | NA | NA | NA | NA | 0.0 | 0.0 | 0.0 | NA | NA | 0.0 |

### Site I

| Pre-shooting | Time spent (number of fixes) (%) | | | | | | Mean ± SE |
|---|---|---|---|---|---|---|---|
| | 56565 | 57137 | 57164 | 57171 | 57178 | 57209 | |
| Total fixes | 37 | 1182 | 79 | 237 | 426 | 226 | |
| <500m | 100 | 63.5 | 100 | 92.4 | 100 | 100 | 92.7 ± 6.0 |
| >500m<700m | 0.0 | 21.7 | 0.0 | 7.6 | 0.0 | 0.0 | 4.9 ± 3.6 |
| >700m<1000m | 0.0 | 9.3 | 0.0 | 0.0 | 0.0 | 0.0 | 1.6 ± 1.6 |
| >1000m<1500m | 0.0 | 5.6 | 0.0 | 0.0 | 0.0 | 0.0 | 0.9 ± 0.9 |
| >1500m<2000m | 0.0 | 0.0 | 0.0 | 0.0 | 0.0 | 0.0 | 0.0 |
| >2000m | 0.0 | 0.0 | 0.0 | 0.0 | 0.0 | 0.0 | 0.0 |
| % fixes outside estate | 0.0 | 0.0 | 0.0 | 0.0 | 0.0 | 0.0 | 0.0 |

| Shooting | Time spent (number of fixes) (%) | | | | | | Mean ± SE |
|---|---|---|---|---|---|---|---|
| | 56565 | 57137 | 57164 | 57171 | 57178 | 57209 | |
| Total fixes | NA | 1844 | NA | NA | NA | NA | |
| <500m | NA | 1.0 | NA | NA | NA | NA | 1.0 |
| >500m<700m | NA | 2.4 | NA | NA | NA | NA | 2.4 |
| >700m<1000m | NA | 15.9 | NA | NA | NA | NA | 15.9 |
| >1000m<1500m | NA | 78.4 | NA | NA | NA | NA | 78.4 |
| >1500m<2000m | NA | 2.3 | NA | NA | NA | NA | 2.3 |
| >2000m | NA | 0.0 | NA | NA | NA | NA | 0.0 |
| ≥50% fixes >500m | - | Yes | - | - | - | - | - |
| ≥50% fixes >1000m | - | Yes | - | - | - | - | - |
| % fixes outside estate | NA | 0.0 | NA | NA | NA | NA | 0.0 |

| Post-shooting | Time spent (number of fixes) (%) | | | | | | Mean ± SE |
|---|---|---|---|---|---|---|---|
| | 56565 | 57137 | 57164 | 57171 | 57178 | 57209 | |
| Total fixes | NA | 1173 | NA | NA | NA | NA | |
| <500m | NA | 7.0 | NA | NA | NA | NA | 7.0 |
| >500m<700m | NA | 17.3 | NA | NA | NA | NA | 17.3 |
| >700m<1000m | NA | 37.1 | NA | NA | NA | NA | 37.1 |
| >1000m<1500m | NA | 37.3 | NA | NA | NA | NA | 37.3 |
| >1500m<2000m | NA | 1.4 | NA | NA | NA | NA | 1.4 |
| >2000m | NA | 0.0 | NA | NA | NA | NA | 0.0 |
| ≥50% fixes >500m | - | Yes | - | - | - | - | - |
| % fixes outside estate | NA | 0.0 | NA | NA | NA | NA | 0.0 |

**Section 3: Direction of dispersal for tagged birds. Directional preference tables show the distribution of the percentage of total GPS fixes relative to compass bearing (45° octants) centred on the release pen, for individual birds and the mean across all tagged birds). Only points lying outside of the release pen are included in directionality results. Post-shooting results are presented only for sites with more than one bird alive during this phase.**

#### Site A

| Pre-shooting | | Bearing sector (% total fixes per bearing) | | | | | | | |
|---|---|---|---|---|---|---|---|---|---|
| **Bird** | **Tot. Fixes** | **0-45** | **45-90** | **90-135** | **135-180** | **180-225** | **225-270** | **270-315** | **315-360** |
| **57142** | 862 | 9.5 | 6.4 | 5.3 | 6.8 | 10.7 | 10.7 | 42.2 | 8.4 |
| **57143** | 1494 | 52.3 | 11.6 | 5.0 | 2.6 | 1.8 | 2.4 | 4.8 | 19.5 |
| **57144** | 1328 | 52.6 | 20.8 | 3.7 | 3.7 | 2.6 | 2.0 | 2.6 | 12.0 |
| **57145** | 1520 | 21.7 | 58.0 | 6.0 | 2.0 | 1.3 | 2.2 | 2.2 | 6.5 |
| **57146** | 1314 | 7.0 | 9.5 | 6.1 | 14.7 | 44.7 | 8.4 | 5.5 | 4.0 |
| **57147** | 1489 | 73.5 | 8.8 | 3.7 | 2.6 | 1.2 | 1.7 | 1.6 | 7.0 |
| **57148** | 1517 | 66.7 | 12.7 | 6.3 | 3.8 | 1.2 | 1.6 | 1.6 | 6.1 |
| **57149** | 1512 | 44.0 | 39.3 | 5.7 | 2.8 | 1.5 | 1.1 | 1.8 | 3.8 |
| **57150** | 1481 | 70.9 | 11.2 | 3.2 | 1.8 | 0.7 | 1.8 | 1.9 | 8.6 |
| **57151** | 1370 | 12.3 | 5.3 | 2.8 | 5.8 | 10.9 | 36.4 | 17.1 | 9.3 |
| **Mean** | 1389 | 41.1 | 18.3 | 4.8 | 4.6 | 7.7 | 6.8 | 8.1 | 8.5 |

| Shooting | | Bearing sector (% total fixes per bearing) | | | | | | | |
|---|---|---|---|---|---|---|---|---|---|
| **Bird** | **Tot. Fixes** | **0-45** | **45-90** | **90-135** | **135-180** | **180-225** | **225-270** | **270-315** | **315-360** |
| **57142** | NA | NA | NA | NA | NA | NA | NA | NA | NA |
| **57143** | 2407 | 19.6 | 77.1 | 0.2 | 0.1 | 0.2 | 0.1 | 1.2 | 1.6 |
| **57144** | 2349 | 13.4 | 76.7 | 1.8 | 1.0 | 5.8 | 1.1 | 0.2 | 0.1 |
| **57145** | 2495 | 5.2 | 88.3 | 1.6 | 1.3 | 3.2 | 0.2 | 0.2 | 0.1 |
| **57146** | 520 | 0.0 | 1.3 | 0.2 | 7.1 | 89.8 | 1.3 | 0.2 | 0.0 |
| **57147** | NA | NA | NA | NA | NA | NA | NA | NA | NA |
| **57148** | 640 | 76.9 | 20.6 | 0.6 | 0.0 | 0.0 | 0.0 | 0.0 | 1.9 |
| **57149** | 982 | 21.8 | 77.3 | 0.6 | 0.1 | 0.0 | 0.0 | 0.0 | 0.2 |
| **57150** | 635 | 97.8 | 0.0 | 0.2 | 0.0 | 0.0 | 0.0 | 0.0 | 2.0 |
| **57151** | 1021 | 0.0 | 0.0 | 0.0 | 0.8 | 9.6 | 75.4 | 14.2 | 0.0 |
| **Mean** | 1381 | 29.3 | 42.7 | 0.6 | 1.3 | 13.6 | 9.8 | 2.0 | 0.7 |

| Post-shooting | | Bearing sector (% total fixes per bearing) | | | | | | | |
|---|---|---|---|---|---|---|---|---|---|
| **Bird** | **Tot. Fixes** | **0-45** | **45-90** | **90-135** | **135-180** | **180-225** | **225-270** | **270-315** | **315-360** |
| **57142** | NA | NA | NA | NA | NA | NA | NA | NA | NA |
| **57143** | 847 | 0.0 | 100 | 0.0 | 0.0 | 0.0 | 0.0 | 0.0 | 0.0 |
| **57144** | 871 | 2.0 | 98.0 | 0.0 | 0.0 | 0.0 | 0.0 | 0.0 | 0.0 |
| **57145** | 1241 | 0.0 | 99.9 | 0.1 | 0.0 | 0.0 | 0.0 | 0.0 | 0.0 |
| **57146** | NA | NA | NA | NA | NA | NA | NA | NA | NA |
| **57147** | NA | NA | NA | NA | NA | NA | NA | NA | NA |
| **57148** | NA | NA | NA | NA | NA | NA | NA | NA | NA |
| **57149** | NA | NA | NA | NA | NA | NA | NA | NA | NA |
| **57150** | NA | NA | NA | NA | NA | NA | NA | NA | NA |
| **57151** | NA | NA | NA | NA | NA | NA | NA | NA | NA |
| **Mean** | 986 | 29.3 | 42.7 | 0.6 | 1.3 | 13.6 | 9.8 | 2.0 | 0.7 |

### Site B

| Pre-shooting | | Bearing sector (% total fixes per bearing) | | | | | | | |
|---|---|---|---|---|---|---|---|---|---|
| **Bird** | **Tot. Fixes** | **0-45** | **45-90** | **90-135** | **135-180** | **180-225** | **225-270** | **270-315** | **315-360** |
| **57122** | 805 | 0.0 | 0.2 | 5.3 | 33.7 | 54.3 | 5.7 | 0.5 | 0.2 |
| **57123** | 1049 | 3.1 | 8.6 | 16.2 | 33.6 | 20.8 | 5.8 | 8.5 | 3.4 |
| **57124** | 934 | 1.7 | 16.6 | 17.0 | 28.8 | 11.9 | 6.4 | 7.3 | 10.3 |
| **57125** | 1126 | 3.0 | 2.9 | 1.6 | 4.2 | 5.3 | 40.3 | 36.1 | 6.5 |
| **57126** | 987 | 10.1 | 5.3 | 10.6 | 19.8 | 19.0 | 18.6 | 5.7 | 10.8 |
| **57127** | 1068 | 9.2 | 24.6 | 10.0 | 21.8 | 18.1 | 5.1 | 3.0 | 8.2 |
| **57128** | 1280 | 1.7 | 1.6 | 1.5 | 5.9 | 8.8 | 32.2 | 36.7 | 11.6 |
| **57129** | 59 | 22.0 | 30.5 | 15.3 | 13.6 | 10.2 | 3.4 | 1.7 | 3.4 |
| **57130** | 902 | 0.8 | 1.1 | 3.1 | 13.1 | 38.6 | 38.5 | 2.0 | 2.9 |
| **57131** | 1269 | 0.5 | 30.3 | 11.6 | 7.9 | 9.7 | 25.6 | 12.1 | 2.3 |
| **Mean** | 948 | 5.2 | 12.2 | 9.2 | 18.2 | 19.7 | 18.2 | 11.4 | 6.0 |

| Shooting | | Bearing sector (% total fixes per bearing) | | | | | | | |
|---|---|---|---|---|---|---|---|---|---|
| **Bird** | **Tot. Fixes** | **0-45** | **45-90** | **90-135** | **135-180** | **180-225** | **225-270** | **270-315** | **315-360** |
| **57122** | 906 | 0.1 | 0.4 | 0.8 | 14.7 | 42.6 | 38.3 | 3.0 | 0.1 |
| **57123** | 2250 | 0.1 | 0.4 | 1.0 | 3.0 | 1.2 | 1.6 | 88.7 | 4.0 |
| **57124** | 2090 | 0.5 | 57.0 | 30.1 | 10.7 | 0.9 | 0.4 | 0.1 | 0.3 |
| **57125** | 827 | 0.4 | 0.1 | 0.4 | 0.7 | 0.5 | 63.4 | 33.7 | 0.8 |
| **57126** | 2261 | 54.3 | 14.9 | 2.0 | 0.3 | 0.3 | 18.5 | 7.0 | 2.7 |
| **57127** | 2330 | 83.7 | 16.3 | 0.0 | 0.0 | 0.0 | 0.0 | 0.0 | 0.0 |
| **57128** | 2274 | 0.5 | 3.6 | 0.8 | 0.8 | 2.2 | 69.5 | 21.0 | 1.6 |
| **57129** | NA | NA | NA | NA | NA | NA | NA | NA | NA |
| **57130** | 537 | 0.0 | 0.2 | 0.0 | 0.2 | 18.3 | 81.4 | 0.0 | 0.0 |
| **57131** | 2428 | 0.0 | 0.0 | 0.0 | 0.0 | 0.0 | 100.0 | 0.0 | 0.0 |
| **Mean** | 1767.0 | 15.5 | 10.3 | 3.9 | 3.4 | 7.3 | 41.4 | 17.1 | 1.1 |

| Post-shooting | | Bearing sector (% total fixes per bearing) | | | | | | | |
|---|---|---|---|---|---|---|---|---|---|
| **Bird** | **Tot. Fixes** | **0-45** | **45-90** | **90-135** | **135-180** | **180-225** | **225-270** | **270-315** | **315-360** |
| **57122** | NA | NA | NA | NA | NA | NA | NA | NA | NA |
| **57123** | 1204 | 0.0 | 0.0 | 0.0 | 0.0 | 0.4 | 5.1 | 94.5 | 0.0 |
| **57124** | 383 | 0.0 | 1.6 | 84.3 | 13.8 | 0.0 | 0.0 | 0.3 | 0.0 |
| **57125** | NA | NA | NA | NA | NA | NA | NA | NA | NA |
| **57126** | 92 | 35.9 | 64.1 | 0.0 | 0.0 | 0.0 | 0.0 | 0.0 | 0.0 |
| **57127** | 853 | 99.9 | 0.1 | 0.0 | 0.0 | 0.0 | 0.0 | 0.0 | 0.0 |
| **57128** | NA | NA | NA | NA | NA | NA | NA | NA | NA |
| **57129** | NA | NA | NA | NA | NA | NA | NA | NA | NA |
| **57130** | NA | NA | NA | NA | NA | NA | NA | NA | NA |
| **57131** | 400 | 0.0 | 0.0 | 0.0 | 0.0 | 0.0 | 100.0 | 0.0 | 0.0 |
| **Mean** | 586.4 | 27.2 | 13.2 | 16.9 | 2.8 | 0.1 | 21.0 | 19.0 | 0.0 |

### Site C1

| Pre-shooting | | Bearing sector (% total fixes per bearing) | | | | | | | |
|---|---|---|---|---|---|---|---|---|---|
| **Bird** | **Tot. Fixes** | **0-45** | **45-90** | **90-135** | **135-180** | **180-225** | **225-270** | **270-315** | **315-360** |
| **57152** | 1383 | 0.1 | 1.5 | 50.6 | 29.3 | 5.2 | 11.1 | 1.6 | 0.6 |
| **57153** | 1813 | 0.5 | 1.5 | 20.9 | 2.2 | 1.5 | 69.7 | 3.7 | 0.0 |
| **57154** | 36 | 0.0 | 2.8 | 11.1 | 27.8 | 30.6 | 25.0 | 2.8 | 0.0 |
| **57155** | 49 | 0.0 | 0.0 | 16.3 | 0.0 | 53.1 | 30.6 | 0.0 | 0.0 |
| **57156** | 346 | 0.0 | 0.6 | 5.5 | 13.0 | 18.8 | 52.6 | 9.0 | 0.6 |
| **57157** | 16 | 0.0 | 0.0 | 56.3 | 12.5 | 0.0 | 0.0 | 31.3 | 0.0 |
| **57158** | 156 | 0.6 | 1.9 | 27.6 | 14.1 | 18.6 | 10.3 | 26.3 | 0.6 |
| **57159** | 162 | 1.9 | 2.5 | 32.7 | 14.2 | 9.3 | 24.1 | 15.4 | 0.0 |
| **57160** | 2019 | 16.3 | 1.4 | 14.0 | 3.3 | 4.5 | 33.5 | 11.5 | 15.5 |
| **57161** | 1778 | 1.7 | 1.1 | 10.0 | 1.7 | 1.8 | 2.6 | 66.9 | 14.1 |
| **Mean** | 775.8 | 2.1 | 1.3 | 24.5 | 11.8 | 14.3 | 25.9 | 16.8 | 3.1 |

| Shooting | | Bearing sector (% total fixes per bearing) | | | | | | | |
|---|---|---|---|---|---|---|---|---|---|
| **Bird** | **Tot. Fixes** | **0-45** | **45-90** | **90-135** | **135-180** | **180-225** | **225-270** | **270-315** | **315-360** |
| **57152** | 1717 | 0.1 | 0.2 | 0.2 | 0.2 | 0.5 | 95.8 | 2.9 | 0.1 |
| **57153** | 1938 | 0.0 | 0.0 | 0.0 | 0.0 | 0.0 | 100.0 | 0.0 | 0.0 |
| **57154** | NA | NA | NA | NA | NA | NA | NA | NA | NA |
| **57155** | NA | NA | NA | NA | NA | NA | NA | NA | NA |
| **57156** | NA | NA | NA | NA | NA | NA | NA | NA | NA |
| **57157** | NA | NA | NA | NA | NA | NA | NA | NA | NA |
| **57158** | NA | NA | NA | NA | NA | NA | NA | NA | NA |
| **57159** | NA | NA | NA | NA | NA | NA | NA | NA | NA |
| **57160** | 1372 | 82.1 | 2.1 | 0.1 | 0.0 | 0.0 | 0.0 | 0.1 | 15.6 |
| **57161** | 1600 | 9.6 | 6.0 | 0.3 | 0.3 | 1.3 | 5.8 | 67.4 | 9.5 |
| **Mean** | 1656.75 | 23.0 | 2.1 | 0.1 | 0.1 | 0.4 | 50.4 | 17.6 | 6.3 |

| Post-shooting | | Bearing sector (% total fixes per bearing) | | | | | | | |
|---|---|---|---|---|---|---|---|---|---|
| **Bird** | **Tot. Fixes** | **0-45** | **45-90** | **90-135** | **135-180** | **180-225** | **225-270** | **270-315** | **315-360** |
| **57152** | 574 | 0.0 | 0.0 | 0.0 | 0.0 | 0.0 | 94.6 | 5.4 | 0.0 |
| **57153** | 1192 | 0.0 | 0.0 | 0.0 | 0.0 | 0.0 | 100.0 | 0.0 | 0.0 |
| **57154** | NA | NA | NA | NA | NA | NA | NA | NA | NA |
| **57155** | NA | NA | NA | NA | NA | NA | NA | NA | NA |
| **57156** | NA | NA | NA | NA | NA | NA | NA | NA | NA |
| **57157** | NA | NA | NA | NA | NA | NA | NA | NA | NA |
| **57158** | NA | NA | NA | NA | NA | NA | NA | NA | NA |
| **57159** | NA | NA | NA | NA | NA | NA | NA | NA | NA |
| **57160** | NA | NA | NA | NA | NA | NA | NA | NA | NA |
| **57161** | NA | NA | NA | NA | NA | NA | NA | NA | NA |
| **Mean** | 883 | 0.0 | 0.0 | 0.0 | 0.0 | 0.0 | 97.3 | 2.7 | 0.0 |

### Site C2

| Pre-shooting | | Bearing sector (% total fixes per bearing) | | | | | | | |
|---|---|---|---|---|---|---|---|---|---|
| **Bird** | **Tot. Fixes** | **0-45** | **45-90** | **90-135** | **135-180** | **180-225** | **225-270** | **270-315** | **315-360** |
| **57162** | 1732 | 1.2 | 2.2 | 37.2 | 54.6 | 0.3 | 1.2 | 1.0 | 2.2 |
| **57163** | 1595 | 2.1 | 45.4 | 45.2 | 4.4 | 1.0 | 0.3 | 0.3 | 1.4 |
| **57164** | 21 | 0.0 | 33.3 | 42.9 | 14.3 | 4.8 | 4.8 | 0.0 | 0.0 |
| **57165a** | 158 | 3.2 | 17.1 | 68.4 | 7.0 | 0.6 | 1.3 | 0.0 | 2.5 |
| **57166** | 1281 | 4.5 | 24.5 | 45.7 | 5.8 | 7.8 | 3.4 | 2.8 | 5.5 |
| **57167** | 1188 | 15.0 | 2.3 | 20.5 | 5.6 | 18.3 | 14.0 | 18.1 | 6.4 |
| **57169** | 1243 | 32.8 | 21.5 | 20.0 | 3.5 | 4.9 | 3.3 | 2.1 | 11.9 |
| **57170** | 1608 | 1.0 | 58.9 | 37.8 | 0.7 | 0.2 | 0.6 | 0.3 | 0.5 |
| **57171** | 124 | 29.0 | 20.2 | 6.5 | 0.8 | 2.4 | 23.4 | 4.0 | 13.7 |
| **57137*** | 26 | 3.8 | 0.0 | 3.8 | 0.0 | 80.8 | 7.7 | 0.0 | 3.8 |
| **57165b*** | 1236 | 2.2 | 0.3 | 0.9 | 0.1 | 0.6 | 4.0 | 62.4 | 29.5 |
| **Mean** | 928.4 | 8.6 | 20.5 | 29.9 | 8.8 | 11.1 | 5.8 | 8.3 | 7.0 |

| Shooting | | Bearing sector (% total fixes per bearing) | | | | | | | |
|---|---|---|---|---|---|---|---|---|---|
| **Bird** | **Tot. Fixes** | **0-45** | **45-90** | **90-135** | **135-180** | **180-225** | **225-270** | **270-315** | **315-360** |
| **57162** | 13 | 0.0 | 0.0 | 15.4 | 84.6 | 0.0 | 0.0 | 0.0 | 0.0 |
| **57163** | 17 | 0.0 | 100.0 | 0.0 | 0.0 | 0.0 | 0.0 | 0.0 | 0.0 |
| **57164** | NA | NA | NA | NA | NA | NA | NA | NA | NA |
| **57165a** | NA | NA | NA | NA | NA | NA | NA | NA | NA |
| **57166** | 850 | 13.1 | 13.9 | 29.2 | 5.6 | 20.7 | 3.3 | 9.6 | 4.6 |
| **57167** | 1942 | 1.7 | 69.4 | 24.2 | 4.2 | 0.4 | 0.0 | 0.0 | 0.2 |
| **57169** | 273 | 44.3 | 4.8 | 0.0 | 0.4 | 1.8 | 1.1 | 2.9 | 44.7 |
| **57170** | 1252 | 2.2 | 97.8 | 0.0 | 0.0 | 0.0 | 0.0 | 0.0 | 0.0 |
| **57171** | NA | NA | NA | NA | NA | NA | NA | NA | NA |
| **57137*** | NA | NA | NA | NA | NA | NA | NA | NA | NA |
| **57165b*** | NA | NA | NA | NA | NA | NA | NA | NA | NA |
| **Mean** | 724.5 | 10.2 | 47.6 | 11.5 | 15.8 | 3.8 | 0.7 | 2.1 | 8.2 |

### Site D

| Pre-shooting | | Bearing sector (% total fixes per bearing) | | | | | | | |
|---|---|---|---|---|---|---|---|---|---|
| **Bird** | **Tot. Fixes** | **0-45** | **45-90** | **90-135** | **135-180** | **180-225** | **225-270** | **270-315** | **315-360** |
| **57132** | 1078 | 25.2 | 0.6 | 4.6 | 26.8 | 10.0 | 9.7 | 0.7 | 22.3 |
| **57133** | 1097 | 0.1 | 1.1 | 7.8 | 24.2 | 60.7 | 3.8 | 1.8 | 0.5 |
| **57134** | 521 | 0.2 | 5.6 | 10.4 | 36.1 | 11.9 | 9.6 | 23.4 | 2.9 |
| **57135** | 1092 | 0.0 | 0.3 | 3.2 | 9.4 | 2.0 | 83.3 | 1.6 | 0.1 |
| **57136** | 1073 | 0.5 | 2.8 | 3.1 | 14.6 | 8.4 | 12.3 | 41.6 | 16.8 |
| **57137** | 67 | 0.0 | 0.0 | 34.3 | 28.4 | 25.4 | 10.4 | 1.5 | 0.0 |
| **57138** | 1334 | 18.7 | 1.6 | 1.5 | 1.3 | 2.4 | 69.8 | 2.3 | 2.3 |
| **57139** | 981 | 17.4 | 34.1 | 20.2 | 21.4 | 1.3 | 1.2 | 1.5 | 2.8 |
| **57140** | 1147 | 62.1 | 33.8 | 1.5 | 1.1 | 0.5 | 0.3 | 0.3 | 0.3 |
| **57141** | 878 | 0.7 | 2.2 | 23.6 | 38.4 | 8.2 | 8.0 | 3.2 | 15.8 |
| **Mean** | 724.5 | 12.5 | 8.2 | 11.0 | 20.2 | 13.1 | 20.9 | 7.8 | 6.4 |

| Shooting | | Bearing sector (% total fixes per bearing) | | | | | | | |
|---|---|---|---|---|---|---|---|---|---|
| **Bird** | **Tot. Fixes** | **0-45** | **45-90** | **90-135** | **135-180** | **180-225** | **225-270** | **270-315** | **315-360** |
| **57132** | 794 | 41.9 | 0.8 | 0.0 | 0.4 | 0.0 | 0.1 | 5.4 | 51.4 |
| **57133** | 443 | 0.0 | 0.0 | 0.0 | 1.8 | 97.7 | 0.2 | 0.2 | 0.0 |
| **57134** | NA | NA | NA | NA | NA | NA | NA | NA | NA |
| **57135** | 2314 | 0.0 | 0.0 | 0.0 | 0.0 | 0.0 | 99.9 | 0.1 | 0.0 |
| **57136** | 2137 | 3.5 | 0.05 | 0.05 | 0.3 | 0.1 | 2.8 | 20.9 | 72.3 |
| **57137** | NA | NA | NA | NA | NA | NA | NA | NA | NA |
| **57138** | 844 | 14.1 | 0.1 | 1.5 | 50.8 | 2.6 | 13.6 | 1.2 | 16.0 |
| **57139** | 1908 | 1.9 | 96.8 | 1.0 | 0.2 | 0.2 | 0.1 | 0.0 | 0.0 |
| **57140** | 2290 | 0.1 | 99.9 | 0.0 | 0.0 | 0.0 | 0.0 | 0.0 | 0.0 |
| **57141** | NA | NA | NA | NA | NA | NA | NA | NA | NA |
| **Mean** | 1532.9 | 8.8 | 28.2 | 0.4 | 7.6 | 14.4 | 16.7 | 4.0 | 20.0 |

| Post-shooting | | Bearing sector (% total fixes per bearing) | | | | | | | |
|---|---|---|---|---|---|---|---|---|---|
| **Bird** | **Tot. Fixes** | **0-45** | **45-90** | **90-135** | **135-180** | **180-225** | **225-270** | **270-315** | **315-360** |
| **57132** | NA | NA | NA | NA | NA | NA | NA | NA | NA |
| **57133** | NA | NA | NA | NA | NA | NA | NA | NA | NA |
| **57134** | NA | NA | NA | NA | NA | NA | NA | NA | NA |
| **57135** | 1238 | 0.0 | 0.0 | 0.0 | 0.0 | 0.0 | 100.0 | 0.0 | 0.0 |
| **57136** | 565 | 1.6 | 0.0 | 0.0 | 0.0 | 0.0 | 0.0 | 0.0 | 98.4 |
| **57137** | NA | NA | NA | NA | NA | NA | NA | NA | NA |
| **57138** | NA | NA | NA | NA | NA | NA | NA | NA | NA |
| **57139** | NA | NA | NA | NA | NA | NA | NA | NA | NA |
| **57140** | 1210 | 0.0 | 100.0 | 0.0 | 0.0 | 0.0 | 0.0 | 0.0 | 0.0 |
| **57141** | NA | NA | NA | NA | NA | NA | NA | NA | NA |
| **Mean** | 1004.3 | 0.5 | 33.3 | 0.0 | 0.0 | 0.0 | 33.3 | 0.0 | 32.8 |

### Site E

| Pre-shooting | | Bearing sector (% total fixes per bearing) | | | | | | | |
|---|---|---|---|---|---|---|---|---|---|
| **Bird** | **Tot. Fixes** | **0-45** | **45-90** | **90-135** | **135-180** | **180-225** | **225-270** | **270-315** | **315-360** |
| **57202** | 326 | 72.7 | 3.4 | 1.8 | 7.7 | 2.1 | 0.3 | 2.8 | 9.2 |
| **57203** | 206 | 44.7 | 18.9 | 11.2 | 12.1 | 3.4 | 1.9 | 0.0 | 7.8 |
| **57204** | 265 | 11.7 | 2.3 | 4.5 | 19.6 | 38.9 | 12.1 | 7.2 | 3.8 |
| **57205** | 159 | 1.9 | 3.8 | 5.7 | 34.0 | 47.2 | 4.4 | 2.5 | 0.6 |
| **57206** | 300 | 13.0 | 10.0 | 59.0 | 9.3 | 2.7 | 1.3 | 0.0 | 4.7 |
| **57207** | 133 | 22.6 | 12.0 | 3.0 | 29.3 | 12.8 | 2.3 | 8.3 | 9.8 |
| **57208** | 203 | 4.9 | 2.5 | 11.8 | 36.9 | 30.0 | 3.0 | 3.4 | 7.4 |
| **57209** | 99 | 1.0 | 21.2 | 7.1 | 22.2 | 47.5 | 1.0 | 0.0 | 0.0 |
| **57210** | 391 | 19.2 | 9.5 | 27.1 | 9.2 | 14.1 | 9.7 | 5.4 | 5.9 |
| **57211** | 200 | 41.0 | 13.5 | 5.5 | 9.5 | 9.0 | 1.0 | 1.0 | 19.5 |
| **Mean** | 228.2 | 23.3 | 9.7 | 13.7 | 19.0 | 20.8 | 3.7 | 3.1 | 6.9 |

| Shooting | | Bearing sector (% total fixes per bearing) | | | | | | | |
|---|---|---|---|---|---|---|---|---|---|
| **Bird** | **Tot. Fixes** | **0-45** | **45-90** | **90-135** | **135-180** | **180-225** | **225-270** | **270-315** | **315-360** |
| **57202** | 178 | 84.3 | 10.1 | 0.6 | 0.0 | 0.0 | 0.0 | 0.0 | 5.1 |
| **57203** | 1845 | 12.8 | 6.8 | 39.7 | 29.8 | 2.1 | 0.4 | 0.2 | 8.1 |
| **57204** | 2311 | 0.2 | 0.1 | 0.4 | 34.6 | 21.2 | 42.5 | 0.4 | 0.5 |
| **57205** | 1639 | 2.9 | 91.3 | 3.5 | 1.3 | 0.7 | 0.0 | 0.0 | 0.2 |
| **57206** | 553 | 28.0 | 4.7 | 25.0 | 6.5 | 8.3 | 10.3 | 0.9 | 16.3 |
| **57207** | NA | NA | NA | NA | NA | NA | NA | NA | NA |
| **57208** | 401 | 10.7 | 9.0 | 54.4 | 9.0 | 1.5 | 1.2 | 1.7 | 12.5 |
| **57209** | NA | NA | NA | NA | NA | NA | NA | NA | NA |
| **57210** | 2175 | 0.0 | 5.1 | 63.0 | 13.6 | 18.3 | 0.0 | 0.0 | 0.0 |
| **57211** | 2228 | 7.8 | 0.9 | 0.0 | 0.5 | 0.1 | 10.1 | 71.8 | 8.8 |
| **Mean** | 1416.3 | 18.3 | 16.0 | 23.3 | 11.9 | 6.5 | 8.1 | 9.4 | 6.4 |

### Site F1

| Pre-shooting | | Bearing sector (% total fixes per bearing) | | | | | | | |
|---|---|---|---|---|---|---|---|---|---|
| **Bird** | **Tot. Fixes** | **0-45** | **45-90** | **90-135** | **135-180** | **180-225** | **225-270** | **270-315** | **315-360** |
| **57172** | 1696 | 51.2 | 15.8 | 8.7 | 1.2 | 1.1 | 2.1 | 6.5 | 13.3 |
| **57173** | 1672 | 66.9 | 8.7 | 0.3 | 0.9 | 1.6 | 2.7 | 6.8 | 12.1 |
| **57174** | 170 | 69.4 | 0.0 | 0.0 | 0.0 | 1.2 | 0.6 | 2.4 | 26.5 |
| **57175** | 1730 | 62.7 | 5.8 | 2.4 | 0.1 | 0.9 | 2.8 | 8.8 | 16.5 |
| **57176** | 1754 | 25.5 | 10.3 | 27.5 | 12.1 | 2.5 | 5.9 | 8.7 | 7.6 |
| **57177** | 217 | 54.8 | 0.5 | 0.5 | 0.0 | 0.0 | 0.0 | 1.8 | 42.4 |
| **57178** | 102 | 71.6 | 0.0 | 0.0 | 0.0 | 1.0 | 2.9 | 1.0 | 23.5 |
| **57179** | 1658 | 67.7 | 2.3 | 0.1 | 0.1 | 0.7 | 2.7 | 8.7 | 17.9 |
| **57180** | 119 | 56.3 | 0.8 | 0.0 | 0.0 | 0.0 | 0.8 | 1.7 | 40.3 |
| **57181** | 5 | 40.0 | 0.0 | 20.0 | 0.0 | 0.0 | 0.0 | 0.0 | 40.0 |
| **Mean** | 912.3 | 56.6 | 4.4 | 5.9 | 1.4 | 0.9 | 2.1 | 4.6 | 24.0 |

| Shooting | | Bearing sector (% total fixes per bearing) | | | | | | | |
|---|---|---|---|---|---|---|---|---|---|
| **Bird** | **Tot. Fixes** | **0-45** | **45-90** | **90-135** | **135-180** | **180-225** | **225-270** | **270-315** | **315-360** |
| **57172** | 1595 | 71.6 | 20.8 | 5.8 | 0.1 | 0.3 | 0.2 | 0.4 | 0.8 |
| **57173** | 1018 | 93.8 | 2.7 | 0.1 | 0.0 | 0.1 | 0.0 | 0.0 | 3.3 |
| **57174** | NA | NA | NA | NA | NA | NA | NA | NA | NA |
| **57175** | 449 | 85.7 | 0.7 | 0.2 | 0.0 | 0.0 | 0.0 | 5.6 | 7.8 |
| **57176** | 1705 | 0.8 | 2.3 | 24.3 | 1.5 | 0.1 | 15.9 | 55.0 | 0.2 |
| **57177** | NA | NA | NA | NA | NA | NA | NA | NA | NA |
| **57178** | NA | NA | NA | NA | NA | NA | NA | NA | NA |
| **57179** | 23 | 82.6 | 0.0 | 0.0 | 0.0 | 0.0 | 0.0 | 0.0 | 17.4 |
| **57180** | NA | NA | NA | NA | NA | NA | NA | NA | NA |
| **57181** | NA | NA | NA | NA | NA | NA | NA | NA | NA |
| **Mean** | 958.0 | 66.9 | 5.3 | 6.1 | 0.3 | 0.1 | 3.2 | 12.2 | 5.9 |

### Site F2

| Pre-shooting | | Bearing sector (% total fixes per bearing) | | | | | | | |
|---|---|---|---|---|---|---|---|---|---|
| **Bird** | **Tot. Fixes** | **0-45** | **45-90** | **90-135** | **135-180** | **180-225** | **225-270** | **270-315** | **315-360** |
| **57182** | 50 | 14.0 | 2.0 | 0.0 | 0.0 | 0.0 | 6.0 | 28.0 | 50.0 |
| **57183** | 80 | 17.5 | 0.0 | 0.0 | 0.0 | 0.0 | 1.3 | 11.3 | 70.0 |
| **57184** | 18 | 16.7 | 0.0 | 0.0 | 0.0 | 0.0 | 5.6 | 16.7 | 61.1 |
| **57185** | 1476 | 7.0 | 2.6 | 1.2 | 2.0 | 22.8 | 45.8 | 9.6 | 9.0 |
| **57186** | 676 | 2.5 | 13.8 | 0.7 | 0.7 | 9.3 | 34.2 | 23.1 | 15.7 |
| **57187** | 1420 | 3.2 | 3.0 | 4.0 | 24.6 | 23.5 | 17.5 | 10.4 | 13.9 |
| **57188** | 1674 | 6.0 | 85.8 | 0.4 | 0.1 | 0.1 | 0.4 | 2.4 | 4.8 |
| **57189** | 71 | 7.0 | 4.2 | 1.4 | 0.0 | 1.4 | 0.0 | 36.6 | 49.3 |
| **57190** | 1216 | 5.6 | 0.3 | 0.0 | 0.8 | 27.7 | 42.2 | 14.5 | 8.9 |
| **57191** | 338 | 10.7 | 4.7 | 0.6 | 0.3 | 0.3 | 11.5 | 35.2 | 36.7 |
| **Mean** | 701.9 | 9.0 | 11.6 | 0.8 | 2.9 | 8.5 | 16.4 | 18.8 | 31.9 |

| Shooting | | Bearing sector (% total fixes per bearing) | | | | | | | |
|---|---|---|---|---|---|---|---|---|---|
| **Bird** | **Tot. Fixes** | **0-45** | **45-90** | **90-135** | **135-180** | **180-225** | **225-270** | **270-315** | **315-360** |
| **57182** | NA | NA | NA | NA | NA | NA | NA | NA | NA |
| **57183** | NA | NA | NA | NA | NA | NA | NA | NA | NA |
| **57184** | NA | NA | NA | NA | NA | NA | NA | NA | NA |
| **57185** | 1163 | 0.0 | 0.0 | 0.0 | 2.7 | 67.3 | 29.6 | 0.4 | 0.0 |
| **57186** | NA | NA | NA | NA | NA | NA | NA | NA | NA |
| **57187** | 1531 | 0.1 | 1.9 | 4.2 | 24.1 | 54.7 | 15.0 | 0.1 | 0.0 |
| **57188** | 1943 | 12.9 | 87.0 | 0.1 | 0.0 | 0.0 | 0.0 | 0.0 | 0.0 |
| **57189** | NA | NA | NA | NA | NA | NA | NA | NA | NA |
| **57190** | 1678 | 0.1 | 0.0 | 0.4 | 0.4 | 59.9 | 38.4 | 0.8 | 0.0 |
| **57191** | NA | NA | NA | NA | NA | NA | NA | NA | NA |
| **Mean** | 1578.8 | 3.3 | 22.2 | 1.2 | 6.8 | 45.5 | 20.7 | 0.3 | 0.0 |

### Site G

| Pre-shooting | | Bearing sector (% total fixes per bearing) | | | | | | | |
|---|---|---|---|---|---|---|---|---|---|
| **Bird** | **Tot. Fixes** | **0-45** | **45-90** | **90-135** | **135-180** | **180-225** | **225-270** | **270-315** | **315-360** |
| **56561** | 112 | 1.8 | 1.8 | 0.0 | 0.9 | 3.6 | 40.2 | 43.8 | 8.0 |
| **56562** | 605 | 14.7 | 27.8 | 6.9 | 13.6 | 18.2 | 5.8 | 8.9 | 4.1 |
| **57212** | 726 | 29.1 | 5.2 | 0.1 | 0.0 | 0.0 | 1.8 | 28.4 | 35.4 |
| **57213** | 843 | 0.4 | 0.0 | 1.2 | 3.6 | 22.8 | 47.0 | 24.8 | 0.4 |
| **57214** | 655 | 2.7 | 0.6 | 0.5 | 30.1 | 38.5 | 9.9 | 11.0 | 6.7 |
| **57215** | 923 | 1.2 | 0.3 | 0.0 | 0.5 | 5.7 | 34.8 | 50.6 | 6.8 |
| **57216** | 815 | 8.1 | 1.2 | 0.2 | 0.1 | 1.5 | 13.5 | 35.6 | 39.8 |
| **57217** | 580 | 9.8 | 1.0 | 0.5 | 6.6 | 7.8 | 10.0 | 32.1 | 32.2 |
| **57218** | 892 | 0.9 | 10.5 | 3.4 | 41.0 | 28.6 | 5.6 | 7.8 | 2.1 |
| **57219** | 528 | 2.1 | 2.3 | 0.8 | 14.4 | 49.2 | 17.4 | 12.7 | 1.1 |
| **57220** | 494 | 0.2 | 0.2 | 0.2 | 4.9 | 43.1 | 30.4 | 18.2 | 2.8 |
| **57221** | 916 | 1.6 | 0.9 | 0.5 | 18.9 | 45.2 | 21.0 | 8.7 | 3.2 |
| **Mean** | 674.1 | 6.1 | 4.3 | 1.2 | 11.2 | 22.0 | 19.8 | 23.5 | 11.9 |

| Shooting | | Bearing sector (% total fixes per bearing) | | | | | | | |
|---|---|---|---|---|---|---|---|---|---|
| **Bird** | **Tot. Fixes** | **0-45** | **45-90** | **90-135** | **135-180** | **180-225** | **225-270** | **270-315** | **315-360** |
| **56561** | NA | NA | NA | NA | NA | NA | NA | NA | NA |
| **56562** | 569 | 10.4 | 77.3 | 3.3 | 1.8 | 1.4 | 0.5 | 1.4 | 3.9 |
| **57212** | 1641 | 21.4 | 20.8 | 40.0 | 9.9 | 2.6 | 0.7 | 1.2 | 3.4 |
| **57213** | 1467 | 0.0 | 0.0 | 0.0 | 0.0 | 7.5 | 13.0 | 79.0 | 0.5 |
| **57214** | 2033 | 46.1 | 9.4 | 3.9 | 32.0 | 5.6 | 0.4 | 0.6 | 1.9 |
| **57215** | 2496 | 0.0 | 0.04 | 0.0 | 1.9 | 24.7 | 65.0 | 7.7 | 0.6 |
| **57216** | 526 | 4.8 | 0.2 | 21.9 | 25.7 | 3.0 | 1.1 | 12.0 | 31.4 |
| **57217** | 1447 | 30.4 | 15.8 | 0.7 | 3.2 | 2.6 | 2.1 | 13.4 | 31.9 |
| **57218** | 1959 | 0.8 | 0.0 | 0.6 | 57.9 | 38.7 | 1.4 | 0.5 | 0.1 |
| **57219** | 1659 | 2.9 | 1.9 | 2.7 | 17.7 | 29.4 | 26.3 | 16.8 | 2.4 |
| **57220** | 1901 | 1.7 | 0.2 | 2.4 | 34.6 | 40.2 | 14.4 | 4.2 | 2.3 |
| **57221** | 2437 | 0.0 | 0.0 | 0.0 | 0.0 | 1.1 | 98.9 | 0.0 | 0.0 |
| **Mean** | 1648.6 | 10.8 | 11.4 | 6.9 | 16.8 | 14.2 | 20.4 | 12.4 | 7.1 |

| Post-shooting | | Bearing sector (% total fixes per bearing) | | | | | | | |
|---|---|---|---|---|---|---|---|---|---|
| **Bird** | **Tot. Fixes** | **0-45** | **45-90** | **90-135** | **135-180** | **180-225** | **225-270** | **270-315** | **315-360** |
| **56561** | NA | NA | NA | NA | NA | NA | NA | NA | NA |
| **56562** | NA | NA | NA | NA | NA | NA | NA | NA | NA |
| **57212** | NA | NA | NA | NA | NA | NA | NA | NA | NA |
| **57213** | NA | NA | NA | NA | NA | NA | NA | NA | NA |
| **57214** | 479 | 37.2 | 1.3 | 4.8 | 10.9 | 19.8 | 14.4 | 8.4 | 3.3 |
| **57215** | 718 | 0.1 | 0.0 | 0.0 | 2.5 | 27.6 | 30.9 | 38.9 | 0.0 |
| **57216** | NA | NA | NA | NA | NA | NA | NA | NA | NA |
| **57217** | 627 | 42.1 | 1.3 | 0.0 | 1.6 | 2.4 | 1.3 | 7.5 | 43.9 |
| **57218** | 1009 | 0.0 | 0.0 | 4.3 | 63.7 | 32.0 | 0.0 | 0.0 | 0.0 |
| **57219** | 432 | 0.2 | 0.9 | 0.0 | 14.4 | 30.8 | 43.1 | 10.4 | 0.2 |
| **57220** | NA | NA | NA | NA | NA | NA | NA | NA | NA |
| **57221** | 394 | 0.0 | 0.0 | 0.0 | 0.0 | 0.8 | 99.2 | 0.0 | 0.0 |
| **Mean** | 609.8 | 13.3 | 0.6 | 1.5 | 15.5 | 18.9 | 31.5 | 10.9 | 7.9 |

### Site H

| Pre-shooting | | Bearing sector (% total fixes per bearing) | | | | | | | |
|---|---|---|---|---|---|---|---|---|---|
| **Bird** | **Tot. Fixes** | **0-45** | **45-90** | **90-135** | **135-180** | **180-225** | **225-270** | **270-315** | **315-360** |
| **56565** | 128 | 1.6 | 0.8 | 0.0 | 2.3 | 2.3 | 1.6 | 22.7 | 68.8 |
| **57129** | 142 | 21.8 | 7.7 | 8.5 | 5.6 | 9.9 | 10.6 | 15.5 | 20.4 |
| **57192** | 592 | 4.2 | 25.2 | 67.4 | 1.7 | 0.2 | 0.0 | 0.8 | 0.5 |
| **57196** | 295 | 12.2 | 26.8 | 2.4 | 0.7 | 2.7 | 0.3 | 40.3 | 14.6 |
| **57198** | 203 | 10.8 | 5.4 | 3.9 | 3.9 | 2.5 | 29.6 | 26.1 | 17.7 |
| **57199** | 304 | 22.7 | 27.6 | 6.9 | 0.0 | 1.3 | 27.3 | 5.6 | 8.6 |
| **57200** | 605 | 15.0 | 36.2 | 9.8 | 2.0 | 0.7 | 15.0 | 16.5 | 4.8 |
| **57201** | 294 | 16.3 | 42.2 | 19.7 | 5.1 | 0.3 | 0.3 | 1.0 | 15.0 |
| **57363** | 331 | 6.0 | 1.5 | 0.6 | 3.9 | 3.6 | 4.8 | 55.6 | 23.9 |
| **57364** | 215 | 12.6 | 3.7 | 0.9 | 5.1 | 4.7 | 18.1 | 32.1 | 22.8 |
| **57365** | 538 | 0.9 | 0.6 | 0.9 | 2.8 | 6.9 | 31.4 | 45.7 | 10.8 |
| **Mean** | 331.5 | 11.3 | 16.2 | 11.0 | 3.0 | 3.2 | 12.6 | 23.8 | 18.9 |

| Shooting | | Bearing sector (% total fixes per bearing) | | | | | | | |
|---|---|---|---|---|---|---|---|---|---|
| **Bird** | **Tot. Fixes** | **0-45** | **45-90** | **90-135** | **135-180** | **180-225** | **225-270** | **270-315** | **315-360** |
| **56565** | NA | NA | NA | NA | NA | NA | NA | NA | NA |
| **57129** | 137 | 0.7 | 0.0 | 0.7 | 0.7 | 0.0 | 37.2 | 54.0 | 6.6 |
| **57192** | 1409 | 0.1 | 35.4 | 61.8 | 2.5 | 0.0 | 0.0 | 0.1 | 0.1 |
| **57196** | 1134 | 4.2 | 1.3 | 3.1 | 7.9 | 4.9 | 6.0 | 55.4 | 17.1 |
| **57198** | NA | NA | NA | NA | NA | NA | NA | NA | NA |
| **57199** | NA | NA | NA | NA | NA | NA | NA | NA | NA |
| **57200** | 1089 | 0.9 | 1.7 | 2.9 | 4.2 | 7.2 | 14.9 | 62.9 | 5.3 |
| **57201** | 2245 | 0.1 | 9.6 | 88.0 | 2.0 | 0.2 | 0.0 | 0.0 | 0.05 |
| **57363** | 1177 | 2.0 | 3.0 | 1.2 | 5.2 | 31.9 | 4.7 | 33.4 | 18.6 |
| **57364** | 2054 | 0.0 | 0.0 | 0.1 | 0.1 | 0.9 | 87.5 | 10.3 | 1.0 |
| **57365** | 152 | 0.7 | 0.7 | 2.0 | 32.2 | 10.5 | 14.5 | 35.5 | 3.9 |
| **Mean** | 1174.6 | 1.1 | 6.5 | 20.0 | 6.9 | 7.0 | 20.6 | 31.5 | 6.6 |

| Post-shooting | | Bearing sector (% total fixes per bearing) | | | | | | | |
|---|---|---|---|---|---|---|---|---|---|
| **Bird** | **Tot. Fixes** | **0-45** | **45-90** | **90-135** | **135-180** | **180-225** | **225-270** | **270-315** | **315-360** |
| **56565** | NA | NA | NA | NA | NA | NA | NA | NA | NA |
| **57129** | NA | NA | NA | NA | NA | NA | NA | NA | NA |
| **57192** | NA | NA | NA | NA | NA | NA | NA | NA | NA |
| **57196** | NA | NA | NA | NA | NA | NA | NA | NA | NA |
| **57198** | NA | NA | NA | NA | NA | NA | NA | NA | NA |
| **57199** | NA | NA | NA | NA | NA | NA | NA | NA | NA |
| **57200** | 445 | 0.0 | 0.4 | 1.1 | 0.9 | 7.2 | 56.0 | 29.2 | 5.2 |
| **57201** | 1243 | 0.0 | 4.2 | 95.8 | 0.0 | 0.0 | 0.0 | 0.0 | 0.0 |
| **57363** | 727 | 1.7 | 1.0 | 0.1 | 0.6 | 5.5 | 30.1 | 7.6 | 53.5 |
| **57364** | NA | NA | NA | NA | NA | NA | NA | NA | NA |
| **57365** | NA | NA | NA | NA | NA | NA | NA | NA | NA |
| **Mean** | 805.0 | 0.6 | 1.9 | 32.4 | 0.5 | 4.2 | 28.7 | 12.3 | 19.6 |

### Site I

| Pre-shooting | | Bearing sector (% total fixes per bearing) | | | | | | | |
|---|---|---|---|---|---|---|---|---|---|
| **Bird** | **Tot. Fixes** | **0-45** | **45-90** | **90-135** | **135-180** | **180-225** | **225-270** | **270-315** | **315-360** |
| **56565** | 22 | 4.5 | 31.8 | 40.9 | 4.5 | 13.6 | 0.0 | 0.0 | 4.5 |
| **57137** | 975 | 7.7 | 2.9 | 2.3 | 0.4 | 1.0 | 22.9 | 58.4 | 4.5 |
| **57164** | 63 | 79.4 | 3.2 | 12.7 | 3.2 | 0.0 | 0.0 | 1.6 | 0.0 |
| **57171** | 101 | 31.7 | 14.9 | 19.8 | 5.0 | 3.0 | 5.9 | 8.9 | 10.9 |
| **57178** | 177 | 11.3 | 18.6 | 22.0 | 3.4 | 5.1 | 4.0 | 21.5 | 14.1 |
| **57209** | 88 | 5.7 | 29.5 | 28.4 | 5.7 | 5.7 | 2.3 | 9.1 | 13.6 |
| **Mean** | 237.7 | 23.4 | 16.8 | 21.0 | 3.7 | 4.7 | 5.8 | 16.6 | 8.0 |

| Shooting | | Bearing sector (% total fixes per bearing) | | | | | | | |
|---|---|---|---|---|---|---|---|---|---|
| **Bird** | **Tot. Fixes** | **0-45** | **45-90** | **90-135** | **135-180** | **180-225** | **225-270** | **270-315** | **315-360** |
| **56565** | NA | NA | NA | NA | NA | NA | NA | NA | NA |
| **57137** | 1844 | 0.0 | 0.0 | 0.0 | 0.0 | 0.0 | 0.0 | 95.0 | 5.0 |
| **57164** | NA | NA | NA | NA | NA | NA | NA | NA | NA |
| **57171** | NA | NA | NA | NA | NA | NA | NA | NA | NA |
| **57178** | NA | NA | NA | NA | NA | NA | NA | NA | NA |
| **57209** | NA | NA | NA | NA | NA | NA | NA | NA | NA |
| **Mean** | 1844 | 0.0 | 0.0 | 0.0 | 0.0 | 0.0 | 0.0 | 95.0 | 5.0 |
